# Transgenerational pathogen effects: Maternal pathogen exposure reduces offspring fitness

**DOI:** 10.1101/2023.03.14.532659

**Authors:** Kristina M. McIntire, Marcin K. Dziuba, Elizabeth B. Haywood, Miles L. Robertson, Megan Vaandrager, Emma Baird, Fiona E. Corcoran, Taleah Nelson, Michael H. Cortez, Meghan A. Duffy

## Abstract

Pathogens can alter the phenotype not only of exposed hosts, but also of future generations. Transgenerational immune priming, where parental infection drives reduced susceptibility of offspring, has been particularly well explored, but pathogens can also alter life history traits of offspring. Here, we examined the potential for transgenerational impacts of a microsporidian pathogen, *Ordospora pajunii*, by experimentally measuring the impact of maternal exposure on offspring fitness in the presence and absence of parasites, and then developing mathematical models that explored the population-level impacts of these transgenerational effects. We did not find evidence of transgenerational immune priming: offspring of exposed mothers became infected at high rates, similar to offspring of unexposed mothers, and the infection burden did not differ between these two groups. We also did not find any evidence of transgenerational tolerance, where daughters of exposed mothers have higher fitness after infection. Instead, we found evidence for negative transgenerational impacts of infection: uninfected offspring of exposed mothers had substantially greater early life mortality than uninfected offspring of unexposed mothers. Offspring of exposed mothers also had reduced growth rate, fewer clutches, and fewer offspring. We propose that these observations should be considered transgenerational virulence, where a pathogen reduces the fitness of the offspring of infected hosts. Our parameterized mathematical model allowed us to explore the impacts of transgenerational virulence at the population level. If transgenerational virulence manifests as decreased reproduction or increased mortality in offspring, as we saw in the empirical portion of our study, this reduces total host density, infection prevalence, and infected host density, which could have implications for both host conservation and spillover risk. We propose that transgenerational virulence might be common and is a concept worthy of further empirical and theoretical exploration.

## Introduction

Transgenerational effects occur when a parental phenotype affects an offspring’s phenotype (Ben-Ami et al., 2020), and can be induced by environmental stressors. Examples of such transgenerational effects include maternal predation exposure altering offspring anti-predation behavior and growth (Cattelan et al., 2020; McGhee et al., 2012; Storm and Lima, 2010; Coslovsky and Richner, 2011) and food restriction impacting offspring size and survival (Reznick et al., 1996; Jann and Ward, 1999; Garbutt and Little, 2017; Coakley et al., 2018). Pathogens have been shown to induce transgenerational effects in a wide range of taxa (e.g., Ben-Ami et al. 2020; Rasekh et al. 2022; Sadd and Schmid-Hempel 2009; Lim et al. 2021), especially in the form of altered immune responses in the offspring of exposed hosts (Moret, 2006; Sadd et al., 2005; Lim et al., 2021). In particular, hosts that are exposed to a pathogen can reduce the susceptibility of their offspring to that pathogen or different pathogens, a phenomenon known as transgenerational immune priming (TGIP; Ben-Ami et al., 2020; Little et al., 2003; Rasekh et al., 2022; Moret, 2006; Sadd et al., 2005; Rutkowski et al., 2023). TGIP has been demonstrated in a range of host systems, such as in bumblebees and beetles, where maternal exposure to bacteria leads to greater antimicrobial activity in offspring (Sadd et al., 2005; Moret, 2006) and in beetles and pipefish where paternal exposure has been found to elicit the potential for greater antimicrobial response (Beemelmanns and Roth, 2016; Eggert et al., 2014); similarly, in *Daphnia*, maternal bacterial challenge reduces offspring susceptibility by as much as 50% (Ben-Ami et al., 2020).

In addition to altering immune responses of offspring, maternal pathogen exposure can also impact offspring life history traits such as body size, growth rate, reproduction, and longevity (Prior et al., 2011; Paraskevopoulou et al., 2022). The majority of studies looking for transgenerational effects of pathogens on offspring life history traits have focused on the potential for transgenerational tolerance – that is, increased fitness of offspring who are also exposed to pathogens. For example, offspring of fungus-exposed *Daphnia magna* mothers increased reproduction when exposed to the fungus (Paraskevopoulou et al., 2022); similarly, in *Drosophila*, immune-challenged offspring of immune-challenged mothers increased reproduction compared to those with control mothers (Nystrand and Dowling, 2014). Thus, it is clear both that impacts of exposure to pathogens can carry across generations and that these impacts could involve both resistance and tolerance mechanisms. While maternal exposure to a pathogen might drive increased resistance or tolerance of offspring, that might come at a cost. Across taxa, immune responses are often costly (French et al., 2007; Hasselquist and Nilsson, 2012; Jehan et al., 2022). These costs are often due to resource constraints, and can manifest as trade-offs with life-history traits (such as growth rate and reproduction; French et al., 2007; Bashir-Tanoli and Tinsley, 2014; Urlacher et al., 2018; Garcia et al., 2020; Jehan et al., 2022). Immune priming – both within generations and transgenerationally – has been shown to have costs, including impacts on the quantity and/or quality of offspring (Schulz et al., 2023). For example, in mosquitoes, prior *Plasmodium* challenge increases resistance upon subsequent exposure, but also reduces reproductive success (Contreras-Garduño et al., 2014). Similarly, TGIP reduces offspring quantity, growth, or extends development time in beetles (Dhinaut et al., 2018; Prakash et al., 2022; Schulz et al., 2019), and reduces offspring reproduction in hornworms and crickets (Trauer and Hilker, 2013; McNamara et al., 2014). In total, pathogen effects can span generations, with the potential for both positive and negative impacts on offspring fitness.

Somewhat surprisingly, despite an extensive body of theoretical studies exploring the impacts of pathogens on hosts, the effect of transgenerational impacts of pathogens on disease dynamics has largely been overlooked. Incorporating such transgenerational impacts seems likely to alter predicted dynamics, given that even small changes in host resistance and tolerance levels may have a large impact (Duffy and Sivars-Becker, 2007; Wilber et al., 2017); for example, in amphibian-chytrid infections, resistance and tolerance interact to predict host population persistence (Wilber et al., 2017). Models that have been extended to include immune priming within an individual demonstrate that such priming has important implications for host-pathogen disease dynamics (Tidbury et al., 2012); notably, immune priming has the potential to increase pathogen persistence and to destabilize host and pathogen populations (Tidbury et al., 2012). Therefore, changes in host resistance and tolerance due to the transgenerational effects of pathogens, including via TGIP or transgenerational tolerance, seem likely to be important for host-pathogen interactions, but this has not been rigorously explored. In addition to impacting the host and pathogen populations themselves, transgenerational effects of pathogens might also impact other aspects of infection dynamics, including the potential for pathogen spillover (i.e., transmission of pathogens to uninfected host populations or species; Wells and Clark, 2019).

Here, we explore the potential for TGIP and transgenerational disease tolerance, as well as the impact of transgenerational effects of pathogens on predicted patterns of disease, combining empirical and theoretical studies. First, we used the freshwater crustacean, *Daphnia dentifera*, to examine the impact of maternal exposure to a microsporidian pathogen, *Ordospora pajunii*, on offspring susceptibility and life history traits. We did not find evidence of TGIP or transgenerational tolerance. Instead, we found strongly reduced fitness in the offspring of microsporidian-exposed mothers. An additional comparison helped us contextualize these transgenerational fitness impacts of maternal exposure to the microsporidian: transgenerational impacts of maternal *O. pajunii* - exposure on early mortality of uninfected offspring was greater than the within generation impact of infection by a fungal pathogen, *Metschnikowia bicuspidata*, that is considered highly virulent due to its effects on mortality. Based on these findings, we then developed mathematical models that incorporate transgenerational impacts of pathogen exposure, using these to explore how the transgenerational effects we observed in the empirical portion of this study might impact population and infection dynamics.

## Empirical Methods

### Host-Pathogen System

We conducted laboratory experiments with the “S” genotype of *Daphnia dentifera* (also referred to as the “Standard” or “Std” genotype in other studies on this system), a North American freshwater microcrustacean as host. In addition to being ecologically important, *Daphnia* species and their pathogens are model systems for understanding host-pathogen interactions (Ebert, 2005). *Daphnia* are cyclically parthenogenetic, and we maintained the S genotype under standard conditions that maintained clonal reproduction throughout this study. We used the BDWalsh isolate (isolated from Walsh Lake, Washtenaw Co. Michigan USA) of *Ordospora pajunii* (Dziuba et al., 2024*b*; de Albuquerque et al., 2022) as our focal pathogen. This microsporidian commonly infects *D. dentifera* gut epithelial cells after ingestion during routine filter feeding, and is generally considered to be a pathogen of low virulence (Dziuba et al., 2024*b*). In a recent study of six lake populations, five had epidemics of *O. pajunii* in *D. dentifera*, with peak infection prevalences of 18-37% (Davenport et al., 2024).

### Laboratory Methods

#### Overview of experimental treatments

We were interested in the potential for transgenerational immune priming, as well as in characterizing the fitness impacts on hosts of maternal exposure and within generation exposure (referred to as ‘current’ exposure hereafter). To study this, we had ten ‘maternal’ (F0) generation mothers who were exposed to *O. pajunii* and ten who were not exposed (Figure 1). For the mothers who were not exposed, we used a split brood design, where we collected three offspring per mother, and allocated one to the control treatment (‘None/None’ animals in Figure 1), one to the *O. pajunii*-exposure treatment (‘None/Op’ treatment), and one to the *M. bicuspidata*-exposure treatment (‘None/Mb’ treatment). We also intended to use a split brood design for the *O. pajunii* maternal exposure treatment, collecting two animals per mother and exposing one to *O. pajunii* (‘Op/Op’ animals in Figure 1) and one to the control treatment (‘Op/None’). However, two of the maternal *O. pajunii*-exposure treatment animals only produced a single offspring. Therefore, there were six mothers from this treatment that contributed two offspring (which were split between the two current exposure treatments), two that contributed a single offspring, and two that contributed three offspring (which were split between the two current exposure treatments). Overall, this yielded 50 total ‘current exposure’ treatment animals (the F1 generation; Figure 1): 40 that were part of the four exposure treatments in the main experiment, and 10 ‘additional comparison’ animals (that is, the ‘None/Mb’ animals) that allowed us to compare transgenerational impacts of *O. pajunii* exposure with the within generation impacts of the fungus *Metschnikowia bicuspidata*. Sample size in this work was limited to ten F1 individuals per experimental treatment (Figure 1), due to the practical limitations of synchronizing daughter production from maternal (F0) individuals and the available quantities of infection doses.

**Figure 1:**
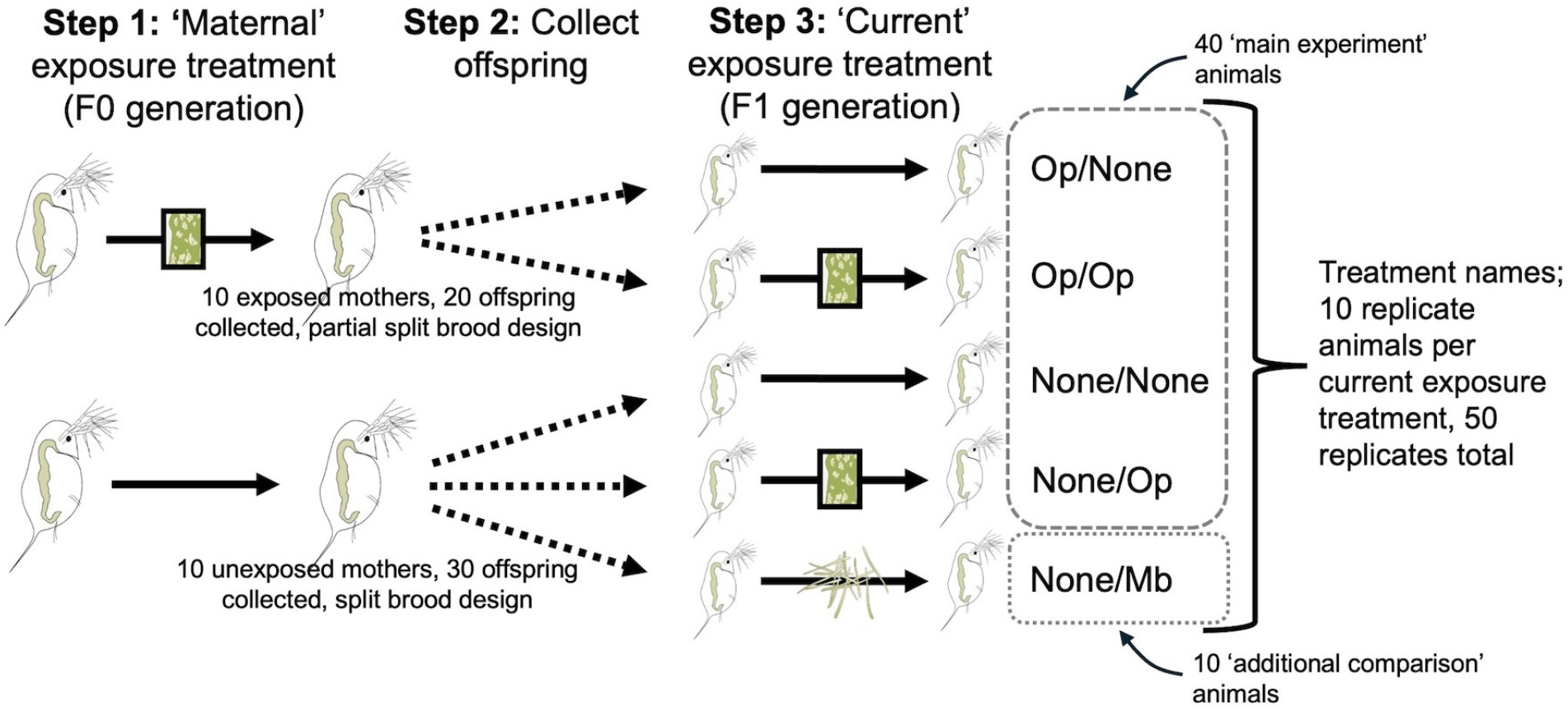
Overview of the five exposure combinations in the empirical portion of the work. We first imposed a maternal exposure treatment on individuals in the maternal generation (i.e., the ‘F0’ generation). Ten of these individuals were exposed to *O. pajunii* (‘Op’) and ten were not exposed to a pathogen (‘None’). We collected three F1 offspring from each unexposed mother; one was exposed to *O. pajunii* in the ‘current exposure’ treatment, one was not exposed, and the third was exposed to the fungus *M. bicuspidata* (‘Mb’). Similarly, we collected offspring from the exposed mothers. We aimed to use a similar split brood design, but with only two F1 offspring per mother, one exposed to *O. pajunii* and one not exposed. However, two mothers only produced one offspring; therefore, we collected two offspring from six mothers (one assigned to the *O. pajunii* treatment, one to the control), one offspring from two mothers, and three offspring from two mothers (split between the two exposure treatments). There were 50 total F1 individuals in the experiment, 40 in the ‘main experiment’ and 10 ‘additional comparison’ animals. *Daphnia* and parasite drawings were created by John Megahan.

#### Maternal exposures

For the mother (F0) generation, we collected 1-day-old animals and placed them individually into 50 mL beakers with 40 mL filtered lake water; this was designated day M1, where M denotes the mother generation. For the duration of the experiment, all experimental beakers were housed in environmental chambers at 20 ^*°*^C with 16:8hrs light:dark. Beakers were maintained within plastic storage containers for ease of handling and to reduce evaporation. Beakers were haphazardly assigned to environmental chambers and locations within chambers daily after checks to control for potential environmental variation among and within chambers. On day M1, we assigned these animals to an exposure treatment. On day M1 and every day thereafter, each beaker received 20,000 cells/mL *Ankistrodesmus falcatus* (AJT strain; Schomaker and Dudycha 2021) as food.

Throughout the experiment, *A. falcatus* was grown in the lab under standard protocols aimed at maintaining consistent food quality over time. 20,000 cells per mL prepared under these conditions is well past the point at which food becomes saturating for *D. dentifera* (Dziuba et al., 2024*a*). On day M5, the animals were transferred to new 50 mL beakers with 10 mL filtered lake water and exposed to either *O. pajunii* or a sham control solution for 48 hours. *O. pajunii* transmission stage spores were collected for this work by homogenization of entire laboratory-infected adult *D. dentifera* (inclusive of the spore-laden gut tissue). We used a ratio of one infected *D. dentifera* per experimental animal to create a spore slurry. Doses were given such that each *O. pajunii* exposed animal received a fraction of the slurry equivalent to 1 infected animal; while we did not quantify the number of spores in the slurry, subsequent measures of spores per infected host indicate that each exposed animal likely received approximately 15,000 spores from this slurry. A similar sham slurry was made for the control solution; this used an equivalent number of laboratory-reared *D. dentifera*, but in this case the animals were uninfected. On Day M7, each animal was moved to a clean, pathogen-free 50 mL beaker with 40 mL filtered lake water. The pathogen exposure window was designed so that *O. pajunii* was removed from the environment prior to brood release by the F0 mothers (and likely prior to embryo production, given expected embryo production on Day M7) to ensure daughters were not exposed during the initial pathogen exposure and that any effects on the F1 generation were transgenerational. This is consistent with a definition of transgenerational as effects seen in subsequent generations after exposure has ended (as in Suarez et al. 2023). We did not quantify infections in the F0 generation individuals who were exposed to *O. pajunii*, but other experiments with these conditions (Dziuba et al., 2024*a*,*b*) and the F1 infections (described below) generally yield 100% infections for this host-pathogen genotype pairing.

#### Current exposures

We collected daughter (F1) generation individuals on Day M9 as 24 hr old neonates from the mother’s first clutch; we designate this Day D1, where D indicates the timings related to the daughter generation. Daughters were isolated individually in 50 mL beakers with 40 mL filtered lake water and 20,000 cells/mL *A. falcatus* food; as in the mother generation, each beaker received this food dose daily throughout the experiment.

On Day D5, we transferred animals to new 50 mL beakers with 10 mL filtered lake water and began the current exposure treatments. ‘Main experiment’ animals were exposed to either *O. pajunii* or a control solution for 48 hours, as described in the ‘Overview of experimental treatments’ section above and Figure 1. *O. pajunii* transmission stage spores were collected and dosed as for the maternal exposures, as was the sham control solution. The 10 ‘additional comparison’ animals were exposed to *M. bicuspidata* (500 spores/mL) for 48 hours, also beginning on Day D5. On Day D7, we moved each individual to a clean, pathogen-free 50 mL beaker with 40 mL filtered lake water.

#### Data collection

We then reared individuals from the daughter (F1) generation, tracking infection, reproduction, and time to mortality. Infection was diagnosed visually under a dissecting microscope (40-110x magnification) beginning at Day 14 and continuing every 5 days thereafter. Animals were considered infected with *O. pajunii* if they showed the distinctive changes in the gut associated with *O. pajunii* infection (Dziuba et al., 2024*b*); all animals that developed signs of infection continued to show signs of infection throughout the rest of the experiment. For any animals that showed signs of *O. pajunii* infection, we quantified burden by imaging the upper midgut of animals diagnosed as infected at 40x magnification using an Olympus DP73 camera. We then overlaid a grid onto the image and counted the number of grid cells containing *O. pajunii* to assess pathogen gut coverage. This was done on the day infection was first diagnosed and every 5 days thereafter until the animal died. We then summed the total spore burden estimate for each day it was measured and divided that by the total number of days the animal lived to get an integrated measure of spore burden over the lifespan. We diagnosed *M. bicuspidata* infections visually using a dissecting microscope. Nine of the ten animals that were exposed to *M. bicuspidata* became infected; the uninfected animal was excluded from analysis, as our focus was comparing the effects of *M. bicuspidata* infection with the transgenerational impacts of *O. pajunii*.

We checked F1 individuals for offspring production and mortality daily. We collected the F2 offspring they produced within 24 hours of birth and measured body length by taking photos under a compound microscope and using Olympus CellSens software to measure individuals. Body length is a proxy for vigor, as it predicts toxicity resistance, starvation resistance, and future offspring production in *Daphnia* (Vesela and Vijverberg, 2007; Lampert, 1993; Gliwicz and Guisande, 1992; Gorbi et al., 2011).

#### Additional experiments

We carried out a follow up experiment to test whether there were differences in maternal provisioning of embryos between F1 individuals whose mothers had been exposed to *O. pajunii* and those whose mothers were not exposed to a pathogen. We also carried out a separate experiment in which we found no evidence of vertical transmission in the *D. dentifera-O. pajunii* system. Methods and results for both of these studies can be found in Appendix S1: Sections S1.2 and S1.3, respectively.

### Statistical Analysis

#### Transgenerational immune priming (TGIP)

TGIP would be indicated by lower infection prevalence or pathogen burden in the F1 daughters of F0 mothers who were exposed to *O. pajunii* (that is, ‘Op/Op’ individuals vs. ‘None/Op’ individuals). To test this, we analyzed the impact of maternal exposure on pathogen burden, which was analyzed as mean gut pathogen coverage per day of life using a Wilcoxon Rank Test with maternal pathogen exposure as the main effect. This analysis used only the 12 individuals from the ‘None/Op’ and ‘Op/Op’ groups that survived to the infection assessment dates and were found to be infected. Unfortunately, due to the high amount of early death in individuals whose mothers had been exposed to *O. pajunii*, we were not able to run formal analyses for infection prevalence.

#### Impacts of transgenerational and current exposure on host mortality

For our analysis of lifespan, we particularly focused on early mortality (within the first 21 days of life); this is when the first three clutches of offspring are produced and is therefore most important to population dynamics (Hutchinson et al., 1978). We assessed the impact of pathogen exposure on the early mortality of F1 generation individuals using restricted mean time analysis. This analysis only allows one independent variable, so we conducted multiple analyses. First, we grouped the 40 main experiment F1 animals based on maternal (F0 generation) exposure (Figure 1), with 20 animals per group. Second, we grouped the 40 main experiment F1 animals based on their ‘current’ exposure treatment (Figure 1), with 20 animals per group. To better isolate the effect of maternal exposure on mortality in unexposed hosts, we also specifically compared the early life mortality of the 10 ‘None/none’ animals with the 10 ‘Op/none’ animals (Figure 1). Finally, to assess the transgenerational impact of *O. pajunii* exposure with the within generation impact of *M. bicuspidata*, we compared the 10 ‘Op/None’ F1 animals to the 9 ‘None/Mb’ F1 animals that became infected with *M. bicuspidata* (Figure 1).

#### Impacts of transgenerational and current exposure on host reproduction

We tested for effects of maternal (F0 generation) exposure and current (F1 generation) exposure on offspring life history traits related to reproduction using Aligned Rank Transform (ART) ANOVA, which allows for nonparametric analyses of variance on factorial models (Wobbrock et al., 2011). We modeled the number of clutches, number of (F2) offspring produced in the first three clutches (for all animals and for only those that produced 3 clutches, separately), and size at first reproduction as the response variables and maternal exposure, current exposure, and their interaction as predictor variables. Transgenerational tolerance would be indicated by higher fitness (e.g., greater reproduction or larger F2 offspring size) in the individuals whose mothers had been exposed to *O. pajunii*.

We also analyzed time to first reproduction by F1 generation individuals with a Cox proportional hazard time to event analysis, considering the main effects of maternal pathogen exposure and current exposure and their interaction. Finally, we analyzed (F2) offspring neonate size. Similar to the analyses in the previous paragraph, this used an ART ANOVA. In addition to having maternal exposure, current exposure, and their interaction as predictor variables, we also included the identity of the F1 mother in the analysis to account for non-independence of offspring produced from a single mother. Details of statistical analyses (including sample sizes and R packages used) can be found in Appendix S1: Table S1. All statistical analyses were conducted in R (version 4.4.0).

## Empirical Results

We did not see evidence of transgenerational immune priming (TGIP): maternal exposure did not impact infection intensity in the F1 individuals who were exposed to *O. pajunii* (W=10, P= 0.350; Table 1). While we could not conduct a formal statistical analysis on infection prevalence due to high mortality in the ‘Op/Op’ treatment, there was no clear signature of an impact of maternal exposure on infection prevalence in exposed offspring (infection proportions: 8 out of 8 animals infected for ‘None/Op’ treatment, 4 out of 5 animals infected for ‘Op/Op’ treatment). However, we acknowledge that our limited sample size reduced our statistical power to detect small effect sizes.

**Table 1:**
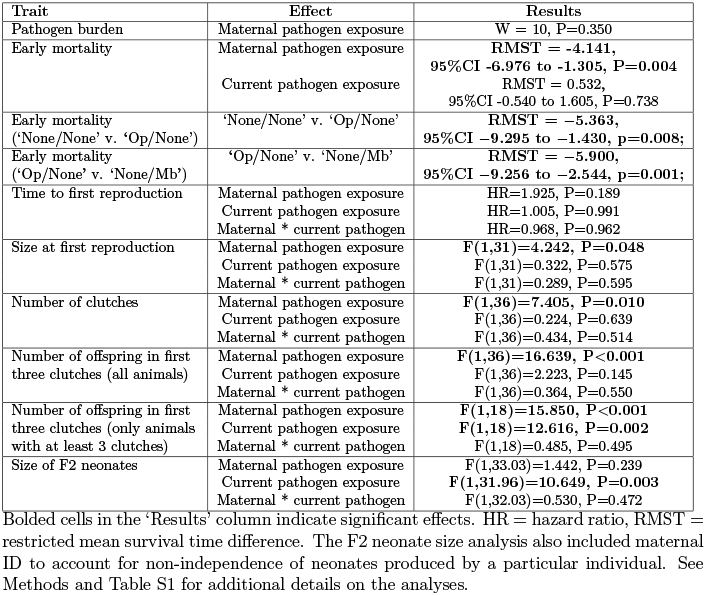
Summary of statistical results for analyses of main experiment animals.

| Trait | Effect | Results |
| --- | --- | --- |
| Pathogen burden | Maternal pathogen exposure | W = 10, P=0.350 |
| Early mortality | Maternal pathogen exposure | **RMST = -4.141,**<br>**95%CI -6.976 to -1.305, P=0.004** |
|  | Current pathogen exposure | RMST = 0.532,<br>95%CI -0.540 to 1.605, P=0.738 |
| Early mortality<br>(‘None/None’ v. ‘Op/None’) | ‘None/None’ v. ‘Op/None’ | **RMST = -5.363,**<br>**95%CI -9.295 to -1.430, p=0.008;** |
| Early mortality<br>(‘Op/None’ v. ‘None/Mb’) | ‘Op/None’ v. ‘None/Mb’ | **RMST = -5.900,**<br>**95%CI -9.256 to -2.544, p=0.001;** |
| Time to first reproduction | Maternal pathogen exposure | HR=1.925, P=0.189 |
|  | Current pathogen exposure | HR=1.005, P=0.991 |
|  | Maternal * current pathogen | HR=0.968, P=0.962 |
| Size at first reproduction | Maternal pathogen exposure | **F(1,31)=4.242, P=0.048** |
|  | Current pathogen exposure | F(1,31)=0.322, P=0.575 |
|  | Maternal * current pathogen | F(1,31)=0.289, P=0.595 |
| Number of clutches | Maternal pathogen exposure | **F(1,36)=7.405, P=0.010** |
|  | Current pathogen exposure | F(1,36)=0.224, P=0.639 |
|  | Maternal * current pathogen | F(1,36)=0.434, P=0.514 |
| Number of offspring in first<br>three clutches (all animals) | Maternal pathogen exposure | **F(1,36)=16.639, P<0.001** |
|  | Current pathogen exposure | F(1,36)=2.223, P=0.145 |
|  | Maternal * current pathogen | F(1,36)=0.364, P=0.550 |
| Number of offspring in first<br>three clutches (only animals<br>with at least 3 clutches) | Maternal pathogen exposure | **F(1,18)=15.850, P<0.001** |
|  | Current pathogen exposure | **F(1,18)=12.616, P=0.002** |
|  | Maternal * current pathogen | F(1,18)=0.485, P=0.495 |
| Size of F2 neonates | Maternal pathogen exposure | F(1,33.03)=1.442, P=0.239 |
|  | Current pathogen exposure | **F(1,31.96)=10.649, P=0.003** |
|  | Maternal * current pathogen | F(1,32.03)=0.530, P=0.472 |

Of the 40 main experiment animals, the strongest predictor of offspring early life survival (that is, survival through day 21 in the F1 generation) was maternal exposure to *O. pajunii* : F1 offspring of F0 mothers that had been exposed to the microsporidian were much more likely to die early in life compared to those whose mothers had not been exposed (restricted mean survival time (RMST) difference: −4.141, 95% CI −6.976 to −1.305, p=0.004; Table 1). This reduction in early life survival occurred regardless of whether they had themselves been exposed to *O. pajunii* (Figure 2); there was not a significant effect of current exposure on early life mortality (RMST difference: 0.532, 95%CI −0.540 to 1.605, p=0.738; Table 1). Looking just at F1 individuals who were not exposed to the pathogen (that is, comparing ‘None/None’ individuals vs. ‘Op/None’ individuals), early mortality was still significantly greater for individuals whose mothers had been exposed (RMST difference = −5.363, 95%CI −9.295 to −1.430, p=0.008; Figure 2). Surprisingly, the transgenerational impact of maternal *O. pajunii*-exposure on early life mortality was stronger than the within generation impact of infection by *M. bicuspidata* (RMST difference of ‘Op/None’ vs. ‘None/Mb’ individuals: −5.900, 95%CI −9.256 to −2.544, p=0.001; Figure 2), which is considered highly virulent due to its impact on host lifespan.

**Figure 2:**
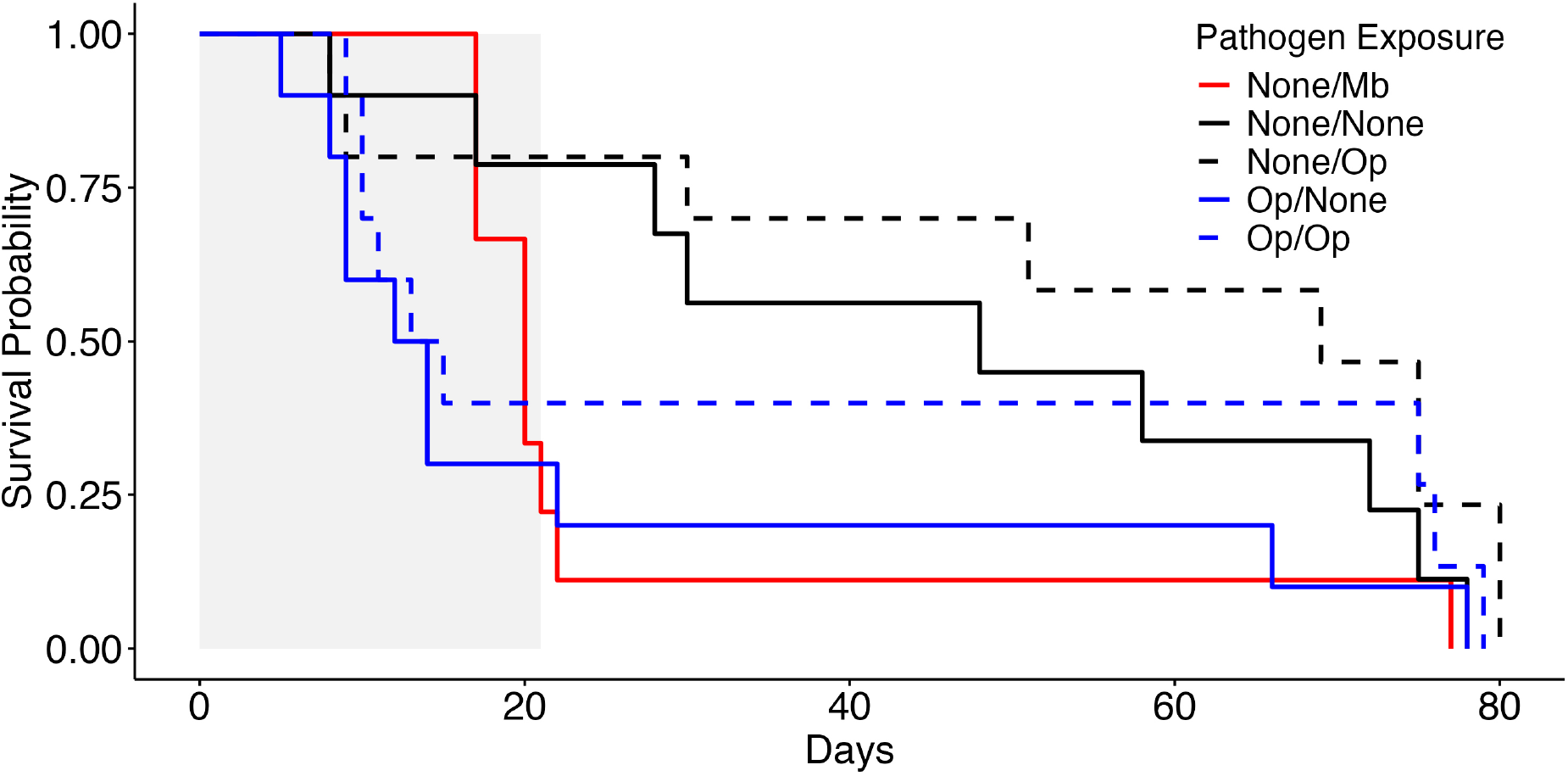
Daughters of exposed mothers had reduced lifespan. The figure shows survivorship curves for all experimental treatments. The ‘Pathogen Exposure’ legend labels correspond to the treatments indicated in Figure 1. Main experiment animals are shown with black and blue lines: blue lines represent offspring of mothers that had been exposed to *O. pajunii*, whereas black lines indicate main experiment animals whose mothers were not exposed; dashed lines represent current *O. pajunii* exposure, while solid lines represent no current *O. pajunii* exposure. The red line indicates ‘additional comparison’ animals which were exposed to the virulent fungus *M. bicuspidata*. Grey shading indicates the critical early life period (through day 21).

Maternal exposure did not significantly impact time to first reproduction in F1 individuals (hazard ratio (HR)=1.925, p=0.189; Table 1), but F1 daughters of *O. pajunii*-exposed mothers were smaller at first reproduction (F_(1,31)_=4.24, p=0.048; Figure 3; Table 1), indicating reduced growth rate. Additionally, maternal exposure substantially reduced the overall number of clutches produced by F1 daughters (F_(1,36)_=7.41, p=0.010; Figure 3; Table 1) and the number of offspring they produced in the first 3 clutches (for all animals: F_(1,36)_ = 16.64, P<0.001; Figure 3; for only animals that produced 3 clutches: F_(1,18)_=15.85, p=0.001; Table 1). The size of F2 neonates was only significantly influenced by current (F1 generation) pathogen exposure (F_(1,31.96)_= 10.649, p=0.003; Figure 3; Table 1).

**Figure 3:**
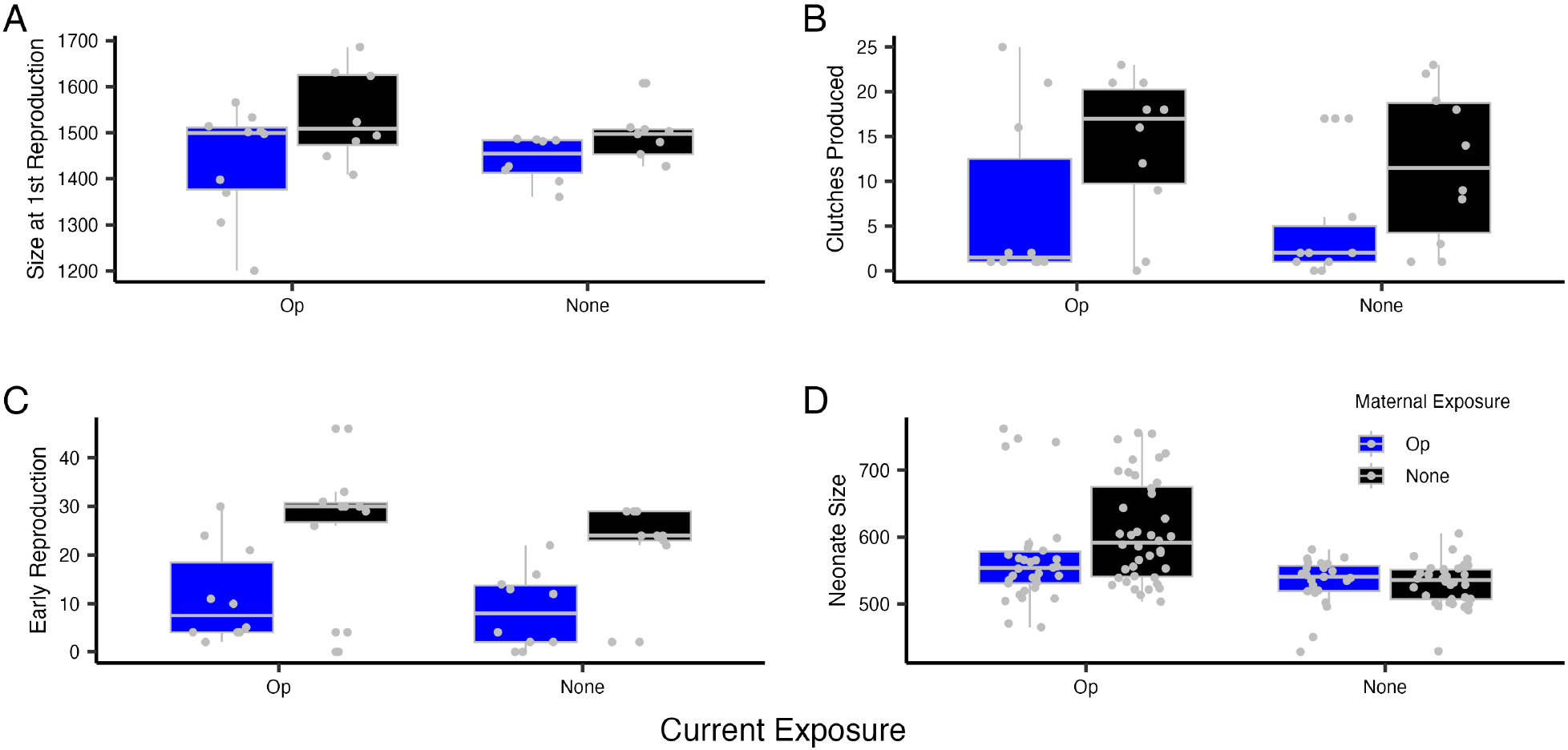
Maternal exposure to *O. pajunii* reduced growth of and reproduction by offspring, regardless of their current infection status. Individuals whose mothers were exposed to *O. pajunii* were smaller at first reproduction (A). They also produced fewer clutches (B) and had fewer offspring in the first three clutches (‘early reproduction’, panel C). The data plotted in C are for all main experiment animals; there was still an effect of maternal pathogen exposure on the number of offspring in the first three clutches for just the subset of animals that produced at least three clutches (Table 1). The size of F2 neonates (measured in micrometers) produced by F1 animals was influenced by current (F1) pathogen exposure (D). The small gray circles on panel D show each individual neonate, but the statistical model included a term for the F1 individual who produced them. ‘Op’ = *Ordospora pajunii*

## Mathematical Modeling Methods

The empirical results suggest that, rather than transgenerational immune priming or transgenerational tolerance, offspring of exposed mothers had reduced fitness. To explore the population-level effects of these transgenerational impacts, we analyzed an SIR-type environmental transmission model that describes the dynamics of the densities of: susceptible (*S*) and infected (*I*) individuals who are the offspring of susceptible individuals; susceptible (*C*) and infected (*D*) individuals who are the offspring of infected individuals (hereafter, compromised and decimated individuals, respectively); and infectious propagules in the environment (*P*). Note that the *S* class corresponds to ‘None/None’ animals in the experiment (Figure 1), *I* corresponds to ‘None/Op’ individuals, *C* corresponds to ‘Op/None’ individuals, and *D* corresponds to ‘Op/Op’ individuals that were successfully infected. As in the *Daphnia-O. pajunii* system, the model assumes infection occurs when susceptible or compromised individuals encounter infectious propagules that were shed previously into the environment by infected or decimated individuals.

The model equations are

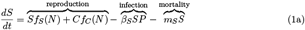

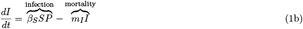

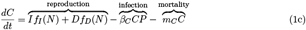

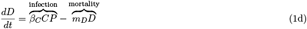

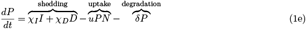

Table 2 summarizes the model notation. In equation (1a), susceptible density increases due to reproduction by susceptible (*Sf*_*S*_(*N*)) and compromised (*Cf*_*C*_(*N*)) individuals; for simplicity, we model the density-dependent reproduction rates using a Lotka-Volterra form, *f*_*i*_(*N*) = *r*_*i*_(1− *N/K*), where *r*_*i*_ is the maximum reproduction rate for individuals in class *i, K* is the host’s carrying capacity, and *N* = *S* + *C* + *I* + *D* is total host density. The other terms in equation (1a) account for decreases in susceptible density due to infection (*β*_*S*_*SP*) and non-disease mortality (*m*_*S*_*S*). In equation (1b), infected density increases when susceptible individuals become infected (*β*_*S*_*SP*) and decreases due to mortality from disease and non-disease sources (*m*_*I*_*I*). In equation (1c), compromised density increases due to reproduction by infected individuals (*If*_*I*_(*N*)) and decimated individuals (*Df*_*D*_(*N*)) and decreases due to infection (*β*_*C*_*CP*) and mortality due to non-disease sources (*m*_*C*_*C*); the per capita reproduction rates have the same Lotka-Volterra form as above. In equation (1d), decimated density increases when compromised individuals become infected (*β*_*C*_*CP*) and decreases due to mortality from disease and non-disease sources (*m*_*D*_*D*). In equation (1e), infectious propagule density increases due to shedding by infected and decimated individuals (*χ*_*I*_*I* + *χ*_*D*_*D*) and decreases due to uptake by all individuals (*uPN*) and degradation (*δP*). Importantly, to account for the transgenerational effects of infection, when compared to susceptible individuals, compromised individuals have (i) equal or higher mortality rates (*m*_*C*_ ≥ *m*_*S*_), (ii) equal or lower reproductive output (*f*_*C*_ ≤ *f*_*S*_), and (iii) equal or higher transmission rates (*β*_*C*_ ≥ *β*_*S*_).

**Table 2:**
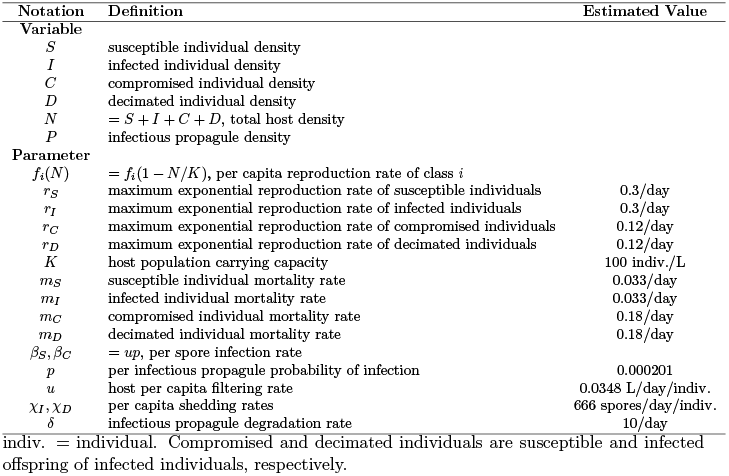
Definitions and estimated values for model parameters and variables.

The simplifying assumptions built into our model include infected and decimated individuals shed identical infectious propagules, susceptible and compromised offspring have identical offspring (i.e., there are no grandmaternal effects of infection), and all individuals have equal uptake rates (*u*). We also assume the negative effects of host density on reproduction rates are independent of the make-up of the community. Said another way, the intraspecific competitive effects of all individuals are identical, regardless of infection or maternal status. Mathematically, this corresponds to assuming all per capita reproductions rates, *f*_*i*_(*N*), only depend on total host density, not the densities of the infected and susceptible classes. In addition, we assume infection causes infected and decimated individuals to have equal or lower reproduction rates than susceptible and compromised individuals, respectively (*r*_*I*_ ≤ *r*_*S*_ and *r*_*D*_ ≤ *r*_*C*_).

To parameterize model (1) for the *D. dentifera-0. pajunii* system, we estimated model parameters using the data from the main experiment. The only exceptions are that the values for the susceptible host reproduction rate (*r*_*S*_), carrying capacity (*K*), and uptake rate (*u*) were taken from a previous study on the host *D. dentifera* (Searle et al., 2016). Appendix S1: Section S3.1 provides details about how parameters were estimated and Table 2 lists the estimated values. Our estimated parameters match the experimental data in the following ways. First, susceptible and infected individuals have equal reproduction rates (*r*_*S*_ = *r*_*I*_) and equal mortality rates (*m*_*S*_ = *m*_*I*_) because infection does not significantly affect host reproduction or lifespan. Second, when compared to susceptible and infected individuals, compromised and decimated individuals have smaller reproduction rates (*r*_*C*_, *r*_*D*_ < *r*_*S*_, *r*_*I*_) and larger mortality rates (*m*_*C*_, *m*_*D*_ > *m*_*S*_, *m*_*I*_) because maternal exposure status reduced offspring reproduction by approximately 60% and increased mortality rate by approximately 5.5 times. Third, the infection rates (*β*_*S*_ = *β*_*C*_), shedding rates (*χ*_*D*_ = *χ*_*I*_), and probabilities of infection (*u*) are the same across classes because those processes did not depend on maternal infection status.

We explored the population-level effects of the transgenerational fitness impacts of pathogen exposure in the *D. dentifera-0. pajunii* system in the following way. First, we simulated model (1) using our estimated values in Table 2. This simulation includes the transgenerational effects of infection because compromised individuals have lower reproduction rates and higher mortality rates than susceptible individuals (*r*_*C*_ < *r*_*S*_ and *m*_*C*_ > *m*_*S*_, respectively). Then we simulated model (1) parameterized such that susceptible and compromised individuals have equal reproduction and mortality rates (*r*_*C*_ = *r*_*S*_ and *m*_*C*_ = *m*_*S*_, respectively). This simulation has no transgenerational effects of infection. Comparing the dynamics of the two simulations allows us to identify how the transgenerational impacts of pathogen exposure on reproduction and mortality alter the disease dynamics of the *Daphnia-0. pajunii* system.

To assess the generality of the patterns predicted for the *Daphnia-0. pajunii* system and to make predictions that could apply to other systems, we explored how transgenerational impacts alter disease dynamics across parameter space in model (1). Following prior studies (Cortez, 2021; Cortez and Duffy, 2021), we used the Jacobian-based methods in (Yodzis, 1988) to compute the local sensitivities of model equilibria to variation in parameters. Specifically, we computed explicit formulas for the local sensitivities of total host density (*N* ^∗^), total infected density (*I*^∗^ + *D*^∗^), and infection prevalence ([*I*^∗^ + *D*^∗^]*/N* ^∗^) at equilibrium to the parameter values for the reproduction (*r*_*C*_), mortality (*m*_*C*_), and transmission (*β*_*C*_) rates of compromised individuals. These sensitivities are derivatives whose signs determine how changes in a parameter value affect the equilibrium values of the model. For example, if the transgenerational impact increases mortality of compromised individuals, then the derivative *∂N* ^∗^*/∂m*_*C*_ determines whether total host density at equilibrium increases (positive values) or decreases (negative values) because of those transgenerational effects. Similarly, the derivatives of total infected density (*∂*[*I*^∗^ + *D*^∗^]*/∂m*_*C*_) and infection prevalence (*∂*[(*I*^∗^ + *D*^∗^)*/N* ^∗^]*/∂m*_*C*_) at equilibrium determine if those metrics of disease increase or decrease. Because the high dimensionality of model (1) limits the tractability of the sensitivity formulas, in the appendices, we first analyzed a reduced model where infected and decimated individuals are identical; see model (S1a) of Appendix S1: Section S2.1 and its analysis in Appendix S1: Sections S2.2-S2.3. Then we analyzed the full model (1) in Appendix S1: Sections S2.4-S2.5. The reduced model facilitates interpretation by reducing the complexity of the sensitivity formulas; the predictions from the two models are qualitatively similar.

## Mathematical Modeling Results

Our parameterized model predicts that the transgenerational effects of *O. pajunii* reduce both host densities and disease levels. Specifically, Figure 4 compares the model dynamics when compromised individuals have lower reproduction rates and higher mortality rates than susceptible individuals (transgenerational effects present; blue curves in Figure 4) with the model dynamics when compromised and susceptible individuals have identical reproduction and mortality rates (transgenerational effects absent; black curves in Figure 4). In those simulations, the transgenerational effects of *O. pajunii* reduce host density by 30.3% (Fig. 4A), reduce infected host density by 45.4% (Fig. 4B), and reduce infection prevalence by 21.7% (Fig. 4C). A more complete exploration of the model that is parameterized for the *D. dentifera-O. pajunii* system can be found in Appendix S1: Section S3.2.

**Figure 4:**
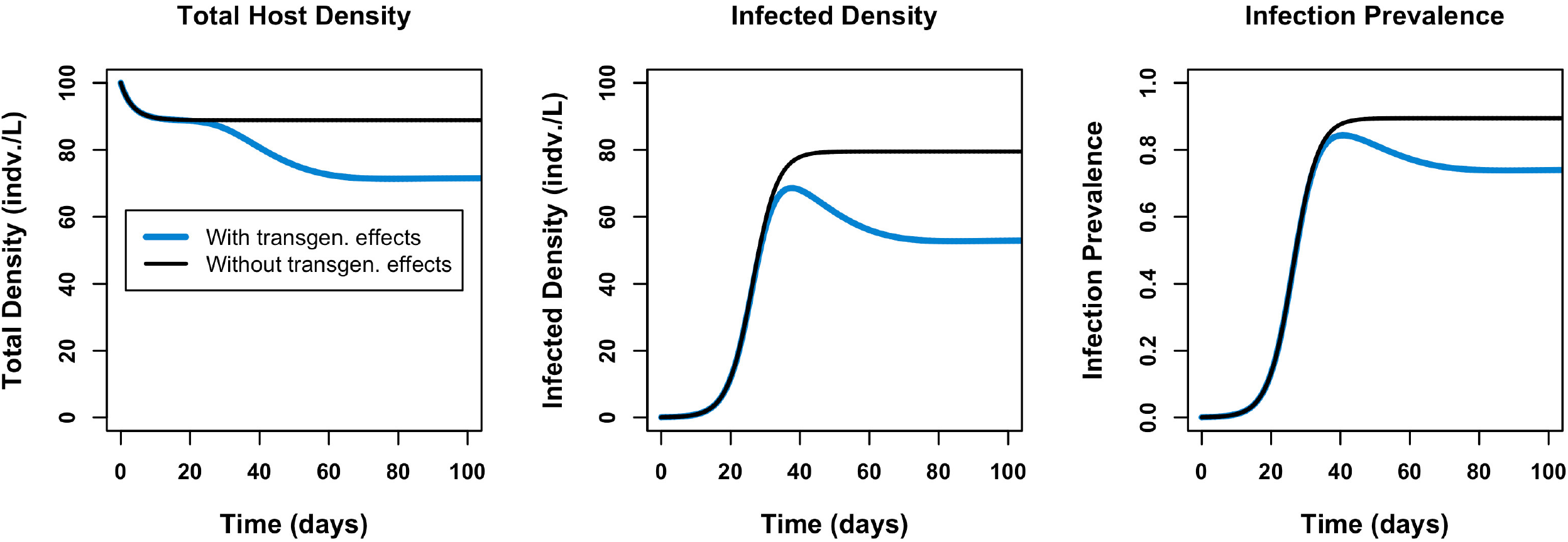
Transgenerational effects of pathogen exposure are predicted to notably reduce total host density (A), infected host density (B), and infection prevalence (C). Thinner black lines show times series from a model parameterized to the *Daphnia-O. pajunii* system where there are no transgenerational impacts of pathogen exposure. Thicker blue lines show time series from a model with transgenerational impacts, where offspring of infected individuals have higher mortality and lower reproduction than the offspring of uninfected individuals.

Our sensitivity analysis of model (1) suggests that transgenerational effects are likely to impact host and disease dynamics in other systems as well. We first consider scenarios where the transgenerational impact of infection is greater mortality or reduced reproduction, which aligns with our experimental findings. We also consider the possibility that maternal exposure increases the susceptibility of offspring; this was not seen in our experiment, but might occur in other systems. Here, we summarize our main results and refer to reader to Appendix S1: Sections S2.2 and S2.4 for additional mathematical details.

If the transgenerational impact causes greater mortality (*m*_*C*_ > *m*_*S*_) or reduced reproduction (*r*_*C*_ < *r*_*S*_) in compromised individuals when compared to susceptible individuals, then we predict decreases in total host density (*N* ^∗^), infection prevalence ([*I*^∗^ + *D*^∗^]*/N*), and total infected density (*I*^∗^ + *D*^∗^) at equilibrium as a result of those transgenerational effects. The intuition is that greater mortality and reduced reproduction each decrease the total growth rate of the host population, resulting in lower total host densities at equilibrium. The decreases in total host density in turn leave fewer individuals that can become infected, which reduces infection rates and leads to lower proportions and densities of infected individuals.

If the transgenerational impact causes increased infection rates for compromised individuals when compared to susceptible individuals (*β*_*C*_ > *β*_*S*_), then we predict decreases in total host density and increases in infection prevalence ([*I*^∗^ + *D*^∗^]*/N* ^∗^) and total infected density (*I*^∗^ + *D*^∗^) at equilibrium. Here, the intuition is that higher infection rates lead to greater densities of decimated individuals and the resulting increase in infectious propagule density (due to shedding by decimated individuals) causes an increase in infected individuals. Combined, these two effects imply an increase in total infected density. The increase in total infected density causes total host density to decrease because more individuals are becoming infected and dying via disease-induced mortality. The combination of higher total infected density (*I*^∗^ + *D*^∗^) and lower total host density (*N* ^∗^) necessarily leads to higher infection prevalence ([*I*^∗^ + *D*^∗^]*/N* ^∗^).

In summary, if transgenerational impacts of infection manifest as decreased reproduction or increased mortality, as was seen in our experiment, then that harms both the host population (via reduced densities) and the pathogen (via reduced infected density and infection prevalence). If transgenerational impacts of infection manifest instead as increased infection rates, then that harms the host population (via reduced densities) and benefits the pathogen (via higher infected density and infection prevalence). If it manifests as all three effects, then it harms the host population, but it can harm or benefit the pathogen depending on how much the reproduction, mortality, and infection rates are altered.

## Discussion

We carried out an experiment looking for transgenerational impacts of pathogens. Rather than finding that the offspring of exposed mothers were more resistant to or tolerant of infection, we found that offspring of exposed mothers had lower fitness (dying early and reproducing less) compared to daughters of unexposed mothers. Uninfected offspring of mothers who were exposed to the microsporidian *O. pajunii* had significantly higher early life mortality than individuals who were currently infected with the fungus *M. bicuspidata*; this means the transgenerational impact of *O. pajunii* exceeds the within generation impact of a pathogen that is generally considered highly virulent because of its effects on mortality. Our parameterized mathematical model reveals that these transgenerational impacts can alter population- and community-level dynamics: increased mortality or reduced fecundity of the offspring of exposed mothers can reduce total host density, infected host density, and infection prevalence.

While there is ample evidence that immune priming can incur costs in terms of reduced growth rate and fecundity (reviewed in Tetreau et al., 2019), our finding is notable because there was no evidence for successful immune priming: exposed daughters of exposed mothers (‘Op/Op’ animals) became infected at high rates, as did exposed daughters of unexposed mothers (‘None/Op’ animals), and there was no significant difference between these two groups in infection burden. Additionally, we did not see signals of transgenerational tolerance. Instead, our data indicate only costs (and not benefits) of maternal exposure to offspring. We do not have data from our study showing the ultimate infection status of the exposed mothers, but the host-pathogen genotype pairing used in this study generally leads to extremely high levels of infection (as seen in this study), particularly under the high food conditions used in this study (Dziuba et al., 2024*a*,*b*). This suggests that the transgenerational impact we observed is a type of virulence – that is, a reduction in host fitness caused by infection (Cressler et al., 2016).

We propose that the effects we measured could be considered transgenerational virulence, where the costs of pathogen exposure manifest in the offspring of the exposed hosts. We propose three criteria for something to be considered transgenerational virulence. First, the impact of infection should occur in offspring that are not themselves infected (or even exposed, which differs from an example like Zika virus, where the offspring becomes infected; Freitas et al., 2020). Second, offspring fitness (e.g., lifespan or reproduction) should be reduced. And, third, the impact should not be due to offspring demise during gestational development as a result of the maternal infection (as occurs with SARS-CoV-2 infections where the placenta is damaged; Schwartz et al., 2022). As far as we are aware, the concept of transgenerational virulence is new, but it seems likely that this effect exists in other systems. Indeed, a study on sticklebacks meets these criteria, where offspring of infected fathers had reduced hatching success and survival (Kaufmann et al., 2014). We propose that further development of the concept of transgenerational virulence – including the mechanisms underlying it, how to quantify it, and its ecological and evolutionary implications – is likely to be fruitful.

Temporarily reducing reproductive investment after pathogen exposure might be a strategy that allows a host to divert resources to an immune response. If exposure to the pathogen caused mothers to reduce investment in (but not stop producing) embryos, this might yield offspring with reduced early survival. In our experiment, the most notable impact was on early mortality of offspring of exposed mothers; this is consistent with the paradigm that any reduction in fitness due to reduced maternal investment would manifest early in life (Tetreau et al., 2019). However, results from additional studies we carried out argue that the transgenerational effects we observed are not simply due to immune activation. Maternal exposure of this host species to two other pathogens, *M. bicuspidata* and *Pasteuria ramosa* (a bacterium) did not result in increased early life mortality of uninfected offspring (McIntire et al, in prep). While this suggests that the transgenerational virulence of *O. pajunii* is not due to immune activation (since *M. bicuspidata* and *P. ramosa* also trigger immune responses; Stewart Merrill and Cáceres, 2018; Auld et al., 2010), it remains possible that there are differences in the nature of the immune responses to the different pathogens; future studies that better quantify immune responses to different pathogens would help address this.

Studies on transgenerational effects of stressors other than pathogens point to potential mechanisms that might underlie our empirical findings. First, molecular mechanisms, such as epigenetic inheritance, often influence transgenerational shifts in response to stressors, as evidenced in a wide variety of taxa; for example, environmental stress-induced epigenetic changes negatively impact offspring fitness in Polpay sheep (due to diet stress; Braz et al., 2022) and *C. elegans* (due to toxicant exposure; Wei et al., 2020). Second, organisms can shift energetics in response to stressors; for example, altering reproduction under conditions of environmental stress (or parasitism) is a well characterized strategy to maximize fitness. While this is commonly seen as changes in the number of offspring (Tessier and Consolatti, 1991; Leventhal et al., 2014), it can also manifest as alterations of nutrients maternally partitioned to offspring (as in *Daphnia*, with reduced egg fat content after predator signals; Stibor and Lüning, 1994; Stibor and Navarra, 2000). In our system, there was evidence of greater fragility of embryos produced by *O. pajunii*-exposed mothers, but there were no detectable differences in embryo or yolk size (Appendix S1: Section S1.2); future work to understand the drivers of the greater fragility of these embryos (e.g., due to changed chemical composition) might help understand the mechanisms underlying the lower fitness of compromised offspring. It is also worth noting that these two mechanisms – that is, molecular/epigenetic changes and altered energetic allocations – may combine: in fruit flies, high sugar maternal diets altered body composition of offspring and modified metabolic gene expression, with fitness impacts carrying throughout the offspring lifespan (Buescher et al., 2013).

Transgenerational effects have been demonstrated to vary by genotype across taxa (Rendina González et al., 2018; Dew-Budd et al., 2016; Alvarez et al., 2020). For example, *Drosophila* genotypes show diverse metabolic transgenerational responses to diet-related stressors (Dew-Budd et al., 2016). Genotypic variation in responses is highly likely in this system, given that the within-generation susceptibility and fitness response to *O. pajunii* exposure vary (Dziuba et al., 2024*b*). These differences have the potential to scale up to host population-level differences in the ability to compensate for transgenerational virulence-driven increases in mortality (Capdevila et al., 2020), with potential implications for host population genetic composition and infection prevalence — providing further evidence of the importance of considering host life history in disease ecology (Valenzuela-Sánchez et al., 2021).

A major concern at the intersection of disease and community ecology is the potential for pathogen spillover (Wells and Clark, 2019). Accurate assessment of spillover risk is critically important, as more than 60% of emerging infectious disease stems from spillover (Jones et al., 2008). Host density is a significant factor in spillover risk (Plowright et al., 2017), and the predicted decreases in total host density from our model may lead to decreases in spillover. Transgenerational virulence also reduced infected host density, which should further reduce spillover risk in systems with density-dependent transmission (Greer et al., 2008). Therefore, transgenerational virulence-related density changes could alter both outbreak prediction and management success (Mysterud et al., 2023). If the host is of conservation concern, the density reduction that results from transgenerational virulence (Figure 4) would mean that the pathogen suppresses host density beyond what would be expected based solely on the within generation impacts of the pathogen. However, if a key conservation concern is spillover of infection from a different host, the reduction in infection prevalence and infected host density that result from transgenerational virulence could reduce spillover risk.

Here, we demonstrated that maternal exposure to a microsporidian pathogen strongly reduces offspring fitness. We propose that the decreased survival and reduced fecundity of the offspring of exposed (and most likely infected) mothers can be considered transgenerational virulence. Our results suggest that the transgenerational virulent impacts of *O. pajunii* are similar to (or exceed) the within generation impacts of a different, highly virulent pathogen. Moreover, they have consequences at the population-level: our model reveals that transgenerational virulence can notably decrease host density, infected host density, and infection prevalence. Given the widespread prior evidence for transgenerational effects of stressors including pathogens, we propose that transgenerational virulence is likely to occur in other host-pathogen systems, with consequences not only for individual fitness, but also for population dynamics for both hosts and pathogens.

## Acknowledgements

We thank members of the Duffy Lab, particularly S-J. Sun, S. Calhoun, K. Monell, and T. Sauer for logistic and experimental support. We also thank anonymous reviewers for helpful feedback on earlier versions of this manuscript. This work was supported by funding from the Gordon and Betty Moore Foundation (GBMF9202 to MAD; https://doi.org/10.37807/GBMF9202) and the US National Science Foundation (DEB-1748729 to MAD, DEB-2015280 to MHC, and GRF 1937954 to MLR).

## Author contributions

Conceptualization: KMM & MAD; Methodology: KMM, MKD, MHC; Empirical data collection: KMM, MKD, EB, MV, FEC, TN; Analysis of empirical data: KMM; Model development and analysis: MHC, EBH, MLR; Funding: MAD & MHC; Resources: MAD; Writing-original draft: KMM; Writing-substantial edits: MAD, KMM, MHC; Writing-review & editing: KMM, MAD, MKD, MHC

## Appendix S1

### Section S1 Additional empirical methods and results

#### Section S1.1 Additional information regarding analyses presented in main text

In Table S1, we present additional information regarding the statistical methods used to analyze the results of the main experiment.

**Table S1:**
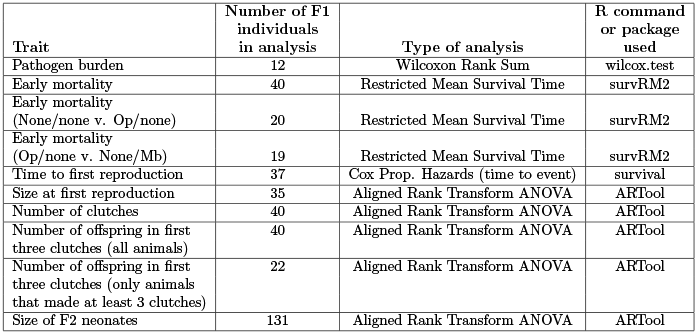
Summary of statistical methods. Results are given in the main text, including in Table 1. References for R packages are: survRM2 (Uno et al., 2014; Tian et al., 2014), survival (Therneau and Grambsch, 2000; Therneau, 2024), and ARTool (Kay et al., 2021; Wobbrock et al., 2011). Treatment names given in parentheses correspond with those in the main text Figure 1.

| Trait | Number of F1 individuals in analysis | Type of analysis | R command or package used |
| --- | --- | --- | --- |
| Pathogen burden | 12 | Wilcoxon Rank Sum | wilcox.test |
| Early mortality | 40 | Restricted Mean Survival Time | survRM2 |
| Early mortality (None/none v. Op/none) | 20 | Restricted Mean Survival Time | survRM2 |
| Early mortality (Op/none v. None/Mb) | 19 | Restricted Mean Survival Time | survRM2 |
| Time to first reproduction | 37 | Cox Prop. Hazards (time to event) | survival |
| Size at first reproduction | 35 | Aligned Rank Transform ANOVA | ARTool |
| Number of clutches | 40 | Aligned Rank Transform ANOVA | ARTool |
| Number of offspring in first three clutches (all animals) | 40 | Aligned Rank Transform ANOVA | ARTool |
| Number of offspring in first three clutches (only animals that made at least 3 clutches) | 22 | Aligned Rank Transform ANOVA | ARTool |
| Size of F2 neonates | 131 | Aligned Rank Transform ANOVA | ARTool |

#### Section S1.2 Experimental test for maternal care during embryonic development

As described in the main text, we found that F1 offspring of mothers who had been exposed to *O. pajunii* (who we refer to as ‘compromised’ offspring) had higher early life mortality than F1 offspring of mothers who had not been exposed to a pathogen (who we refer to as ‘uncompromised’ offspring). We carried out additional studies to identify potential mechanistic causes of the reduced fitness of compromised offspring. We hypothesized that *O. pajunii*-exposed mothers provisioned embryos differently, resulting in lower neonate survival. We predicted that *O. pajunii*-exposed mothers would produce smaller embryos or embryos with less maternal yolk.

For this experiment, we acclimated S genotype *D. dentifera* grandmothers in common conditions to minimize unintended variation within the mother generation. We collected the mother generation (on Day 1) as 24 hr old neonates, placing them individually in 50 mL beakers with 20 mL filtered lake water, and assigning each to one of two pathogen exposure treatments (pathogen exposed or control) and one of three embryo care treatments (see groups A, B, and C, below). Mother beakers received 0.5 mL *Ankistrodesmus falcatus* (AJT strain, 20,000 cells/mL) as food daily.

Pathogen exposure treatments consisted of a 48 hr exposure, beginning on Day 2, to a dose of *O. pajunii* spore slurry or a dose of placebo slurry. We created the spore slurry by crushing individuals with mature (spore producing) infections into DI water in a ratio of 1 infected animal per experimental animal; a sham/control slurry was created similarly with uninfected animals. After this exposure ended, we kept mothers in 40 mL filtered lake water to relieve density constraints with food increased to 1 mL daily. Beginning on Day 5, animals were monitored 4 times daily for early stage (stage 1, indicating the first stage of embryo development) embryo production or freshly deposited eggs. When stage 1 production was noted, we recorded the time and number of embryos and imaged mothers for body size (a significant predictor of offspring size, (Gabsi et al., 2014)), after which we proceeded according to the mother’s embryo care treatment. <u>For group A</u>, we removed embryos from the maternal brood chamber; these were imaged for size (length and width) and yolk diameter (Urabe and Sterner, 2001). This treatment allowed for comparison of immediate maternal resource volume investment across compromised and uncompromised individuals. <u>For group B</u>, we removed embryos from the mother’s brood chamber and transferred them to a well in a 24 well plate with 2 mL filtered lake water. <u>For group C</u>, embryos remained in the brood chamber to develop. The release of embryos from the brood chamber marks the transition from being an embryo to a neonate. When they were still in stage 1, each clutch of group B was paired with a clutch of group C (within a given pathogen treatment group). This was used to control for group B having no release event to mark passage from embryo to neonate; instead, when group C was released from the brood chamber, both that clutch and the one from the pair in Group B were prepared for measurement. Comparison of groups B and C allowed us to check for additional maternal provisioning later in embryonic growth, after initial embryo production.

Embryos were checked twice daily; when embryos were released, we recorded the number of surviving neonates, measured neonate body length, and preserved neonates for weighing. Neonates within a maternal pathogen treatment were weighed in random cohorts of up to 10 animals (weighing neonates individually was impossible due to insufficient sensitivity of the balance). Due to logistical challenges associated with weighing neonates, we only weighed neonates from Group C.

##### Section S1.2.1 Data Analysis

We compared the number of embryos produced for all individuals using a Wilcoxon test with maternal parasite exposure as the factor; embryo care treatment was not included in this analysis as it had not occurred at the time of this data collection. We also compared the number of neonates produced by group B and C mothers and the deficit between embryos produced and neonates that survived, per individual, using separate aligned rank transform ANOVA (ARTtool package) with maternal parasite exposure, embryo care treatment, and their interaction as fixed effects.

Within group A, we analyzed the mean length (per clutch) and the mean yolk/mean length ratio of each clutch via t-test with maternal exposure as a factor; we analyzed within clutch embryo size range, yolk width range (ranges considered representative of within clutch variation), mean yolk width (per clutch), maternal size, and the mean length/maternal size ratio of each clutch by Wilcoxon rank sum test with maternal exposure as a factor. We analyzed data on within clutch variation based on Marshall et al. (2008), which found greater variation in offspring size within a mother in unpredictable environments, indicating that variation in offspring size could be an adaptive strategy in the face of environmental challenge.

Finally, we used a t-test to compare average neonate dry mass within group C with maternal parasite exposure as a factor.

##### Section S1.2.2 Results

There was no significant difference in embryo production across the two maternal pathogen treatments (W=1452, p = 0.1722). However, not all of those embryos survived to become neonates, and embryo loss rate (that is, the difference between the number of embryos and neonates produced) was influenced by an interaction between maternal pathogen exposure and embryo care treatment (F_(1,70)_=22.991, p<0.001, Figure S1A). Specifically, when embryos were removed from the brood chamber, these embryos were less likely to survive to become neonates if they had been produced by mothers that were exposed to *O. pajunii*. Consistent with the prior two results (that is, no significant difference in embryo production and a significant exposure * embryo care treatment on survival to the neonate stage), neonate production varied with the interaction of maternal parasite and embryo care groups (F_(1,70)_=18.032, p<0.001, Figure S1B); pathogen-exposed moms whose embryos were removed from the brood chamber produced fewer neonates than all other mothers. Together, these results suggest that the embryos deposited by exposed mothers are less robust.

While the results suggest greater fragility of embryos produced by *O. pajunii*-exposed mothers, there were no detectable differences in embryo length (T_(21.402)_=-0.227, p=0.823), yolk width (W=83, p=0.519), yolk/length ratio (T_(20.252)_=-0.241, p=0.812), or embryo length/mom size ratio (W=97, p=1.00). There was also no difference detected in average neonate dry mass (T_(16.054)_=1.297, p=0.213). However, the within clutch variation in yolk size (W=31, p=0.004, Figure S1C) and embryo size (W=38, p=0.011, Figure S1D) was greater for *O. pajunii* exposed mothers than for unexposed mothers. Together, these results suggest that exposed mothers do not alter total reproductive investment, but instead may increase within-brood size variation as a strategy in the face of this pathogen challenge.

#### Section S1.3 Experimental test of potential for vertical transmission

We carried out an additional study to test for the possibility of vertical transmission of *O. pajunii*. One possibility is that compromised offspring from *O. pajunii*-exposed mothers may suffer reduced fitness due to vertically transmitted infection. While we have never observed visible symptoms of infection in the offspring of exposed mothers, we wanted to test whether there were very early infections that the embryos then successfully overcame (given that the offspring were never observed as having infections.) If this was the case, compromised offspring would display positive signals of *O. pajunii* infection via digital PCR (dPCR), even though they themselves had not been exposed to *O. pajunii*. We used dPCR because it is well-suited to quantifying even very low abundance targets.

**Figure S1:**
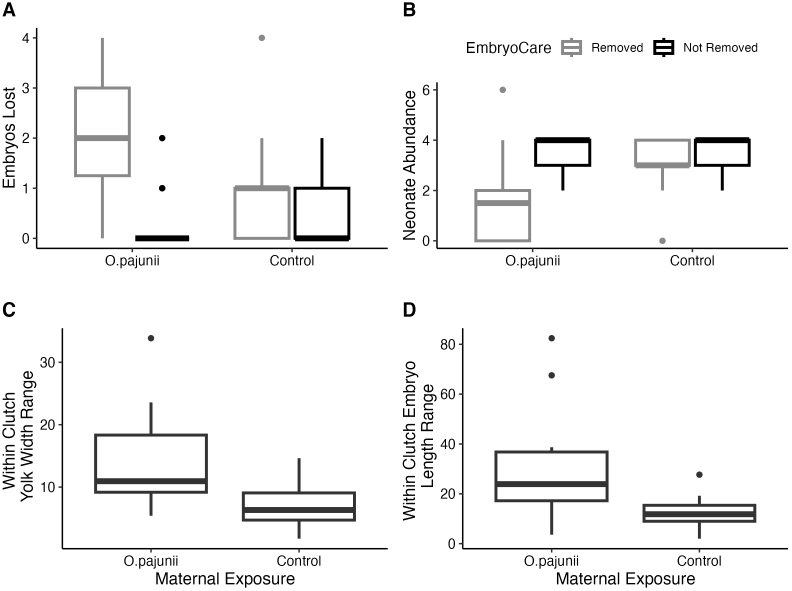
A: Embryo loss (the deficit between embryos produced and neonates surviving) was greater in *O. pajunii*-exposed mothers, only when embryos were removed from the maternal brood chamber. B: As a result, *O. pajunii-*exposed mothers produced fewer neonates, but only when embryos were removed from the brood chamber. C&D: The clutches produced by *O. pajunii*-exposed mothers had greater variation in yolk abundance provisioned to embryos (C) in embryo size (D).

For this experiment, S genotype *D. dentifera* grandmothers were acclimated in common conditions to minimize unintended variation within the mother generation. The mother generation was collected (on Day 1) as 24 hr old neonates, placed individually in 50 mL beakers with 20 mL filtered lake water. Mother beakers received 0.5 mL *A. falcatus* (AJT strain, 20,000 cells/mL) as food daily.

The pathogen exposure treatment consisted of a 48 hr exposure, beginning on Day 2, to a dose of *O. pajunii* spore slurry. We created the spore slurry by crushing individuals with mature (spore producing) infections into DI water in a ratio of 1 infected animal per experimental animal.

After this exposure ended, mothers were transferred through three parasite-free environments for 10 minutes each (to allow for processing and passing of any non-infecting spores remaining in the gut) and then were kept in 40 mL filtered lake water with food increased to 1 mL daily. Beginning on Day 5, animals were monitored daily for embryo production. On Day 7, when all mothers had produced embryos, embryos were extracted from mothers. One embryo from each *O. pajunii* exposed mother was chosen at random and placed into a 1.5 mL snap-top conical tube with 100 *µ*L ATL buffer (Qiagen) and stored for *O. pajunii* DNA quantification. Any positive results from this group would indicate potential vertical transmission during embryonic development in this system.

We extracted whole genomic DNA from each embryo using the DNeasy Blood and Tissue kit (Qiagen). PCRs were performed by the QIAcuity One dPCR system (Qiagen), in 24-well Nanoplates 26K (Qiagen). We used a probe-based assay that was designed and optimized for detection of *O. pajunii* (by Davenport et al. 2023). Temperature and timing conditions were 2 minutes at 95^*°*^C and 40 cycles of 15 seconds at 95^*°*^C and 1 minute at 55^*°*^C. Post-PCR imaging was performed using exposure timing of 300ms. This analysis included 1 known *O. pajunii*-infected animal (as positive control) and 1 blank (as negative control).

##### Section S1.3.1 Results

None of our 22 vertical transmission samples were positive for *O. pajunii* DNA. Given the absence of pathogen DNA in embryos of this treatment group, and that we have never observed a visible infection in the unexposed offspring of *O. pajunii*-exposed mothers, we conclude that *O. pajunii* is not transmitted vertically.

### Section S2 Details about model analysis

This section presents our mathematical models and our analytical and numerical results. Sections Section S2.1-Section S2.2 focus on a model where all infected individuals are identical, regardless of the infection status of their parent. Sections Section S2.3-Section S2.4 focus on a model where infected individuals are divided into infected individuals who are offspring of susceptible individuals (*I*) and those who are offspring of infected individuals (*D*). The results for the first model are more complete than the results for the second model because the lower dimensionality of the first model makes it more analytically tractable. The predictions from the two models are qualitatively similar.

The transgenerational impacts of infection that we consider here are all negative. Thus, as covered in the discussion in the main text, we consider these examples of transgenerational virulence. We use that term in the descriptions of the model in the supplement, as it is more succinct than transgenerational impacts of infection.

#### Section S2.1 SCIP model without interactions between transgenerational virulence and infection status

This model assumes all infected individuals are identical, regardless of the parent’s infection status. The SIR-type environmental transmission model describes the dynamics of the densities of susceptible individuals who are the offspring of susceptible individuals (*S*), infected individuals (*I*), susceptible individuals who are the offspring of infected individuals (hereafter, compromised individuals; *C*), and infectious propagules in the environment (*P*). Infection occurs when susceptible or compromised individuals encounter infectious propagules that were shed previously into the environment by infected individuals.

The equations for the SIC-form of the model are

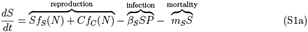

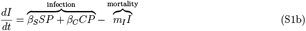

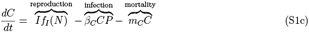

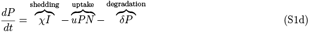

where *N* = *S* + *C* + *I*. In equation (S1a), susceptible individuals increase due to reproduction by susceptible and compromised individuals (*Sf*_*S*_(*N*) and *Cf*_*C*_(*N*), respectively) and decrease due to infection (*β*_*S*_*SP*) and non-disease mortality (*m*_*S*_*S*). In equation (S1b), infected individuals increase when susceptible and compromised become infected (*β*_*S*_*SP* and *β*_*C*_*CP*, respectively) and decrease due to mortality from disease and non-disease sources (*m*_*I*_*I*). In equation (S1c), compromised individuals increase due to reproduction by infected individuals (*If*_*I*_(*N*)) and decrease due to infection (*β*_*C*_*CP*) and mortality due to non-disease sources (*m*_*C*_*C*). In equation (S1d), infectious propagule density increases due to shedding by infected individuals (*χI*) and decreases due to uptake by all individuals (*uPN*) and degradation (*δ P*).

We assume the following in model (S1). First, the density-dependent per capita reproduction rates *f*_*i*_(*N*) (*i* ∈ {*S, C, I*}) are assumed to be decreasing functions of total density. Biologically, this means all individuals intraspecifically compete for resources and the competitive effects experienced by any individual only depend on total host density. Mathematically, we assume 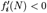 for *i* ∈ {*S, C, I*}. Second, we assume that all infected individuals are identical, regardless of whether they are the offspring of susceptible or infected individuals. Third, we assume the offspring of compromised individuals are identical to those of susceptible individuals (i.e., infection alters the fitness of the offspring of infected individuals, and not the grandchildren of infected individuals). Fourth, we assume the uptake rates for all individuals are equal (*u*). Fifth, when compared to susceptible individuals, we assume compromised individuals have (i) equal or higher mortality rates (*m*_*C*_ ≥ *m*_*S*_), (ii) equal or lower reproductive output (*f*_*C*_ ≤ *f*_*S*_), and (iii) higher, equal, or lower transmission rates (*β*_*C*_ > *β*_*S*_, *β*_*C*_ = *β*_*S*_, or *β*_*C*_ < *β*_*S*_, respectively). Sixth, we assume infected individuals have equal or lower reproductive output than susceptible individuals (*f*_*I*_ ≤ *f*_*S*_) and higher mortality rates (rates *m*_*I*_ > *m*_*S*_).

##### Section S2.1.1 Basic Reproduction Number

We derive the basic reproduction number, ℛ_0_, using the next generation matrix (van den Driessche and Watmough, 2008; Diekmann et al., 2010). The next generation matrix is constructed using the submatrix (𝒥) of the Jacobian corresponding to the equations for the infected class and the infectious propagules. Evaluating that submatrix at the disease-free equilibrium, (*N*, 0, 0, 0), yields

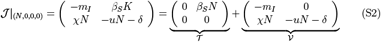

where 𝒯 is a matrix of transmission rates and 𝒱 is the matrix of transition rates. The basic reproduction number is

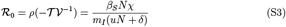

where *ρ*(*·*) is the spectral radius of a matrix. The basic reproduction number is the product of the average lifetime production of infectious propagules by an infected individual (*χ /m*_*I*_) and the average lifetime production of infected hosts by a spore in a complete susceptible host population (*β*_*S*_*N/*[*uN* + *δ*]).

##### Section S2.1.2 Frequency-form of the SCIP model

To facilitate the mathematical calculations, we write the model in terms of total host density (*N* = *S* + *C* + *I*), the fraction of susceptible individuals (*W* = *S/N*), the fraction of infected individuals (*Y* = *I/N*), and the fraction of compromised individuals (*X* = *C/N*). Because the frequencies sum to 1 (*W* + *Y* + *X* = 1), we only need equations for the frequencies of infected and compromised individuals and we compute the frequency of susceptible individuals using *W* = 1− *Y*− *X*.

The frequency-form of the model is

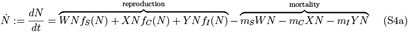

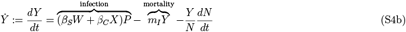

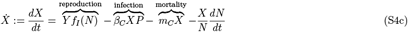

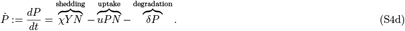

In equation (S4a), total host density increases due to reproduction by all host classes and decreases due to mortality of all host classes from disease and non-disease sources. In equation (S4b), the proportion of infected individuals increases when susceptible and compromised individuals become infected and decreases due to mortality from disease and non-disease sources. The final term in equation (S4b) follows from the quotient rule of calculus, i.e., *dY/dt* = *d*(*I/N*)*/dt* = (1*/N*)(*dI/dt*) − (*Y/N*)(*dN/dt*); that terms accounts for how the proportion of infected individuals changes if individuals are gained or lost from the population. In equation (S4c), the proportion of compromised individuals increases due to reproduction by infected individuals and decreases due to infection and mortality from non-disease sources. The last term in equation (S4c) follows from the quotient rule of calculus. Equation (S4d) is identical in structure to equation (S1d).

Our equilibrium-based analyses use the Jacobian of the frequency-form of the model (S4). When evaluated at a coexistence equilibrium point (*N*^∗^, *Y* ^∗^, *X*^∗^, *P*^∗^), the Jacobian has the structure

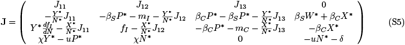

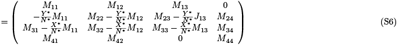

The terms *M*_*ij*_ are defined by the row reduced form of the Jacobian (**M**). The row reduced form of the Jacobian (**M**) and its sign structure are

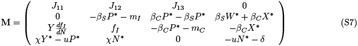

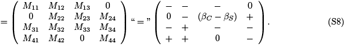

Explanations for the some signs of the entries are given below. Additional properties used in calculations are given in Proposition S1.

*M*_11_: Entry *M*_11_ defines the total amount of intraspecific competition experienced by the host species. The sign of this entry is always negative. To see this, note that the equilibrium condition for the *dN/dt* equation is 0 = (*W*^∗^*f*_*S*_ + *Y* ^∗^*f*_*I*_ + *X*^∗^*f*_*C*_ − *m*_*S*_*W*^∗^ − *m*_*C*_*X*^∗^ − *m*_*I*_*Y* ^∗^). Using this, *M*_11_ simplifies to

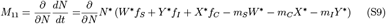

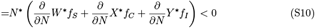

which is negative because each term in the final equation is negative by assumption.

*M*_12_: Entry *M*_12_ is always negative. It defines how the growth rate of the whole population is affected by reduced reproduction of infected individuals and disease induced mortality. Computing the derivative yields

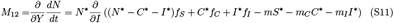

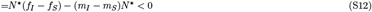

In the second line, the first term is non-positive because infected individuals are assumed to have equal or lower reproductive output than susceptible individuals (*f*_*I*_ ≤ *f*_*S*_) and the second term is negative because infected individuals are assumed to have greater mortality than susceptible individuals (*m*_*I*_ > *m*_*S*_).

*M*_13_: Entry *M*_13_ is always negative. It defines how the growth rate of the whole population is affected by reduced reproduction and increased mortality of compromised individuals. Computing the derivative yields

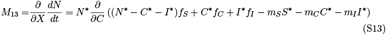

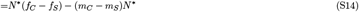

In the second line, the first term is non-positive because compromised individuals are assumed to have equal or less reproductive output than susceptible individuals (*f*_*C*_ ≤ *f*_*S*_) and the second term is non-positive because compromised individuals are assumed to have equal or greater mortality than susceptible individuals (*m*_*C*_ ≥ *m*_*S*_).

*M*_31_: Entry *M*_31_ = *Y* ^∗^*df*_*I*_ */dN* is negative and describes the effects of intraspecific competition from all classes on the reproduction rates of infected individuals.

*M*_41_: Entry *M*_41_ > 0 is the per capita production rate of infectious propagules (i.e., per capita shedding rate minus uptake rate). Entry *M*_41_ is positive because the equilibrium condition *dP/dt* = 0 implies 0 = *χY* ^∗^*N*^∗^ − *uP*^∗^*N*^∗^ − *δP*^∗^, which means *χY* ^∗^ −*uP*^∗^ = *δP*^∗^*/N*^∗^ > 0.

Proposition S1

For all of the systems considered in this study, the entries of the Jacobian (S7) satisfy the following inequalities:

1. *N*^∗^*M*_31_ − *M*_11_ > 0
2. *M*_24_*M*_33_ − *M*_23_*M*_34_ < 0
3. *M*_24_ + *M*_34_ > 0
4. *M*_22_*M*_44_ − *M*_24_*M*_42_ > 0
5. *M*_32_*M*_44_ − *M*_34_*M*_42_ < 0

*Reasoning for 1*

After canceling out terms, we get

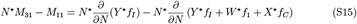

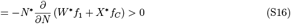

*Reasoning for 2*

After canceling out terms, we get

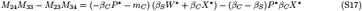

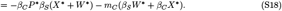

*Reasoning for 3*

It is straightforward to see that *M*_24_ + *M*_34_ = *β*_*S*_*W*^∗^ > 0.

*Reasoning for 4*

Rearranging the equilibrium condition 0 = *dP/dt* to get −*uN*^∗^ −*δ* = − *χY* ^∗^*N*^∗^*/P*^∗^, rearranging the equilibrium condition 0 = *dY/dt* to get *β*_*S*_*W*^∗^ + *β*_*C*_*X*^∗^ = *m*_*I*_*Y* ^∗^*/P*^∗^, and substituting yields

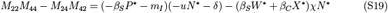

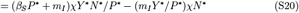

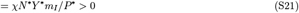

*Reasoning for 5*

Algebraic manipulation yields

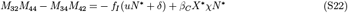

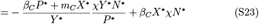

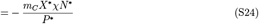

where the last line uses *dX/dt* = 0 to get *f*_*I*_ = (*β*_*C*_*X*^∗^*P*^∗^ + *m*_*C*_*X*^∗^)*/Y* ^∗^ and *dP/dt* = 0 to get

*uN*^∗^ + *δ* = *χY* ^∗^*N*^∗^*/P*^∗^.

##### Section S2.1.3 Density-form of the SCIP model

To facilitate the calculations of the sensitivities of infected density, we write the model in terms of total host density (*N* = *S* + *C* + *I*), susceptible density (*S*), infected density (*I*), and compromised individual density (*C*). Because the densities of the host classes sum to the total density (*N* = *S* + *I* + *C*), we only need equations for the densities of the infected and compromised individuals and we compute susceptible density using *S* = *N* – *I* − *C*.

The density-form of the model is

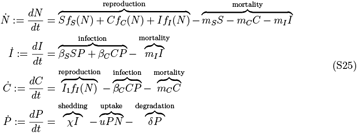

where *S* is evaluated as *S* = *N* −*I* − *C*.

When evaluated at a coexistence equilibrium (*N* ^∗^, *I* ^∗^, *C* ^∗^, *P* ^∗^), the form and structure of the Jacobian of the density-form of the model is

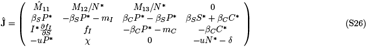

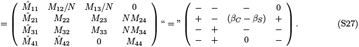

The signs of the *M*_*ij*_ are defined by the row reduced Jacobian (S7) of the frequency-form of the model (S4). The signs of the 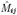 have known signs except for 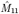.

We expect 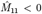 in most systems for the following reason. Entry 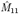 defines the intraspecific competitive effect of susceptible individuals on the reproduction rates of all individuals. (This is because varying *N* while holding *C* and *I* fixed is the same as varying *S*.) Computing the derivative yields

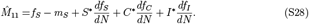

The first term is positive and the other terms are negative. We expect the sum to be negative because the opposite is only possible if the pathogen suppresses the host species to sufficiently low densities such that intraspecific competition is very weak.

##### Section S2.1.4 Translating predictions to direct transmission models

While our results focus on a model that assumes environmental transmission (ET), our results also apply to direct transmission models with density dependent direct transmission (DDDT), frequency dependent direct transmission (FDDT), or direct transmission is that intermediate to DDDT and FDDT. The reason is that in the limit where the infectious propagule dynamics are much faster than the dynamics of the host classes, the environmental transmission model (S1) reduces to model with direct transmission. This means that all of our equilibrium-based predictions for ET pathogens apply to apply DDDT and FDDT pathogens. Mathematical details are provided below; these results parallel those in Cortez and Duffy (2021) and Cortez (2022).

Assume the infectious propagule dynamics are much faster than the dynamics of the host classes. Mathematically, we assume *χ* is very large and *u* or *δ* are very large such that there is a separation of time scales between the infectious propagule equation and the host class equations. The quasi-steady state density for the infectious propagules is *P*^∗^ = *χI/*(*δ* + *uN*). Substitution into the equations for the host classes yields

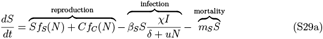

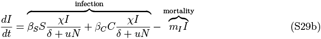

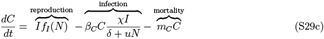

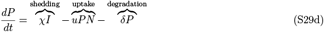

where 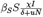 and 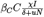 are the direct transmission rates from infected individuals to susceptible and compromised individuals, respectively. When *u* = 0 (i.e., uptake of infectious propagules is negligible relative to degradation), the direct transmission rates reduce to the DDDT forms, 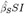 and 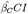, where 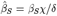 and 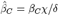. When *δ* = 0 (i.e., degradation of infectious propagules is negligible relative to uptake), the direct transmission rates reduce to the FDDT forms, 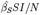 and 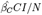, where 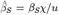 and 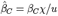. When *δ* and *u* are non-negligible, the transmission rates are intermediate to DDDT and FDDT.

Importantly, the above implies that all of the predictions for ET pathogens can be translated to predictions for DDDT and FDDT pathogens. Specifically, any conditions on *χ, u* and *δ* in the ET model can be interpreted in terms conditions on 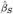 and 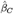 in a direct transmission model.

##### Section S2.1.5 Details about numerical simulations for the SCIP model

In simulations, the reproduction rates of the host classes were modeled using Lotka-Volterra functions. For those simulations, the reproduction rates are

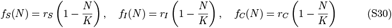

where *K* is the host carrying capacity.

To find examples, we manually searched parameter space using the analytical formulas to help identify likely regions were the desired phenomena would be located. We also searched parameter space by sampling parameters from uniform distributions with lower and upper bounds described below in the following order:

- The maximum reproduction rate for susceptible individuals (*r*_*S*_) and the transmission rates for susceptible (*β* _*S*_) and compromised (*β*_*C*_) individuals were sampled uniformly from the open interval (0,10).
- The maximum reproduction rates for compromised and susceptible individuals were sampled uniformly from the interval (0,*r*_*S*_), which ensures that compromised and infected individuals have lower growth rates than susceptible individuals.
- The host carrying capacity was uniformly sampled from the interval (100, 1,000).
- The mortality rate of susceptible individuals *m*_*S*_ was uniformly sampled from the interval (0, *r*_1_), which ensures that the host population would persist in the absence of the pathogen.
- The mortality rates of compromised and infected individuals were uniformly samples from the interval (*r*_1_,10), which ensures compromised and infected individuals have greater mortality rates than susceptible individuals.
- The uptake (*u*) and degradation (*δ*) rates were uniformly sampled from the interval (0,2). This smaller interval was chosen in order to increase the probability that the parameter set satisfied ℛ _0_ > 1.

#### Section S2.2 Analysis of SCIP model

We use local sensitivity analysis to explore how the equilibrium total host density (*N*^∗^), infection prevalence (*Y* ^∗^ = *I*^∗^*/N*^∗^), and infected density (*I*^∗^) depend on the characteristics of the compromised individuals. For a parameter *σ*, the (local) sensitivities of the equilibrium values to *σ* are computed using the derivatives *∂N*^∗^*/∂σ, ∂Y* ^∗^*/∂σ*, and *∂I*^∗^*/∂σ*. Positive values mean that increases in the parameter cause an increase in the equilibrium value whereas negative values mean that increases in the parameter cause a decrease in the equilibrium value. In the following, we compute the sensitivities to the mortality rate (*m*_*C*_), reproduction rate (*r*_*C*_), and transmission rate (*β*_*C*_) for compromised individuals. This analysis helps identify how increased mortality, reduced reproduction, and altered transmission rates for compromised individuals affects the host’s total density and disease levels at equilibrium.

Explicit formulas for the local sensitivities are computing via the Jacobian-based method used in Cortez and Duffy (2021) and Cortez (2022). In general, for a system of ordinary differential equations 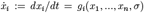 where *σ* is a parameter value,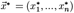 is an equilibrium point, and **J** is the Jacobian evaluated at 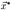, the sensitivity 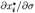 is

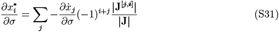

where **J**^[*j,i*]^ is the submatrix of **J** with row *j* and column *i* removed and |**J**| denotes the determinant of a matrix. To simplify the calculations of the sensitivities, we compute the sensitivities using the row reduced equations

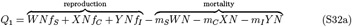

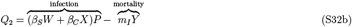

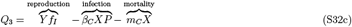

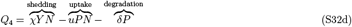

whose Jacobian is the matrix **M** in equation (S7). In total, we compute the sensitivities using the equations

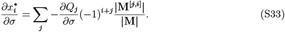

Note that |**M**| > 0 because properties of the determinant imply that |**M**| and |**J**| have the same sign.

##### Section S2.2.1 Sensitivities of equilibrium infection prevalence when total host density is constant

A useful special case to consider is when the total host population size is fixed (i.e., *N* fixed). This allows us to isolate how transgenerational virulence affects disease dynamics independent of its effects on host densities and feedbacks related to intraspecific competition. As a consequence of holding the population size fixed, the effects of transgenerational virulence on infected density and infection prevalence have the same sign and are proportional (because *Y* = *I/N*).

The model dynamics are determined by the *dY/dt, dX/dt* and *dP/dt* equations of frequency-form of the model (S5) where *dN/dt* = 0. The Jacobian of the model reduces to

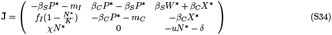

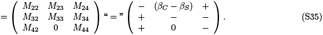

The last matrix shows the expected signs of the entries; explanations for the expected signs are given in Section Section S2. Note that sign 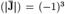 when evaluated at a stable equilibrium.

###### Sensitivity to mortality rate of compromised individuals, m_C_

Algebraic simplification yields

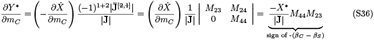

Based on the sign of the term, infection prevalence and infected density decrease if compromised individuals have larger transmission rates than susceptible individuals (*β*_*C*_ > *β*_*S*_). Infection prevalence and infected density increase if compromised individuals have smaller transmission rates than susceptible individuals (*β*_*C*_ < *β*_*S*_).

###### Sensitivity to transmission rate of compromised individuals, *β*_**C**_

Algebraic simplification yields

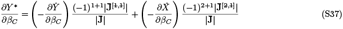

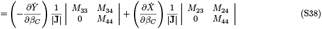

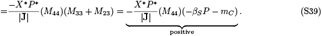

Based on the sign of the term, infection prevalence and infected density increase as the transmission rate of compromised individuals increases.

##### Section S2.2.2 Sensitivities of equilibrium total host density

For a given parameter, *σ*, the sensitivity of the total host population at equilibrium (*N*^∗^) to *σ* is

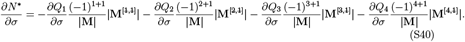

Below, we compute the sensitivities of total host density to *m*_*C*_, *r*_*C*_, and *β*_*C*_.

###### Sensitivity of total host density to the mortality rate of compromised individuals, *m*_*C;*_

Algebraic simplification yields

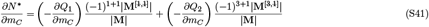

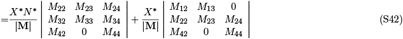

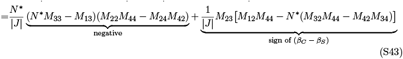

In the last equation, the terms in brackets are positive because of Property S1. Based on these signs, we predict that *∂N*^∗^*/∂m*_*C*_ is negative, unless susceptible individuals have sufficiently smaller transmission rates than compromised individuals (*β*_*S*_ < *β*_*C*_).

###### Sensitivity of total host density to the reproduction rate of compromised individuals, *r*_*C:*_

Let *r*_*C*_ be a parameter satisfying *∂f*_*C*_*/∂r*_*C*_ > 0 and *∂f*_*i*_*/∂r*_*C*_ = 0 for *i* = *S* or *i* = *I*. This means *∂N*? */∂r*_*C*_ = *XN∂f*_*C*_*/∂r*_*C*_ > 0. Algebraic simplification yields

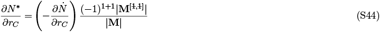

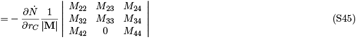

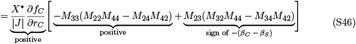

The signs of the terms in the last equation follow from Property S1. Based on these signs, we predict that *∂N*^∗^*/∂m*_*C*_ is positive, unless susceptible individuals have sufficiently smaller transmission rates than compromised individuals (*β*_*S*_ < *β*_*C*_).

###### Sensitivity of total host density to the transmission rate of compromised individuals, *β*_*C*_

Algebraic manipulation yields

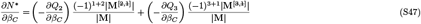

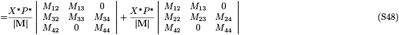

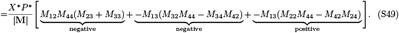

The signs of the terms follow from Property S1. The positive term is larger in magnitude when infected individuals have very large mortality rates and very low reproduction rates relative to susceptible individuals (*M*_13_ large in magnitude) infected individuals have large shedding rate (*M*_42_ large in magnitude). Based on the above, we predict that *∂N*^∗^*/∂β*_*C*_ is negative, unless infected individuals very large mortality rates or very low reproduction rates and infected individuals have large shedding rate.

##### Section S2.2.3 Sensitivities of equilibrium infection prevalence

For a given parameter *σ*, the sensitivity of infection prevalence at equilibrium (*Y* ^∗^) to *σ* is

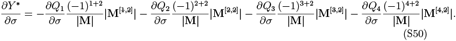

Below, we compute the sensitivities of infection prevalence to *m*_*C*_, *r*_*C*_, and *β*_*C*_.

###### Sensitivity of infection prevalence to the mortality rate of compromised individuals, **m**_**C**_

Algebraic simplification yields

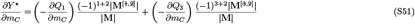

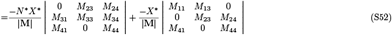

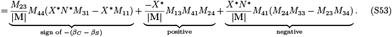

The first term has the same sign as − (*β*_*C*_ − *β*_*S*_) because of Property S1; the second term is positive; and the third term is negative because of Property S1. The second term is large in magnitude when *M*_13_ is large in magnitude, which occurs when compromised individuals have very high mortality rates (*m*_*C*_). Based on this, we predict that *∂Y* ^∗^*/∂m*_*C*_ is negative, unless (i) susceptible individuals have sufficiently larger transmission rates than compromised individuals (*β*_*S*_ > *β*_*C*_) or (ii) compromised individuals have very high mortality rates.

###### Sensitivity of infection prevalence to the reproduction rate of compromised individuals, *r*_*C*_

Let *r*_*C*_ be a parameter satisfying *∂f*_*C*_*/∂r*_*C*_ > 0 and *∂f*_*i*_*/∂r*_*C*_ = 0 for *i* = *S* or *i* = *I*. This means 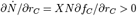. Algebraic simplification yields

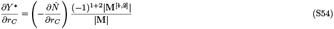

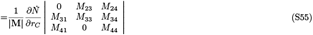

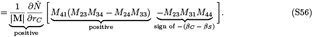

The signs of the terms follow from Property S1. Based on the above, we predict that *∂Y* ^∗^*/∂r*_*C*_ is positive, unless compromised individuals have sufficiently larger transmission rates than susceptible individuals (*β*_*C*_ > *β*_*S*_).

###### Sensitivity of infection prevalence to the transmission rate of compromised individuals, *β*_*C*_

Algebraic simplification yields

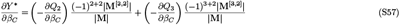

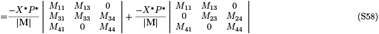

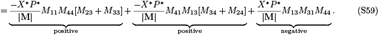

The first term is positive because *M*_23_ + *M*_33_ = −*β*_*S*_*P*^∗^ − *m*_*C*_. The sign of the second term follows from Proposition S1. The third term is large in magnitude when infected individuals have very large mortality rates and compromised individuals have much larger mortality rates than susceptible individuals (*m*_*C*_ ≫ *m*_*S*_; making *M*_13_ large in magnitude). Based on this, we expect *∂Y* ^∗^*/∂β*_*C*_ is positive unless infected individuals and compromised individuals have very large mortality rates relative to susceptible individuals.

##### Section S2.2.4 Sensitivities of equilibrium infected density

We compute the sensitivities of infected density in two ways. First, we use the formula

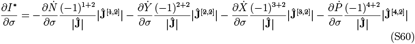

where **Ĵ** is the Jacobian of the density-form of the model (S1), defined in equation (S26). Second, using the quotient rule from calculus, the sensitivities of infected density and infection prevalence are related by

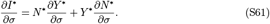

The advantage of the first approach is that it allows us to directly compute the sensitivity of infected density. The advantage of the second approach is that it allows us to determine the conditions under which the sensitivities of infected density and infection prevalence have the same or opposite signs. In particular, the sensitivities of infected density 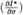 and infection prevalence 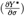 have the same sign unless the sensitivity for total host density 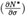 is large in magnitude and of the opposite sign of the sensitivity for infection prevalence. Biologically, this means that infected density and infection prevalence have responses of opposite sign only if there are large changes in host density (e.g., infection prevalence increases but infected density decreases because total host population size decreases).

###### Sensitivity of infected density to the mortality rate of compromised individuals, **m**_**C**_

Algebraic simplification yields

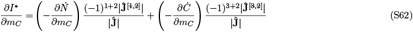

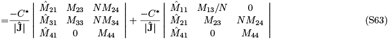

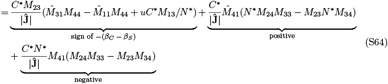

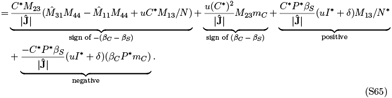

The sign of the first term follows from the expectation that 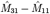 is positive. Overall, the sign of *∂I*^∗^*/∂m*_*C*_ is difficult to determine from the above equation.

Using the formulas for *∂Y* ^∗^*/∂m*_*C*_ and *∂N*^∗^*/∂m*_*C*_ in Section Section S2.2.3, the sensitivity of infected density can be written was

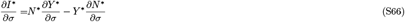

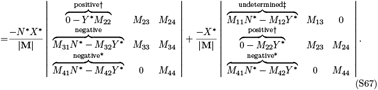

We focus on the signs of the entries in the first column of each submatrix. The sensitivities *∂I*^∗^*/∂m*_*C*_ and *∂Y* ^∗^*/∂m*_*C*_ can have opposite signs when the two terms in each entry have different signs and the second term is larger in magnitude than the first. For the two entries denoted with asterisks (∗), the first term is positive, the second term is negative, and the second term is larger in magnitude than the first when infected individuals have large shedding rate (*χ* large). For the two entries denoted with daggers (†), the second term is positive and large in magnitude when infected individuals have large mortality rates (*m*_*I*_). For the entry denoted with a double dagger (‡), the first term is negative, the second term is positive, and the second term is larger in magnitude when infected individuals have low reproduction rates (*f*_*I*_ small) and disease-induced mortality is large (*m*_*I*_ large).

Based on the above, *∂I*^∗^*/∂m*_*C*_ and *∂Y* ^∗^*/∂m*_*C*_ often have the same sign, implying that *∂I*^∗^*/∂m*_*C*_ < 0 unless (i) susceptible individuals have sufficiently larger transmission rates than compromised individuals (*β*_*S*_ ≫ *β*_*C*_) or (ii) compromised individuals have very high mortality rates. However, *∂I*^∗^*/∂m*_*C*_ and *∂Y* ^∗^*/∂m*_*C*_ can have opposite signs when infected individuals have sufficiently large mortality rates (*m*_*I*_), sufficiently large shedding rate (*χ*), or sufficiently low reproduction rates (*f*_*I*_).

###### Sensitivity of infected density to the reproduction rate of compromised individuals, **r**_**C**_

Let *r*_*C*_ be a parameter satisfying *∂f*_*C*_*/∂r*_*C*_ > 0 and *∂f*_*i*_*/∂r*_*C*_ = 0 for *i* = *S* or *i* = *I*. This means 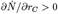. Algebraic simplification yields

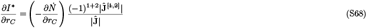

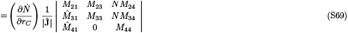

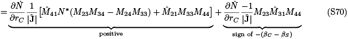

Based on the above, we predict that *∂I*^∗^*/∂r*_*C*_ is positive unless compromised individuals have sufficiently higher transmission rates than susceptible individuals (*β*_*C*_ > *β*_*S*_).

Using the formulas for *∂Y* ^∗^*/∂r*_*C*_ and *∂N*^∗^*/∂r*_*C*_ in Section Section S2.2.3, the sensitivity We focus on the sign of the entries in the first column of the matrix. The sensitivities

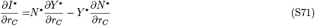

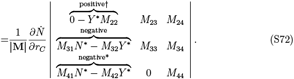

*∂Y* ^∗^*/∂r*_*C*_ and *∂N*^∗^*/∂r*_*C*_ can have opposite signs when the two terms in each entry have different signs and the second term is larger in magnitude than the first. This is possible for the entries denoted with asterisks (*) when infected individuals have sufficiently large mortality (*m*_*I*_) or shedding (*χ*) rates.

In total, equation (S67) shows that *∂I/∂r*_*C*_ is positive unless compromised individuals have sufficiently higher transmission rates than susceptible individuals (*β*_*C*_ > *β*_*S*_). Equation (S72) shows that *∂Y* ^∗^*/∂r*_*C*_ and *∂N*^∗^*/∂r*_*C*_ often have the same sign, but they can have opposite signs when infected individuals have sufficiently large mortality (*m*_*I*_) or shedding (*χ*) rates.

###### Sensitivity of infected density to the transmission rate of compromised individuals, *β*_**C**_

Algebraic simplification yields

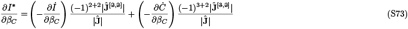

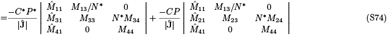

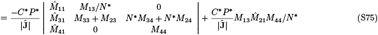

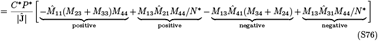

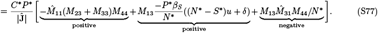

Based on the above, we expect *∂I*^∗^*/∂β*_*C*_ to be positive unless the reproduction rates of infected individuals are strongly affected by competition (making *M*_31_ large in magnitude) and compromised individuals have much large mortality rates than susceptible individuals (making *M*_13_ large in magnitude).

Using the formulas for *∂Y* ^∗^*/∂β*_*C*_ and *∂N*^∗^*/∂β*_*C*_ in Section Section S2.2.3, the sensitivity

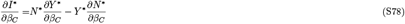

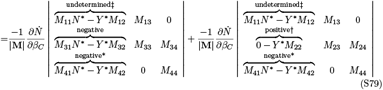

We focus on the sign of the entries in the first column of the each matrix. The sensitivities *∂Y* ^∗^*/∂β*_*C*_ and *∂N*^∗^*/∂β*_*C*_ can have opposite signs when the two terms in each entry have different signs and the second term is larger in magnitude than the first. For the two entries denoted with asterisks (*), the first term is positive, the second term is negative, and the second term is larger in magnitude than the first when infected individuals have large shedding rate (*χ* large). For the entry denoted with a dagger (†), the second term is positive and larger in magnitude when infected individuals have large mortality rates (*m*_*I*_). For the two entries denoted with double daggers (‡), the first term is negative, the second term is positive, and the second term is large in magnitude when infected individuals have low reproduction rates (*f*_*I*_ small) and disease-induced mortality is large (*m*_*I*_ large).

In total, equation (S77) shows that *∂I/∂β*_*C*_ is positive unless the reproduction rates of infected individuals are strongly affected by competition and compromised individuals have much larger mortality rates than susceptible individuals (*m*_*C*_ ≫ *m*_*S*_). Equation (S79) shows that *∂Y* ^∗^*/∂r*_*C*_ and *∂N*^∗^*/∂r*_*C*_ often have the same sign, but they can have opposite signs when infected individuals have sufficiently large mortality rates (*m*_*I*_), sufficiently large shedding rate (*χ*), or sufficiently low reproduction rates (*f*_*I*_).

##### Section S2.2.5 Summary and interpretation of sensitivity results

The expected signs of the sensitivities and conditions under which the sensitivities have the opposite signs are shown in Table S2. In Figures S2-S4, examples where the sensitivities have the expected signs are plotted in bluer hues and examples with the opposite signs are plotted in redder hues. The large dots indicate the values for systems where transgenerational virulence is absent (i.e., compromised individuals are identical to susceptible individuals).

###### Total host density

We predict that in many systems, total host density decreases with increased mortality, reduced reproduction, and increased transmission rates. However, counterintuitively, the predictions can be reversed if compromised individuals have higher mortality rates, lower reproduction rates, and much higher transmission rates than susceptible individuals. Our specific predictions are the following: (1) Increased mortality rates of compromised individuals (*m*_*C*_) decreases total host density (bluer curves in Figure S2A) unless susceptible individuals have sufficiently smaller transmission rates than compromised individuals (*β*_*S*_ < *β*_*C*_; redder curves in Figure S2A). (2) Decreased reproduction rates of compromised individuals (*r*_*C*_) decreases total host density (all curves Figure S2B) unless susceptible individuals have sufficiently smaller transmission rates than compromised individuals (*β*_*S*_ < *β*_*C*_). However, we were unable to find any examples of increasing total density using our model with Lotka-Volterra competition (hence why all curves are increasing in Figure S2B). (3)

Increased compromised infection rates (*β*_*C*_) decreases total host density (bluer curves in Figure S2C) unless infected individuals have large shedding rates (*χ*) and much larger mortality rates or much lower reproduction rates than susceptible individuals (*m*_*I*_ ≫ *m*_*S*_ and *f*_*I*_ ≪ *f*_*S*_, respectively) (redder curves in Figure S2C).

The counter-intuitive responses in total host density allow for the possibility that total host density is higher with transgenerational virulence than not. However, we were unable to find any numerical examples where total host density was higher with transgenerational virulence. We predict that higher total density with transgenerational virulence will be exceeding rare, if not impossible, because the above conditions for the counter-intuitive responses cause large decreases total host density (i.e., higher mortality, lower reproduction, and higher transmission for compromised individuals cause large decreases in total host density).

###### Infection prevalence

We predict that increased mortality, reduced reproduction, and decreased transmission rates of compromised individuals reduces infection prevalence. However, counter-intuitively, these predictions can be reversed if susceptible individuals have much larger transmission rates than compromised individuals or compromised individuals have very high mortality rates. Our specific predictions are the following: (1) Increased mortality rates of compromised individuals decreases infection prevalence (bluer curves in Figure S3A) unless susceptible individuals have sufficiently larger transmission rates than compromised individuals (*β*_*S*_ > *β*_*C*_) or compromised individuals have very high mortality rates (*m*_*C*_ ≫ *m*_*S*_) (redder curves in Figure S3A). (2) Decreased reproduction rates of compromised individuals decreases infection prevalence (bluer curves in Figure S3B) unless susceptible individuals have sufficiently smaller transmission rates than compromised individuals (*β*_*S*_ < *β*_*C*_; redder curves in Figure S3B). (3) Increased compromised infection rates increases infection prevalence (all curves in Figure S3C) unless compromised and infected individuals have very large mortality rates relative to susceptible individuals (*m*_*C*_, *m*_*I*_ ≫ *m*_*S*_). We where unable to find any examples of the latter (hence, why all curves are blue in Figure S3C).

The counter-intuitive responses in total host density allow for the possibility that infection prevalence is higher with could be higher with transgenerational virulence than not. We were only able to find such examples when compromised individuals had much higher transmission rates than susceptible individuals (*β*_*C*_ ≫ *β*_*S*_). We predict that greater infection prevalence with transgenerational virulence will be exceeding rare, if not impossible, whenever this condition is not met (i.e, when compromised individuals have lower transmission rates than susceptible individuals). The reason is that the conditions for the counter-intuitive responses in infection prevalence cause large decreases in infection prevalence (e.g., low compromised reproduction rates and higher compromised mortality rates cause large decreases in infection prevalence).

###### Infected density

The predictions for infected density are qualitatively similar to the predictions for infection prevalence. However, the signs of the responses in infected density and infection prevalence can differ when infected individuals have sufficiently high mortality rates (*m*_*I*_), sufficiently high shedding rates (*χ*), or sufficiently low reproduction rates (*f*_*I*_). One difference we found was that decreased reproduction rates always decreased infected density (all curves blue in Figure S4B). As with infection prevalence, our predictions allow for the possibility that infected density could be higher when compromised individuals have much higher mortality rates, lower reproduction rates, and much lower transmission rates than susceptible individuals. However, we were unable to find any numerical examples where those conditions resulted in higher infected density due to transgenerational virulence.

###### Direct transmission models

All of the above predictions directly translate to predictions for direct transmission pathogens with density-dependent direct transmission, frequencydependent direct transmission, or transmission that is intermediate; see Section Section S2.1.4 for mathematical details. The only differences are that (i) conditions about large and small shedding rates (*χ*_*I*_) should be interpreted in terms of large and small direct transmission parameters (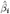for *i ∈* {*S, C*}) and (ii) conditions about large and small infection parameters (*β*_*i*_ for *i* ∈ {*S, C*}) in the environmental transmission model should be interpreted in terms of large and small direct transmission parameters (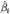 for *i* ∈ {*S, C*}) in the direct transmission models.

**Table S2:** Predicted local sensitivities to parameters of compromised individuals for the SCIP model (S1)

| Sensitivity | Expected Sign | Exceptions |
| --- | --- | --- |
| <u>Total density</u> |  |  |
| $\frac{\partial N^*}{\partial m_C}$ | – | susceptible individuals have sufficiently smaller transmission rates than compromised individuals ($\beta_S < \beta_C$) |
| $\frac{\partial N^*}{\partial r_C}$ | + | susceptible individuals have sufficiently smaller transmission rates than compromised individuals ($\beta_S < \beta_C$) |
| $\frac{\partial N^*}{\partial \beta_C}$ | – | infected individuals have large shedding rates ($\chi$ large)<br>and larger mortality rates or smaller reproduction rates than susceptible individuals ($m_I \gg m_S, f_I \ll f_S$) |
| <u>Infection Prevalence</u> |  |  |
| $\frac{\partial Y^*}{\partial m_C}$ | – | susceptible individuals have sufficiently larger transmission rates than compromised individuals ($\beta_S > \beta_C$)<br>or compromised individuals have very high mortality rates ($m_C \gg m_S$) |
| $\frac{\partial Y^*}{\partial r_C}$ | + | susceptible individuals have sufficiently smaller transmission rates than compromised individuals ($\beta_S < \beta_C$) |
| $\frac{\partial Y^*}{\partial \beta_C}$ | + | infected and compromised individuals have much larger mortality rates than susceptible individuals ($m_I, m_C \gg m_S$) |
| <u>Infected Density†</u> |  |  |
| $\frac{\partial I^*}{\partial m_C}$ | – | susceptible individuals have sufficiently larger transmission rates than compromised individuals ($\beta_S > \beta_C$)<br>or compromised individuals have very high mortality rates ($m_C \gg m_S$) |
| $\frac{\partial I^*}{\partial r_C}$ | + | susceptible individuals have sufficiently smaller transmission rates than compromised individuals ($\beta_S < \beta_C$) |
| $\frac{\partial I^*}{\partial \beta_C}$ | + | infected and compromised individuals have much larger mortality rates than susceptible individuals ($m_I, m_C \gg m_S$) |

†The sensitivities for infection prevalence and infected density often have the same sign, but the sensitivities for infected density can have the opposite sign when infected individuals have sufficiently high mortality rates ( $m_I$ large), sufficiently high shedding rates ( $\chi$ large), or sufficiently low reproduction rates ( $f_I$ small).

**Figure S2:**
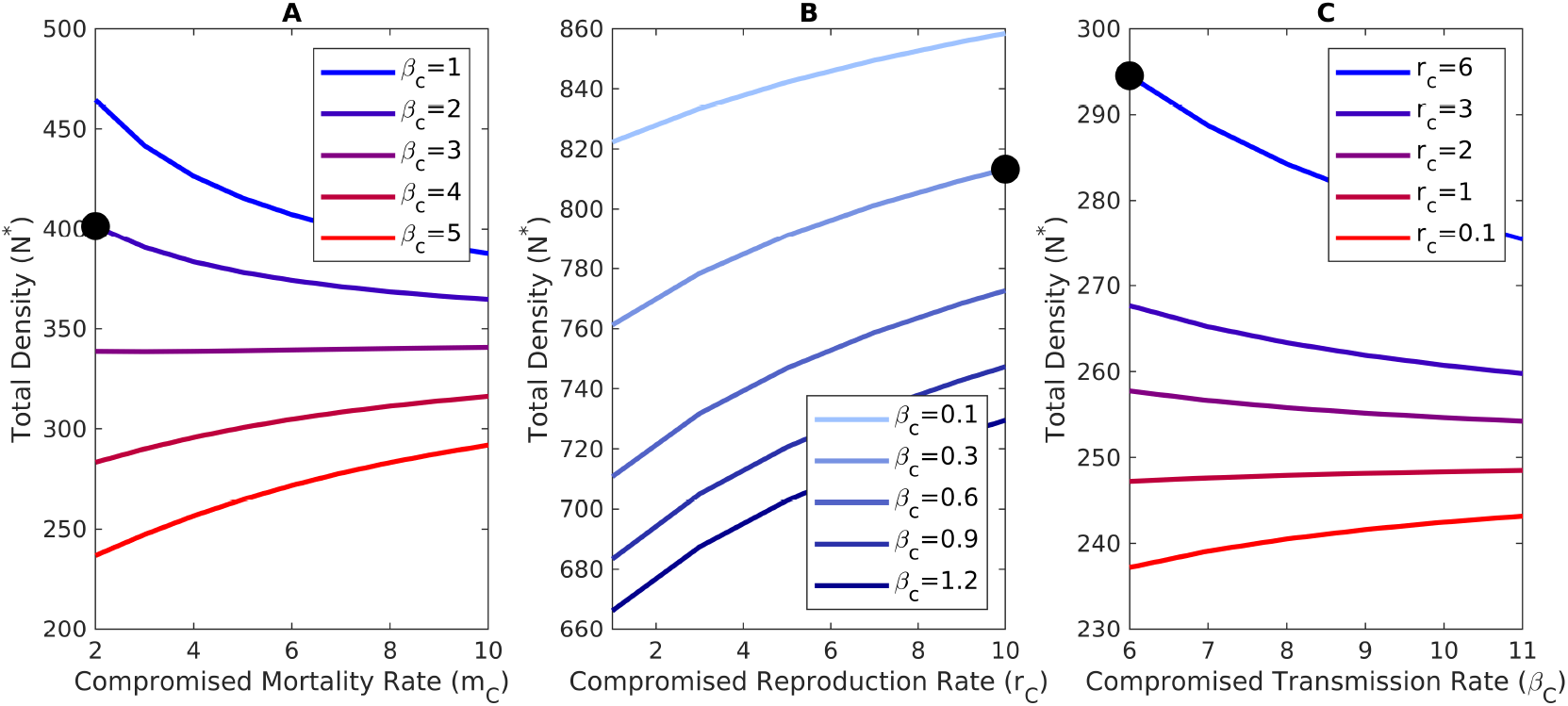
Responses of total density to the effects of transgenerational virulence. In each panel, bluer curves illustrate the responses expected in most systems and redder curves denote responses of the opposite sign. Black circles denote values when there are no transgenerational virulent effects. (A) Total density decreases with increased mortality of compromised individuals (redder curves), unless compromised individuals have sufficiently high transmission rates (bluer curves). (B) Total density increases with increased reproduction of compromised individuals (*r*_*C*_ in a model with Lotka-Volterra competition; all curves). In principle the opposite response can occur if compromised individuals have sufficiently high transmission rates, but we did not find any numerical examples. (C) Total density decreases with increase transmission rate of compromised individuals (bluer curves), unless infected individuals experience sufficiently strong intraspecific competition (which can occur in our model when compromised individuals have lower reproduction rates, *r*_*c*_; redder curves). The parameter values are (A) *r*_*S*_ = 6, *r*_*C*_ = 6, *r*_*I*_ = 3, *K* = 1000, *m*_*S*_ = 2, *m*_*I*_ = 10, *β*_*S*_ = 2, *χ* = 6, *u* = 1, *δ* = 2; (B) *r*_*S*_ = 10, *r*_*I*_ = 4, *K* = 1000, *m*_*S*_ = 0.01, *m*_*I*_ = 2.25, *m*_*C*_ = 0.75, *β*_*S*_ = 0.3, *χ* = 0.44, *u* = 0.277, *δ* = 0.25; and (C) *r*_*S*_ = 6, *r*_*I*_ = 4, *K* = 1000, *m*_*S*_ = 1.5,*m*_*I*_ = 3, *m*_*c*_ = 1, *β*_*S*_ = 6, *χ* = 9.5, *u* = 1, *δ* = 1.

**Figure S3:**
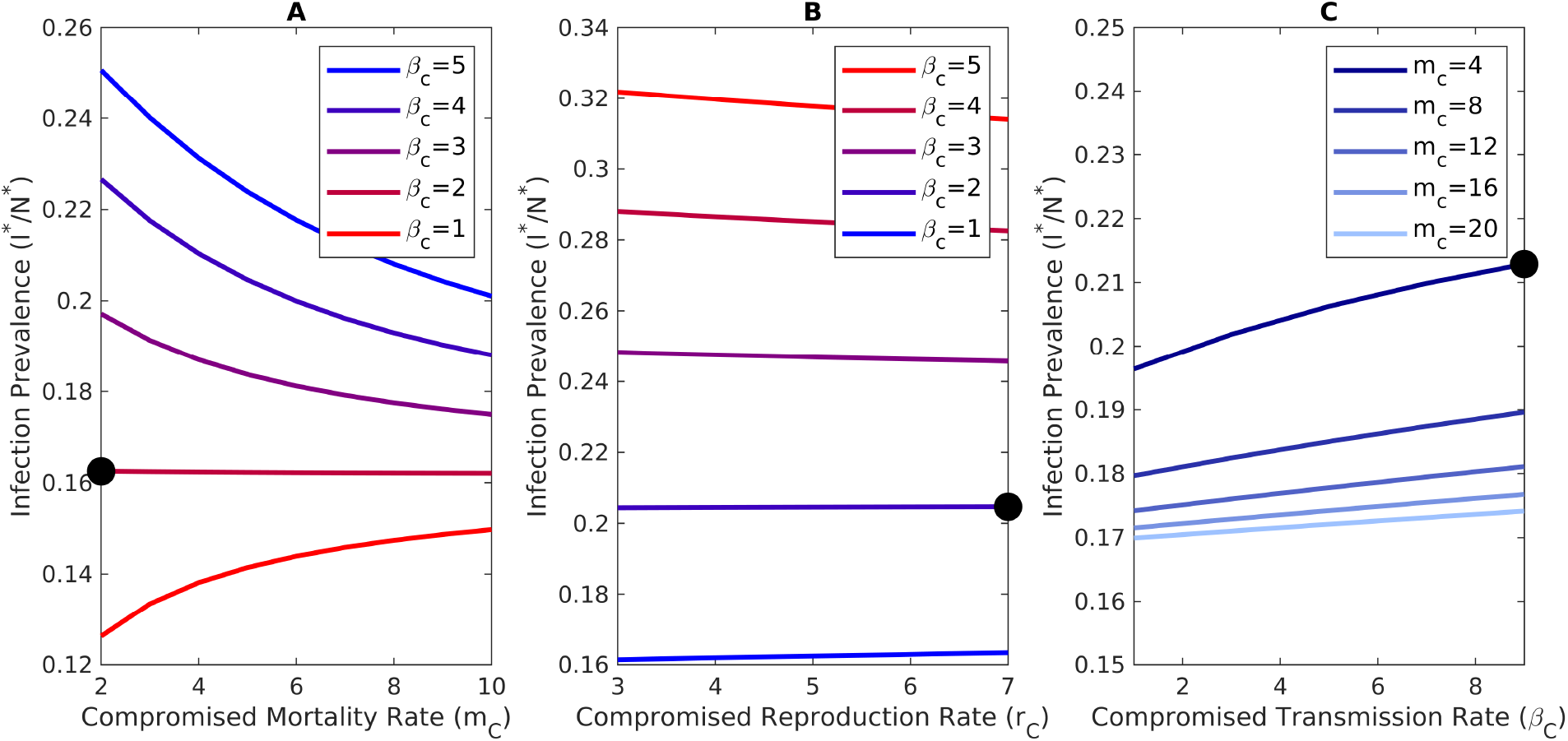
Responses of infection prevalence to the effects of transgenerational virulence. In each panel, bluer curves illustrate the responses expected in most systems and redder curves denote responses of the opposite sign. Black circles denote values when there are no transgenerational virulent effects. (A) Total prevalence decreases with increased mortality of compromised individuals (bluer curves), unless compromised individuals have sufficiently high transmission rates (redder curves). (B) Total prevalence increases with increased reproduction of compromised individuals (*r*_*C*_ in a model with Lotka-Volterra competition; bluer curves), unless compromised individuals have sufficiently high transmission rates (red). (C) Total prevalence increases with increase transmission rate of compromised individuals (all curves). In principle the opposite response can occur if infected individuals experience strong intraspecific competition and compromised individuals have much larger mortality rates than susceptible individuals, but we did not find any numerical examples. The parameter values are (A) *r*_*S*_ = 6, *r*_*C*_ = 6, *r*_*I*_ = 3, *K* = 1000, *m*_*S*_ = 2, *m*_*I*_ = 10, *β*_*S*_ = 2, *χ* = 6, *u* = 1, *δ* = 2; (B) *r*_*S*_ = 7, *r*_*I*_ = 4, *K* = 1000, *m*_*S*_ = 2, *m*_*I*_ = 9.5, *m*_*C*_ = 3, *β*_*S*_ = 2, *χ* = 6, *u* = 1, *δ* = 2; and (C) *r*_*S*_ = 6, *r*_*C*_ = 5, *r*_*I*_ = 3, *K* = 10, *m*_*S*_ = 4, *m*_*I*_ = 8, *β*_*S*_ = 9, *χ* = 2, *u* = 0.5, *δ* = 1.

**Figure S4:**
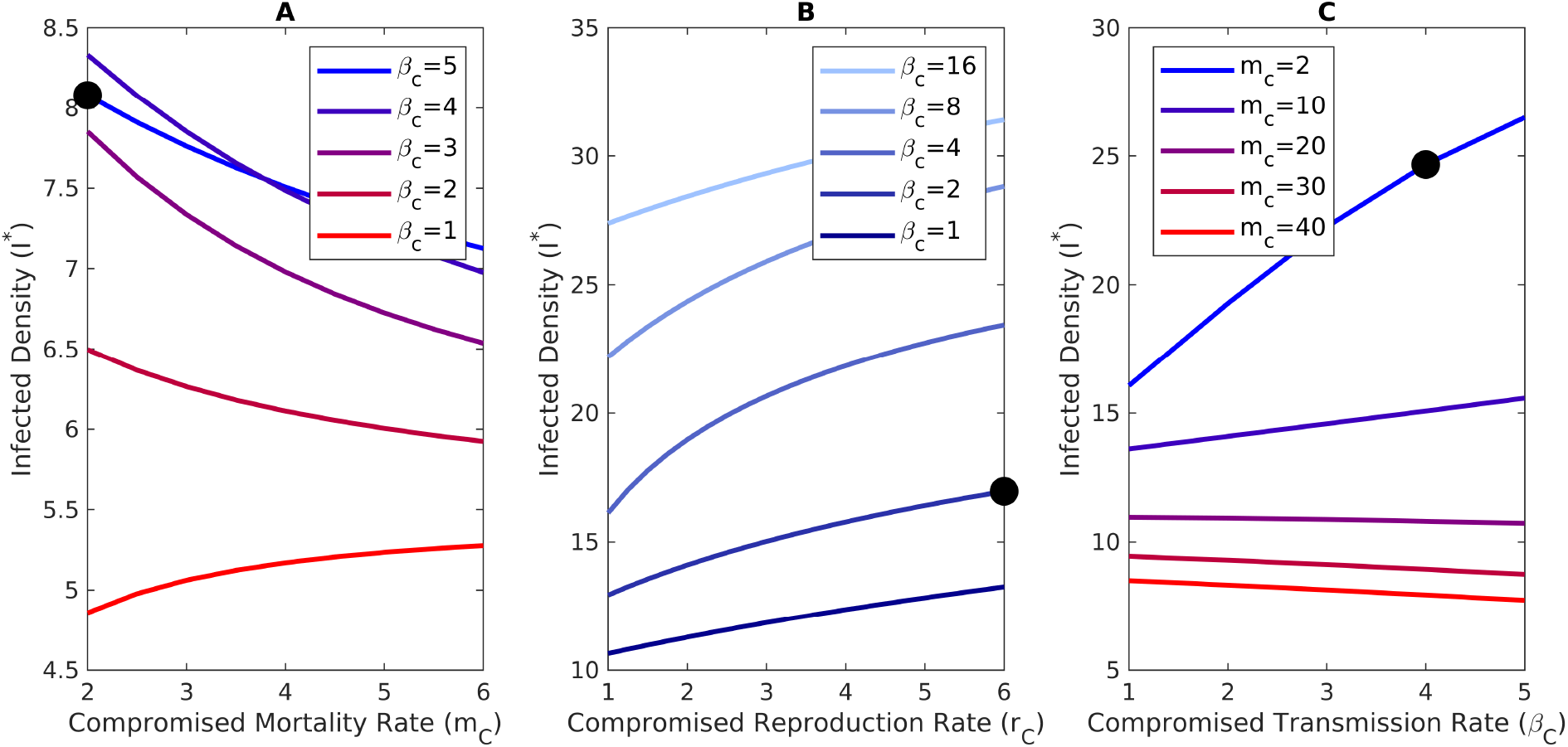
Responses of infected density to the effects of transgenerational virulence. In each panel, bluer curves illustrate the responses expected in most systems and redder curves denote responses of the opposite sign. Black circles denote values when there are no transgenerational virulent effects. (A) Infected density decreases with increased mortality of compromised individuals (bluer curves), unless compromised individuals have sufficiently high transmission rates (redder curves). (B) Infected density decreases with decreased reproduction of compromised individuals (*r*_*C*_ in a model with Lotka-Volterra competition; all curves). In principle the opposite response can occur if compromised individuals have sufficiently high transmission rates, but we did not find any numerical examples. (C) Infected density increases with increase transmission rate of compromised individuals (bluer curves), unless infected individuals are strongly affected by competition and compromised individuals have sufficiently large mortality rates (redder curves). The parameter values are (A) *r*_*S*_ = 6, *r*_*C*_ = 6, *r*_*I*_ = 6, *K* = 100, *m*_*S*_ = 2, *m*_*I*_ = 10, *β*_*S*_ = 2, *χ* = 6, *u* = 1, *δ* = 2; (B) *r*_*S*_ = 6, *r*_*I*_ = 6, *K* = 100, *m*_*S*_ = 2, *m*_*I*_ = 4, *m*_*C*_ = 3, *β*_*S*_ = 2, *χ* = 3, *u* = 1, *δ* = 1; and (C) *r*_*S*_ = 6, *r*_*C*_ = 6, *r*_*I*_ = 6, *K* = 100, *m*_*S*_ = 2, *m*_*I*_ = 4, *β*_*S*_ = 4, *χ* = 3, *u* = 1.5, *δ* = 1.5.

##### Section S2.2.6 Conditions for bistability

Transgenerational virulence can lead to bistability where model (S1) has (i) a locally asymptotically stable disease-free equilibrium, (ii) a locally asymptotically stable endemic equilibrium, and (iii) a saddle-type endemic equilibrium. Our numerical and analytical results (see below) suggest that this occurs when *R*_0_ is less than 1 by a small amount and compromised individuals have much larger transmission rates than susceptible individuals (*β*_*C*_ ≫ *β*_*S*_).

This is important biologically because it shows that transgenerational virulence can allow diseases to persist in systems where they cannot invade from low densities. As a particular example, consider a species of lake *Daphnia* and its environmentally transmitted parasite. In a bistable system, disease outbreaks cannot occur if only small amounts of infectious propagules are introduced, e.g., by migrating animals, human activities, or a small stream. However, if an upwelling event releases a large density of infectious propagules that was trapped in the sediment at the bottom of a lake, then an outbreak could occur. Thus, transgenerational virulence that increases offspring’s susceptibility to disease could allow for outbreaks in systems where outbreaks would otherwise not be possible.

In our numerical simulations of model (S4) with Lotka-Volterra competition, we found bistability when ℛ_0_ was less than 1 by a small amount and compromised individuals had much larger transmission rates than susceptible individuals (*β*_*C*_ *≫ β*_*S*_); see figure S5 for a particular example. We were unable to show that ℛ_0_ < 1 and *β*_*C*_ > *β*_*S*_ are necessary conditions for bistability in model (S4). However, we were able to show those conditions are necessary for the coexistence of multiple endemic equilibria in the special case where the host population density is held constant (i.e., *N* is fixed). In particular, if model (S4) has fixed total host density and all individuals have equal competitive ability (e.g., see equation (S30)), then the following shows that the model has two endemic equilibria only when ℛ_0_ < 1 and *β*_*C*_ > *β*_*S*_. Combined, the analytical result and our numerical simulations suggest that transgenerational virulence can lead to bistability, but it is only likely to do so when compromised individuals have much larger transmission rates than susceptible individuals and disease invasion at low densities is not possible.

Assume total host density is fixed in model (S4) and all individuals have equal competitive ability. This means the model dynamics are defined by the *dY/dt, dX/dt* and *dP/dt* equations and the reproduction rates only depend on total host density, e.g., *f*_*I*_(*S, I, C*) = *f*_*I*_(*N*). The equilibrium of the model satisfies

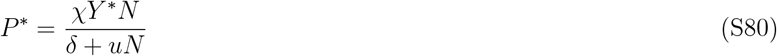

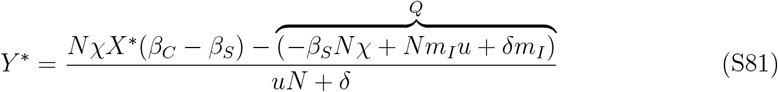

where *Q* = (−*β*_*S*_*Nχ* + *Nm*_*I*_*u* + *δm*_*I*_) = − (ℛ _0_ −1)(*uN* + *δ*)*m*_*I*_. After substituting *P*^∗^ and *Y* ^∗^ into *dX/dt* = 0, we get that the equilibrium values of *X*^∗^ are the roots of the quadratic polynomial

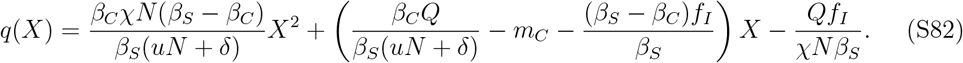

Note that *f*_*I*_(*N*) is a fixed quantity because we assume *N* is fixed. Recall that the roots of the quadratic function *aX*^2^+*bX* +*c* are 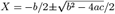 and both roots are positive only if −4*ac* is negative. Applying this to equation (S82) yields the condition (*β*_*S*_ − *β*_*C*_)*Q* < 0, which means *β*_*S*_ − *β*_*C*_ and ℛ _0_ − 1 must have the same sign. Consequently, bistability is possible if either (i) *β*_*S*_ < *β*_*C*_ and _0_ < 1 or (ii) *β*_*S*_ > *β*_*C*_ and ℛ _0_ > 1.

We now rule out the possibility of bistability when *β*_*S*_ > *β*_*C*_ and ℛ _0_ > 1. Equilibrium infection prevalence must satisfy *Y* ^∗^ < 1, which after some algebraic manipulation implies *X*^∗^ < *β*_*S*_(1 −1*/* ℛ _0_)*/*(*β*_*S*_ − *β*_*C*_) = *Q*_2_ where ℛ _0_ = *χβ*_*S*_*N/m*_*I*_(*uN* + *δ*). In order for this condition to be satisfied, the roots of the shifted quadratic polynomial *q*(*X*^∗^ + *Q*_2_) must be negative. Expanding the shifted quadratic polynomial and collecting terms yields *q*(*X*^∗^ + *Q*_2_) = *a*_1_*X*^2^ + *b*_1_*X* + *c* where

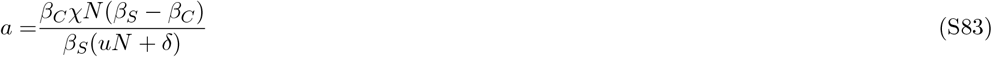

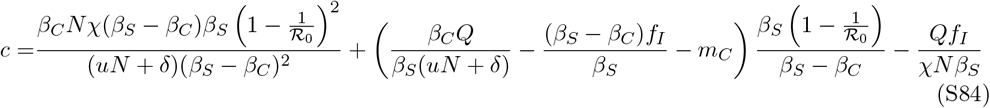

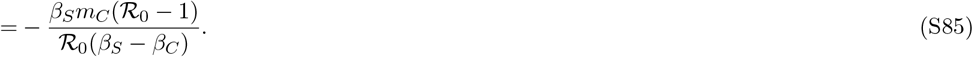

If ℛ _0_ > 1, then it is not possible for *a* and *c* to have the same sign. Because of Descartes’ rule of signs, this means that the shifted polynomial *q*(*X*^∗^ + *Q*_2_) has one positive root. As a result, one of the roots of *q*(*X*^∗^) yields an equilibrium point where *Y* ^∗^ > 1, which is not biologically relevant. Consequently, the model with fixed population size and equal Lotka-Volterra competition coefficients cannot exhibit bistability when ℛ _0_ > 1.

**Figure S5:**
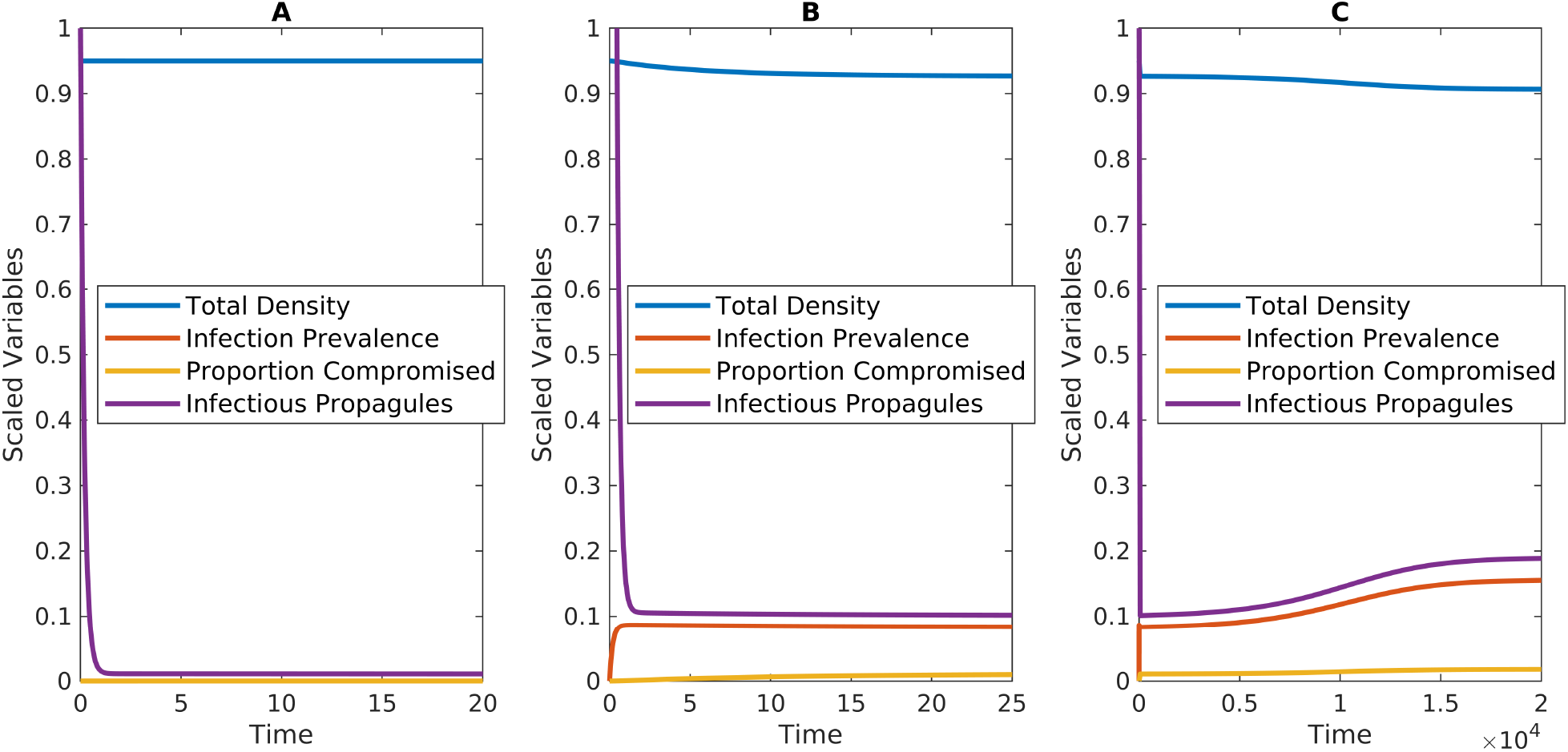
Example of bistability when the basic reproduction number is less than 1 (ℛ _0_ < 1) and compromised individuals have much higher transmission rates than susceptible individuals (*β*_*c*_ > *β*_*S*_). All panels show simulations of a model where there is Lotka-Volterra competition and all individuals have equal competitive ability (see equation (S30)). (A) Convergence to the disease-free equilibrium for sufficiently low initial infectious propagule densities. (B,C) Convergence to the endemic equilibrium for sufficiently high initial infectious propagule densities; both panels show the same simulation over different ranges of time. The parameter values are *r*_*S*_ = 0.2, *r*_*C*_ = 0.2, *r*_*I*_ = 0.2, *K* = 100, *m*_*S*_ = 0.01, *m*_*C*_ = 0.06, *m*_*I*_ = 0.06, *β*_*S*_ = 4 ⋗ 10^−7^, *β*_*C*_ = 45 ⋗ 10^−7^, *χ* = 8100, *u* = 0.05, *δ* = 0.5. For these parameter values, ℛ _0_ = 0.977. The initial conditions (A) (*N* (0), *Y* (0), *X*(0), *P* (0)) = (95, 0, 0, 1000) and (B,C) (*N* (0), *Y* (0), *X*(0), *P* (0)) = (95, 0, 0, 1200000)

#### Section S2.3 SCIDP model with interactions between transgenerational virulence and infection status

We now consider a model where transgenerational virulence affects infected individuals. The SIR-type environmental transmission model describes the dynamics of the densities of susceptible (*S*) and infected (*I*) individuals who are the offspring of susceptible individuals, susceptible (*C*) and infected (*D*) individuals who are the offspring of infected individuals (hereafter, compromised and decimated individuals), and infectious propagules in the environment (*P*). Infection occurs when susceptible or compromised individuals encounter infectious propagules that were shed previously into the environment by infected or decimated individuals.

The equations for the SICD-form of the model are

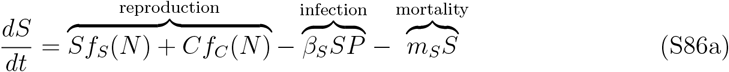

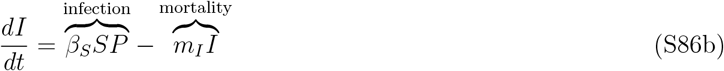

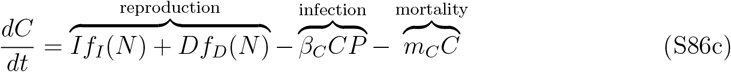

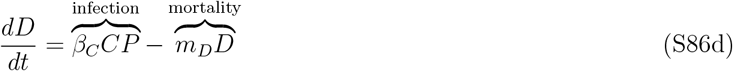

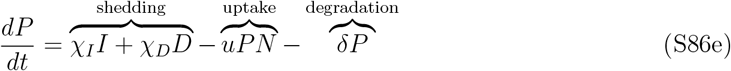

where *N* = *S* + *C* + *I* + *D*. In equation (S86a), susceptible individuals increase due to reproduction by susceptible and compromised individuals (*Sf*_*S*_ and *Cf*_*C*_, respectively) and decrease due to infection (*β*_*S*_*SP*) and non-disease mortality (*m*_*S*_*S*). In equation (S86b), infected individuals increase when susceptible individuals become infected (*β*_*S*_*SP*) and decrease due to mortality from disease and non-disease sources (*m*_*I*_*I*). In equation (S86c), compromised individuals increase due to reproduction by infected individuals (*If*_*I*_) and decimated individuals (*Df*_*D*_) and decrease due to infection (*β*_*C*_*CP*) and mortality due to non-disease sources (*m*_*C*_*C*). In equation (S86d), decimated individuals increase when compromised individuals become infected (*β*_*C*_*CP*) and decrease due to mortality from disease and non-disease sources (*m*_*D*_*D*). In equation (S86e), infectious propagule density increases due to shedding by infected and decimated individuals (*χ*_*I*_*I* + *χ*_*D*_*D*) and decreases due to uptake by all individuals (*uPN*) and degradation (*δP*). Note that the basic reproduction number, ℛ_0_, for model (S86) is identical to the basic reproduction number for model (S1) in Section Section S2.1.1.

We assume the following in model (S86). First, we assume the per capita reproduction rates only depend on total host density and are decreasing functions of total host density, i.e., *f*_*i*_(*N*) = *r*_*i*_*f* (*N*) for *i* ∈ {*S, I, C, D*} where *f*^*′*^(*N*) < 0. This means that individual competitive ability is not affected by infection status or parental infection status.

Second, model (S86) assumes that infected individuals differ based on whether they are the offspring of susceptible or infected individuals. However, we also assume that there are no differences between the infectious propagules shed by infected and decimated individuals. Third, we assume the offspring of compromised individuals are identical to those of susceptible individuals (i.e., infection alters the fitness of the offspring of infected individuals, and not the grandchildren of infected individuals). Fourth, we assume the uptake rates for all individuals are equal (*u*).

Fifth, when compared to susceptible individuals, we assume compromised individuals have (i) equal or higher mortality rates (*m*_*C*_ ≥ *m*_*S*_), (ii) equal or lower reproductive output (*f*_*C*_ ≤ *f*_*S*_; see below), and (iii) higher, equal, or lower transmission rates (*β*_*C*_ > *β*_*S*_, *β*_*C*_ = *β*_*S*_, or *β*_*C*_ < *β*_*S*_, respectively). Sixth, we assume infected individuals have equal or lower reproductive output than susceptible individuals (*f*_*I*_ ≤ *f*_*S*_) and higher mortality rates (*m*_*I*_ > *m*_*S*_). Similarly, decimated individuals have equal or lower reproductive output than compromised individuals (*f*_*D*_ ≤ *f*_*C*_) and higher mortality rates (*m*_*D*_ > *m*_*C*_). When comparing infected and decimated individuals, we assume decimated individuals have lower reproductive output (*f*_*D*_ ≤ *f*_*I*_), higher mortality rates (*m*_*D*_ > *m*_*I*_), and higher, equal, or lower shedding rates (*χ*_*D*_ > *χ*_*I*_, *χ*_*D*_ = *χ*_*I*_, or *χ*_*D*_ < *χ*_*I*_, respectively).

##### Section S2.3.1 Frequency-form of the SCIDP model

To facilitate the mathematical calculations, we transform our variables into total host density (*N* = *S* + *C* + *I* + *D*) and the fractions of susceptible (*W* = *S/N*), compromised (*X* = *C/N*), infected (*Y* = *I/N*), and decimated (*Z* = *D/N*) individuals. Then, we write the model in terms of total host density (*N*), the fractions of compromised (*X* = *C/N*) and decimated (*Z* = *D/N*) individuals, and infection prevalence (Θ = *Y* + *W*, i.e., the total fraction of the population that is infected). Note that infection prevalence includes both infected and decimated individuals and *W* = 1 −*X*− Θ = 1− *X* − *Y* − *Z*.

The frequency form of the SCIDP model is

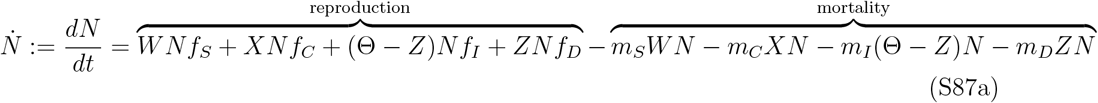

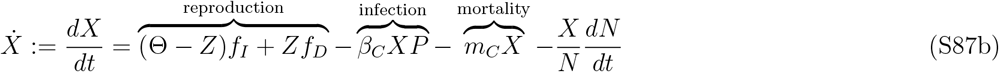

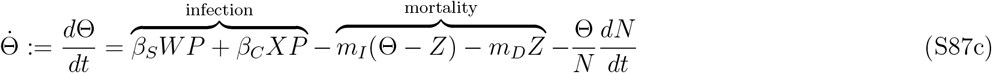

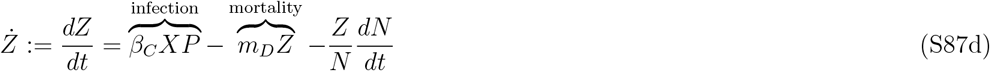

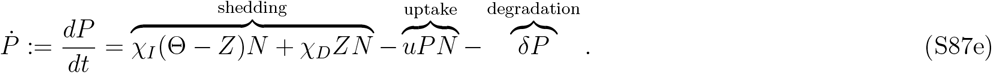

term in each of the middle three equations follows from the quotient rule of calculus, e.g., *dZ/dt* = *d*(*D/N*)*/dt* = (1*/N*)(*dZ/dt*) − (*D/N*)(*dN/dt*).

Our equilibrium-based analyses use the Jacobian of the frequency model (S87). When evaluated at a stable coexistence equilibrium point (*N*^∗^, *X*^∗^, *Y* ^∗^, *Z*^∗^, *P*^∗^), the Jacobian has the structure

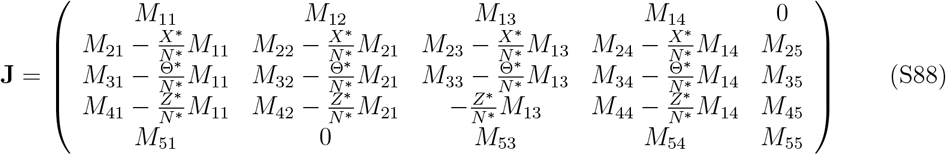

where the terms *M*_*ij*_ are defined by the row reduced form of the Jacobian (**M**), whose entries and sign structure are

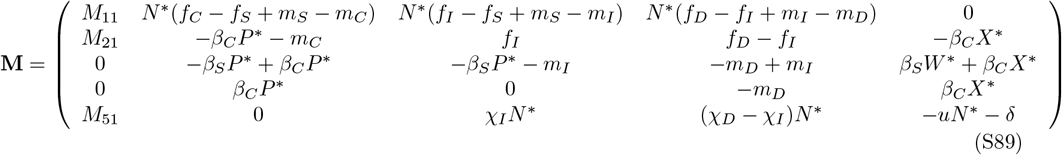

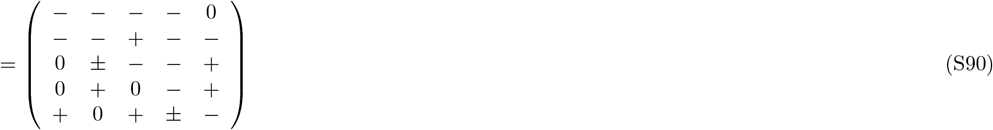

Explanations for the sign of the entries are given below. Note that our analysis focuses on stable coexistence equilibrium points, which means |**J**| < 0.

*M*_11_ Entry *M*_11_ defines the total amount of intraspecific competition experienced by the host species. The sign of this entry is always negative because we assume 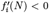 for all *i*. In particular, at a coexistence equilibrium we get

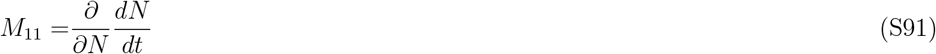

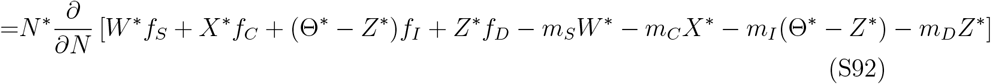

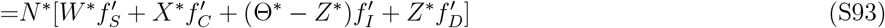

which is negative because each term in the final equation is negative by assumption.

*M*_12_: Entry *M*_12_ is negative because we assume *f*_*C*_ ≤ *f*_*S*_ and *m*_*C*_ ≥ *m*_*S*_.

*M*_13_: Entry *M*_13_ is negative because we assume *f*_*I*_ ≤ *f*_*S*_ and *m*_*I*_ ≥ *m*_*S*_.

*M*_14_: Entry *M*_14_ is negative because we assume *f*_*D*_ ≤ *f*_*I*_ and *m*_*D*_ ≥ *m*_*I*_.

*M*_21_: Entry *M*_21_ is negative because 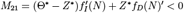.

*M*_51_: Entry *M*_51_ = *χ*_*I*_(Θ^∗^ −*Z*^∗^) + *χ*_*D*_*Z*^∗^ −*uP*^∗^ is positive because the equilibrium condition 0 = *dP/dt* = *χ*_*I*_(Θ^∗^ −*Z*^∗^)*N*^∗^ + *χ*_*D*_*Z*^∗^*N*^∗^ −*uP*^∗^*N*^∗^− *δP*^∗^ implies *χ*_*I*_(Θ^∗^ −*Z*^∗^) + *χ*_*D*_*Z*^∗^ −*uP*^∗^ = *δP*^∗^*/N*^∗^ > 0.

##### Section S2.3.2 Translating predictions to direct transmission models

Our results for the environmental transmission (ET) model (S86) also apply to direct transmission models with density dependent direct transmission (DDDT), frequency dependent direct transmission (FDDT), or direct transmission is that intermediate to DDDT and FDDT. As in Section Section S2.1.4, the reason is that in the limit where the infectious propagule dynamics are much faster than the dynamics of the host classes, the ET model (S86) reduces to model with direct transmission. This means that all of our equilibrium-based predictions for ET pathogens apply to apply DDDT and FDDT pathogens. Mathematical details are provided below; they are nearly identical to those in Section Section S2.1.4.

Assume the infectious propagule dynamics are much faster than the dynamics of the host classes. Mathematically, we assume *χ* is very large and *u* or *δ* are very large such that there is a separation of time scales between the infectious propagule equation and the host class equations. The quasi-steady state density for the infectious propagules is *P*^∗^ = (*χ*_*I*_*I* + *χ*_*D*_*D*)*/*(*δ* + *uN*). Substitution into the equations for the host classes yields

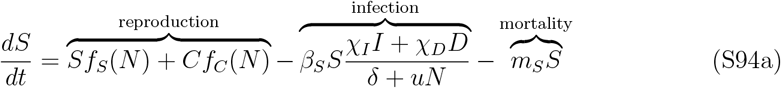

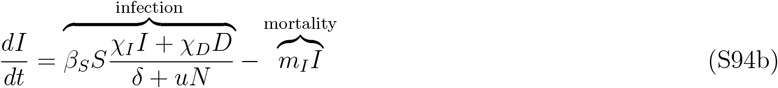

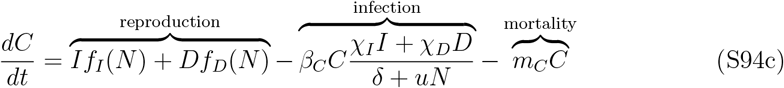

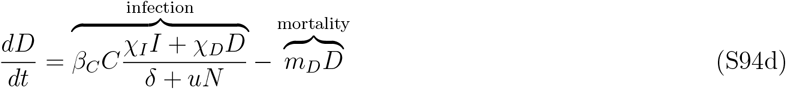

where 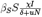 and 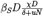 are the direct transmission rates from infected and decimated individuals to susceptible individuals, respectively, and 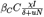 and 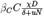 are the direct transmission rates from infected and decimated individuals to compromised individuals, respectively. When *u* = 0 (i.e., uptake of infectious propagules is negligible relative to degradation), the direct transmission rates reduce to the DDDT forms, 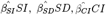, and 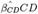, where 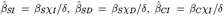, and 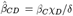. When *δ* = 0 (i.e., degradation of infectious propagules is negligible relative to uptake), the direct transmission rates reduce to the FDDT forms, 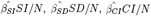, and 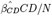, where 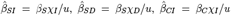, and 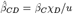. When *δ* and *u* are non-negligible, the transmission rates are intermediate to DDDT and FDDT.

Importantly, the above implies that all of the predictions for ET pathogens can be translated to predictions for DDDT and FDDT pathogens. Specifically, any conditions on *χ*_*I*_, *χ*_*D*_, *u* and *δ* in the ET model can be interpreted in terms conditions on 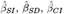, and 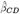 in a direct transmission model.

#### Section S2.4 Analysis of SCIDP model

As before, we use local sensitivity analysis to explore how the equilibrium host density (*N*^∗^), infection prevalence (Θ ^∗^), and infected density (*I*^∗^ + *D*^∗^) depend on the characteristics of the compromised individuals. For a parameter *σ*, the (local) sensitivities of the equilibrium values to *σ* are computed using the derivatives using the derivatives *∂N*^∗^*/∂σ, ∂*Θ^∗^*/∂σ*, and *∂*(*I*^∗^ + *D*^∗^)*/∂σ*. Positive values mean that increases in the parameter cause an increase in the equilibrium value whereas negative values mean that increases in the parameter cause a decrease in the equilibrium value. In the following, we compute the sensitivities to the mortality rates (*m*_*C*_, *m*_*D*_) and reproduction rates (*r*_*C*_, *r*_*D*_) of compromised and decimated individuals, the transmission rate (*β*_*C*_) of compromised individuals, and the shedding rate (*χ*_*D*_) of decimated individuals.

To simplify the computations of the sensitivities, we first set each equation in model (S86) equal to zero and row reduce the equations to get

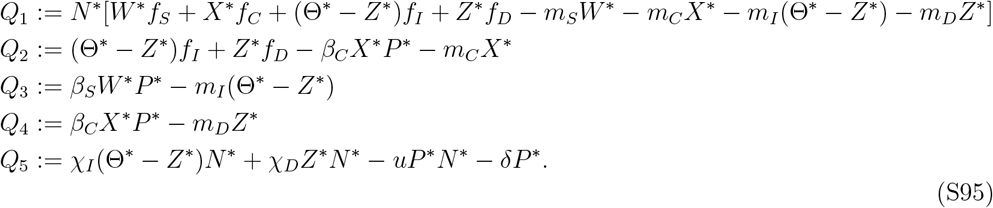

We then compute the sensitivities using the formulas

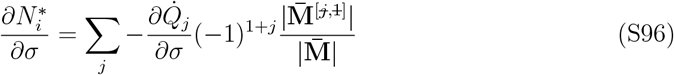

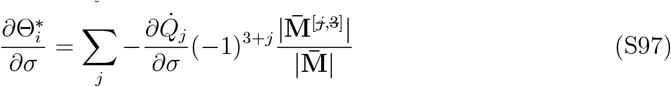

where *σ* is a parameter of interest, 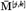 is the submatrix of 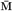 where row *j* and column *k* have been removed, and

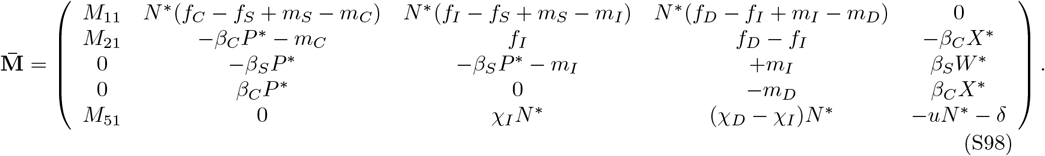

Note that 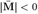 because the properties of the determinant imply that 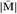 and |**J**| have the same sign.

##### Section S2.4.1 Sensitivities of equilibrium total host density

To get the equations that follow, we expanded the determinants and simplified the formulas by canceling out terms using the equilibrium conditions *Q*_*i*_ = 0.

###### Sensitivity of total host density to the mortality rate of compromised individuals, *m*_*C*_

Algebraic simplification yields

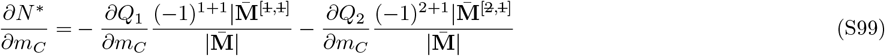

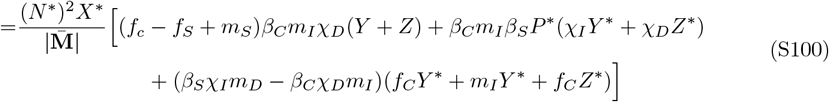

After distributing the factor outside of the brackets, the first term is negative unless *f*_*C*_ < *f*_*S*_, the second term is negative, and the third term is negative unless *β*_*C*_*χ*_*D*_*m*_*I*_ > *β*_*S*_*χ*_*I*_*m*_*D*_. From this, we predict that *∂N*^∗^*/∂m*_*C*_ < 0 unless *f*_*C*_ *≪ f*_*S*_, *χ*_*D*_ *≫ χ*_*I*_, or *β*_*C*_ *≫ β*_*S*_.

###### Sensitivity of total host density to the reproduction rate of compromised individuals, *r*_*C*_

Algebraic simplification yields

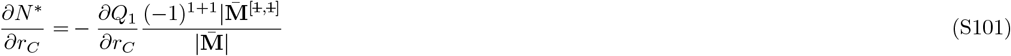

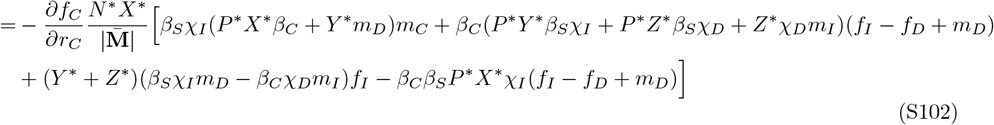

The factor outside of the brackets is positive, the first line is positive, the first term of the second line is positive unless *β*_*S*_*χ*_*I*_*m*_*D*_ − *β*_*C*_*χ*_*D*_*m*_*I*_ < 0 and the last term is negative. Based on this, we predict that *∂N*^∗^*/∂r*_*C*_ is positive unless *β*_*C*_ *≫ β*_*S*_ or *χ*_*D*_ *≫ χ*_*I*_.

###### Sensitivity of total host density to the transmission rate of compromised individuals, *β*_*C*_

Algebraic simplification yields

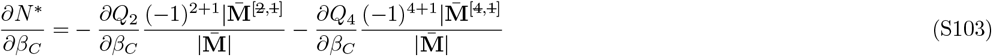

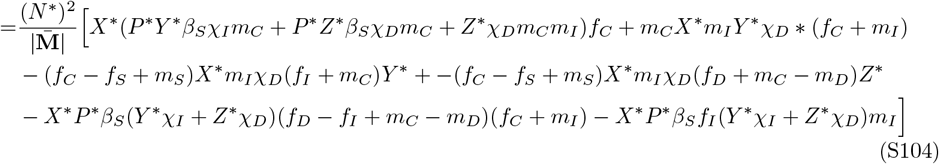

After distributing the factor outside of the brackets, the first line is negative, the second line is negative unless *f*_*D*_ very small and *f*_*C*_ *≈ f*_*S*_, the first term on the third line is negative and the last term is positive. Based on this, we predict *∂N*^∗^*/∂β*_*C*_ is negative unless *f*_*C*_ −*f*_*S*_ + *m*_*S*_ is sufficiently large and negative or *f*_*D*_ + *m*_*C*_ − *m*_*D*_ is sufficiently large and negative.

###### Sensitivity of total host density to the mortality rate of decimated individuals, *m*_*D*_

Algebraic simplification yields

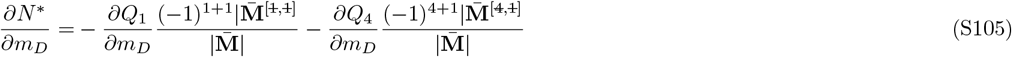

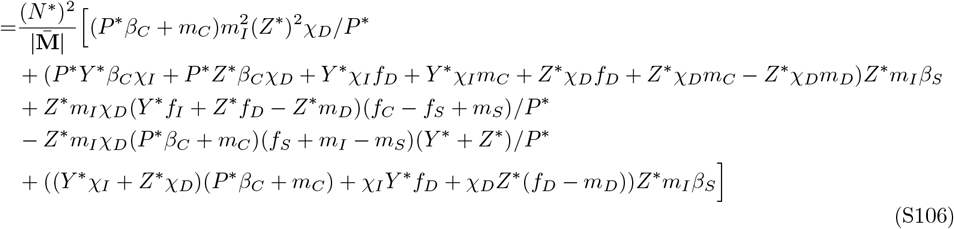

After distributing the negative factor outside of the brackets, the first line is negative, the second line is negative unless *m*_*D*_ *≫ m*_*C*_, the third line is negative unless *f*_*C*_ *≪ f*_*S*_ or *m*_*D*_ is very large, the fourth line is positive, and the fifth line is negative unless *m*_*D*_ is very large. Thus, we predict that *∂N*^∗^*/∂m*_*D*_ is negative unless *f*_*C*_ *≪ f*_*S*_ or *m*_*D*_ *≫ m*_*I*_.

###### Sensitivity of total host density to the reproduction rate of decimated individuals, *r*_*D*_

Algebraic simplification yields

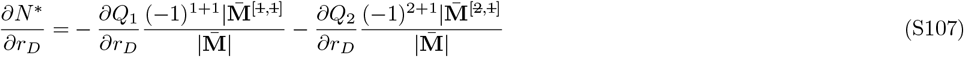

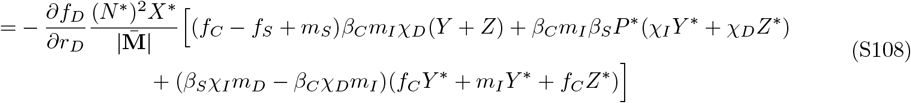

The factor outside of the brackets is positive, the first term is positive unless *f*_*C*_ < *f*_*S*_, the second term is positive, and the third term is positive unless *β*_*C*_*χ*_*D*_*m*_*I*_ > *β*_*S*_*χ*_*I*_*m*_*D*_. From this, we predict that *∂N*^∗^*/∂r*_*D*_ > 0 unless *f*_*C*_ *≪ f*_*S*_, *χ*_*D*_ *≫ χ*_*I*_, or *β*_*C*_ *≫ β*_*S*_.

###### Sensitivity of total host density to the shedding rate of decimated individuals, *χ*_*D*_

Algebraic simplification yields

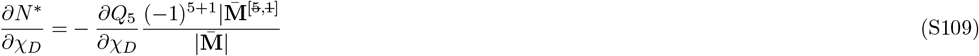

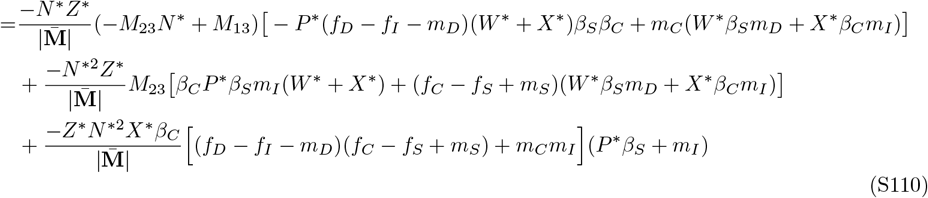

The first line is negative because *M*_23_*N*^∗^ + *M*_13_ < 0. The second line is negative unless *f*_*C*_ −*f*_*S*_ + *m*_*S*_ is negative. The third line is negative unless *f*_*C*_ −*f*_*S*_ + *m*_*S*_ is negative or *m*_*C*_*m*_*I*_ is large in magnitude. Consequently, we predict that *∂N*^∗^*/∂χ*_*D*_ < 0 unless *f*_*C*_ − *f*_*S*_ + *m*_*S*_ is sufficiently large and negative or *m*_*C*_ and *m*_*I*_ are both very large.

##### Section S2.4.2 Sensitivities of equilibrium infection prevalence

To get the equations that follow, we expanded the determinants and organized the terms based on their dependencies on *M*_11_ − *N*^∗^*M*_13_ < 0 and *M*_13_ < 0. In addition, we simplified some formulas by canceling out terms using the equilibrium conditions *Q*_*i*_ = 0.

###### Sensitivity of infection prevalence to the mortality rate of compromised individuals, *m*_*C*_

Algebraic simplification yields

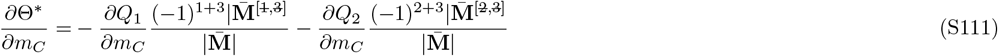

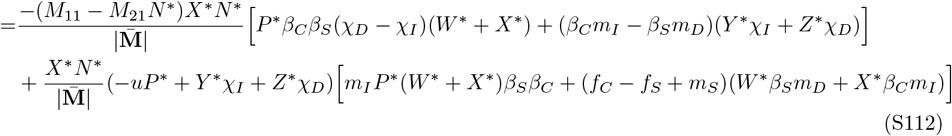

In the first line, the factor outside of the brackets is negative because *M*_11_ − *M*_21_*N*^∗^ < 0. After distributing, the first term is negative unless *χ*_*D*_ − *χ*_*I*_ < 0 and the second term is negative unless *β*_*C*_*m*_*I*_ − *β*_*S*_*m*_*D*_ < 0. In the second line, the factor outside of the brackets is negative because −*uP*^∗^ + *Y* ^∗^*χ*_*I*_ + *Z*^∗^*χ*_*D*_ = *δP*^∗^*/N*^∗^ > 0. After distributing, the first term is negative and the second term is negative unless *f*_*C*_ − *f*_*S*_ + *m*_*S*_ < 0. Based on the signs of the terms, we predict *∂*Θ^∗^*/∂m*_*C*_ < 0 unless *f*_*C*_ − *f*_*S*_ + *m*_*S*_, *χ*_*D*_ −*χ*_*I*_, or *β*_*C*_*m*_*I*_ −*β*_*S*_*m*_*D*_ is negative and large in magnitude.

###### Sensitivity of infection prevalence to the reproduction rate of compromised individuals, *r*_*C*_

Algebraic simplification yields

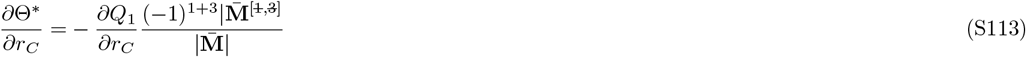

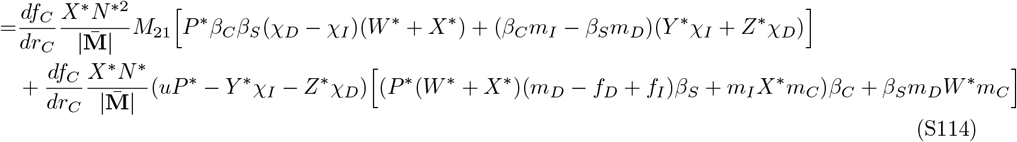

For the first line, after distributing the positive factor outside of the brackets, the first term is positive unless *χ*_*D*_ − *χ*_*I*_ < 0 and the second term is positive unless *β*_*C*_*m*_*I*_ − *β*_*S*_*m*_*D*_ < 0. The second line is positive. Based on the signs of the terms, we expect *∂*Θ*/∂r*_*C*_ > 0 unless *χ*_*D*_ − *χ*_*I*_ or *β*_*C*_*m*_*I*_ − *β*_*S*_*m*_*D*_ is negative and large in magnitude.

###### Sensitivity of infection prevalence to the transmission rate of compromised individuals, *β*_*C*_

Algebraic simplification yields

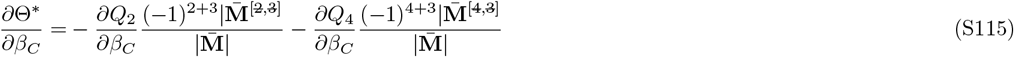

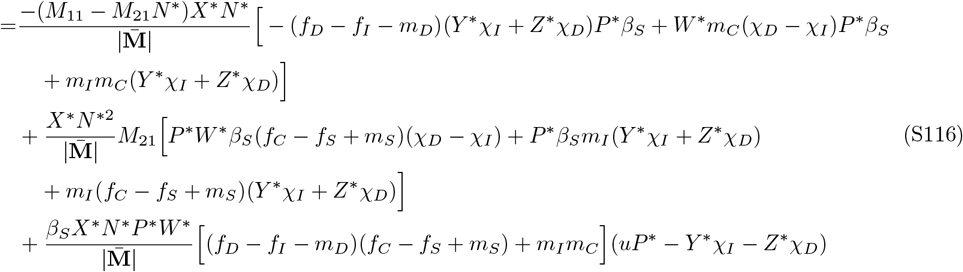

For the first line, after distributing the negative factor outside of the brackets, the first and third terms are negative and the second term is negative unless *χ*_*D*_ − *χ*_*I*_ < 0. For the second line, after distributing the positive factor outside of the brackets, the first term is positive unless *f*_*C*_ − *f*_*S*_ + *m*_*S*_ < 0 or *χ*_*D*_ − *χ*_*I*_ < 0, the second term is positive, and the third term is positive unless *f*_*C*_ − *f*_*S*_ + *m*_*S*_ < 0. For the third line, after distributing the positive factors outside of the brackets, the terms inside the brackets are positive unless *f*_*C*_ − *f*_*S*_ + *m*_*S*_ < 0. Based on the signs of the terms, we expect this equation to be positive unless *f*_*C*_ − *f*_*S*_ + *m*_*S*_ is sufficiently large and negative or *χ*_*D*_ − *χ*_*I*_ is sufficiently large and negative.

###### Sensitivity of infection prevalence to the mortality rate of decimated individuals, *m*_*D*_

Algebraic simplification yields

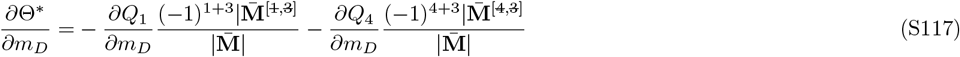

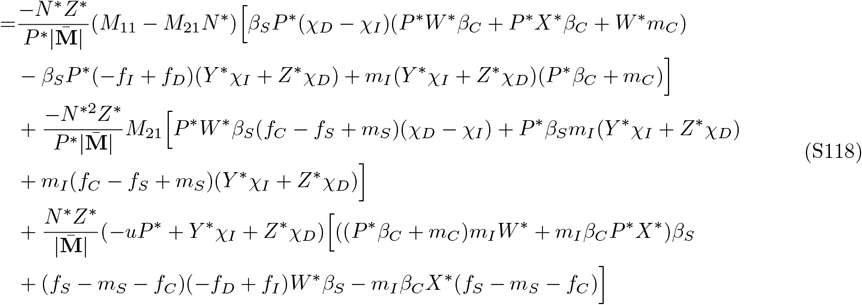

For the first line, after distributing the negative factor outside of the brackets, the first term is negative unless *χ*_*D*_ − *χ*_*I*_ < 0 and the second and third terms are negative. For the second line, after distributing the negative factor outside of the brackets, the first term is negative unless *f*_*C*_ − *f*_*S*_ + *m*_*S*_ < 0 or *χ*_*D*_ − *χ*_*I*_ < 0, the second term is negative, and the third term is negative unless *f*_*C*_ − *f*_*S*_ + *m*_*S*_ < 0. For the third line, after distributing the negative factor outside of the brackets, the first term is negative, the second term is negative unless *f*_*C*_ − *f*_*S*_ + *m*_*S*_ < 0, and the third term is positive unless *f*_*C*_ − *f*_*S*_ + *m*_*S*_ < 0. Based on the signs of the terms, we expect this equation to be negative unless *f*_*C*_− *f*_*S*_ + *m*_*S*_ or *χ*_*D*_ − *χ*_*I*_ is negative and large in magnitude.

###### Sensitivity of infection prevalence to the reproduction rate of decimated individuals, *r*_*D*_

Algebraic simplification yields

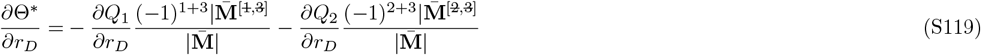

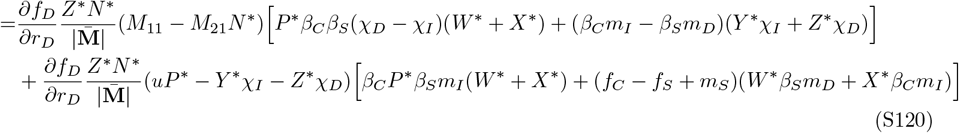

In the first line, the factor outside of the brackets is positive. After distributing, the first term is positive unless *χ*_*D*_ − *χ*_*I*_ < 0 and the second term is positive unless *β*_*C*_*m*_*I*_ − *β*_*S*_*m*_*D*_ < 0. In the second line, the factor outside of the brackets is positive. After distributing, the first term is positive and the second term is positive unless *f*_*C*_ − *f*_*S*_ + *m*_*S*_ < 0. Based on the signs of the terms, we predict *∂*Θ^∗^*/∂r*_*D*_ > 0 unless *f*_*C*_ − *f*_*S*_ + *m*_*S*_, *χ*_*D*_ −*χ*_*I*_, or *β*_*C*_*m*_*I*_ − *β*_*S*_*m*_*D*_ is negative and large in magnitude.

###### Sensitivity of infection prevalence to the shedding rate of decimated individuals, *χ*_*D*_

Algebraic simplification yields

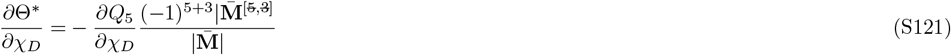

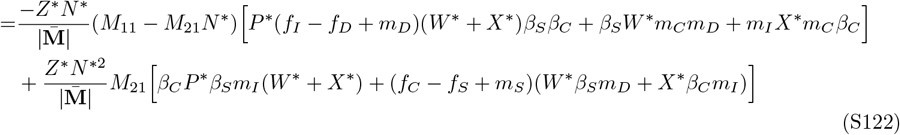

The first line is positive. The second line is positive unless *f*_*C*_ − *f*_*S*_ + *m*_*S*_ < 0. Based on the signs of the terms, we predict this equation to be positive unless *f*_*C*_ − *f*_*S*_ + *m*_*S*_ is negative and large in magnitude.

##### Section S2.4.3 Sensitivities of equilibrium total infected density

We compute the sensitivities of total infected density using the sensitivities of infection prevalence and total host density via the quotient rule from calculus. As noted earlier, the advantage of this approach is that it allows us to determine the conditions under which the sensitivities of total infected density and infection prevalence have the same or opposite signs. In particular, the sensitivity to parameter *σ* becomes

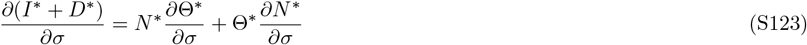

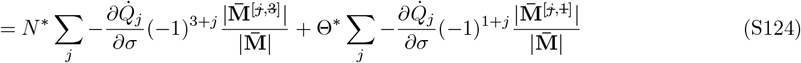

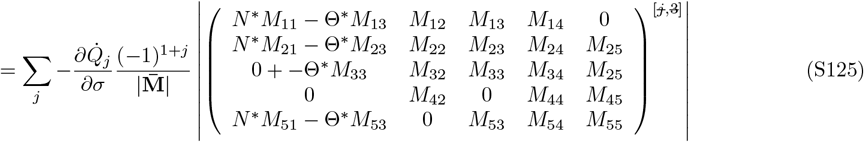

where the superscript for the matrix indicates that the *j*th row and 3rd column are removed. For a given parameter, the equations for the sensitivity of infection prevalence (*∂θ*^∗^*/∂σ*) and the sensitivity of total infected density (*∂*(*I*^∗^ + *D*^∗^)*/∂σ*) differ in sign only if for some entry in the first column of the matrix, the two terms have opposite signs and the second is larger in magnitude.

The first entry of column 1 can cause the sensitivities of infection prevalence and total infected density to have different signs if

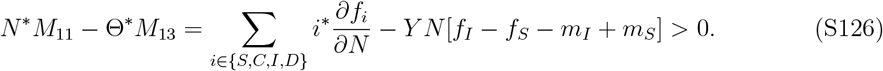

This requires that infected individuals have much smaller reproduction rates and much larger mortality rates than susceptible individuals (*f*_*I*_ ≪ *f*_*S*_ and *m*_*I*_ ≫ *m*_*S*_). The second entry of column 1 is unlikely cause the sensitivities of infection prevalence and total infected density to have different signs. This is because *N*^∗^*M*_21_ − Θ^∗^*M*_23_ is always negative. The third entry of column 1 can cause the sensitivities of infection prevalence and total infected density to have different signs if *M*_33_ is sufficiently large in magnitude. This requires that susceptible individuals have large transmission rates (*β*_*S*_ large) and compromised individuals have large mortality rates (*m*_*C*_ large). The fourth entry of column 1 can cause the sensitivities of infection prevalence and total infected density to have different signs if *N*^∗^*M*_51_ − Θ ^∗^*M*_53_ = *χ*_*D*_*Z*^∗^*N*^∗^ − *uP*^∗^*N*^∗^ < 0 is sufficiently large in magnitude. This requires that infected individuals have sufficiently large shedding rates relative to decimated individuals (*χ*_*I*_ sufficiently large).

In total, the signs of the sensitivities for infection prevalence (*∂*Θ^∗^*/∂σ*) and total infected density (*∂*(*I*^∗^ + *D*^∗^)*/∂σ*) can differ when

i. infected individuals have much smaller reproduction rates and much larger mortality rates than susceptible individuals (*f*_*I*_ *≪ f*_*S*_ and *m*_*I*_ *≫ m*_*S*_),
ii. susceptible individuals have large transmission rates (*β*_*S*_ large) and compromised individuals have large mortality rates (*m*_*C*_ large), or
iii. infected individuals have much larger shedding rates than decimated individuals (*χ*_*I*_ *≫ χ*_*D*_).

Under any of these conditions changes in the parameters for compromised or decimated individuals cause large changes in total host density. If the change in total host density is in the opposite direction of the changes in infection prevalence, then infection prevalence and total infected density to change opposite directions. Note that because different rows of the matrix are removed when computing the sensitivities for the different parameters, not all conditions apply to every parameter.

###### Sensitivity of total infected density to the mortality rate of compromised individuals, *m*_*C*_

The sensitivity of total infected density is

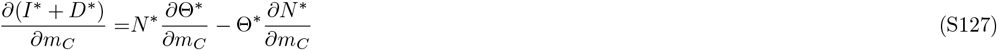

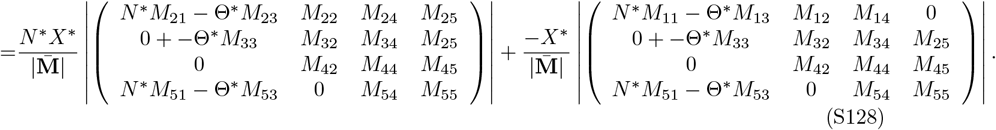

The signs of the sensitivities for infection prevalence and total infected density can differ when conditions (i), (ii), or (iii) are satisfied.

Sensitivity of total infected density to the reproduction rate of compromised individuals to the reproduction rate of compromised individuals, *r*_*C*_: The sensitivity of total infected density is

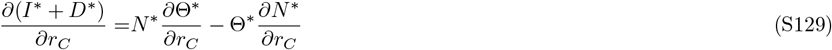

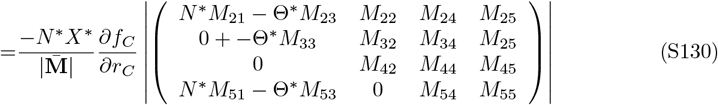

The signs of the sensitivities for infection prevalence and total infected density can differ when conditions (ii) or (iii) are satisfied.

Sensitivity of total infected density to the transmission rate of compromised individuals, *β*_**C**_: The sensitivity of total infected density is

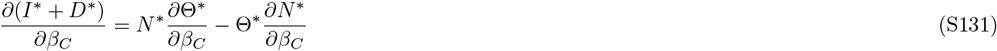

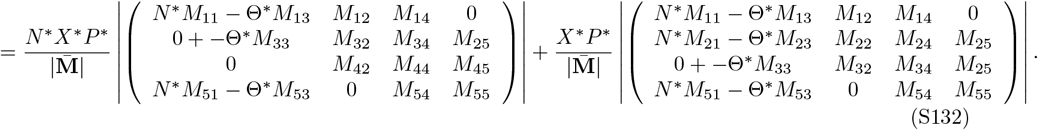

The signs of the sensitivities for infection prevalence and total infected density can differ when conditions (i), (ii), or (iii) are satisfied.

###### Sensitivity of total infected density to the mortality rate of decimated individuals, *m*_*D*_

The sensitivity of total infected density is

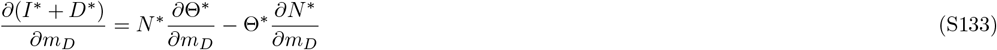

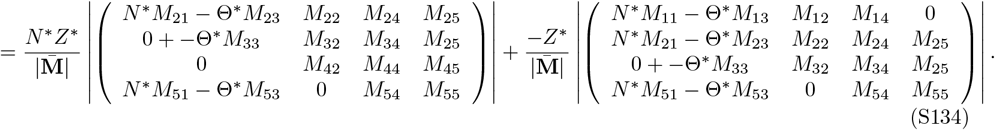

The signs of the sensitivities for infection prevalence and total infected density can differ when conditions (i), (ii), or (iii) are satisfied.

###### Sensitivity of total infected density to the reproduction rate of decimated individuals, *r*_*D*_

The sensitivity of total infected density is

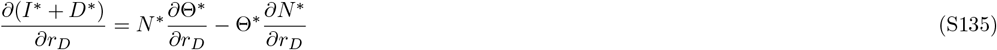

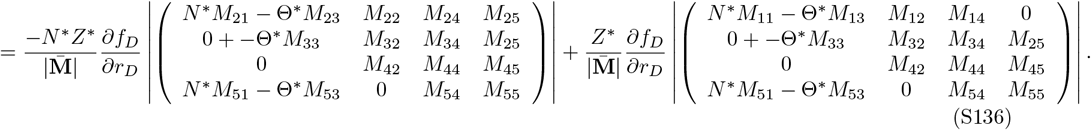

The signs of the sensitivities for infection prevalence and total infected density can differ when conditions (i), (ii), or (iii) are satisfied.

###### Sensitivity of total infected density to the shedding rate of decimated individuals, *χ*_*D*_

The sensitivity of total infected density is

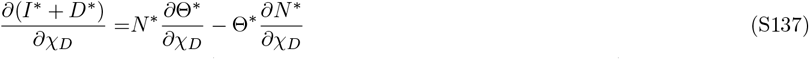

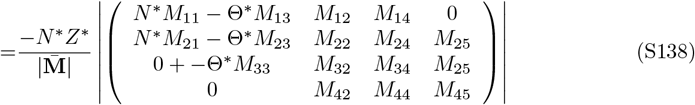

The signs of the sensitivities for infection prevalence and total infected density can differ when conditions (i) or (ii) are satisfied.

##### Section S2.4.4 Summary and interpretation of sensitivity results

The expected signs of the sensitivities and the conditions under which the sensitivities have the opposite signs are shown in Table S3. In Figures S6-S8, examples where the sensitivities have the expected signs are plotted in bluer hues and examples with the opposite signs are plotted in redder hues.

###### Total host density

We predict that in many systems, total host density decreases with increased mortality, reduced reproduction, and increased transmission rates of compromised individuals and increased mortality, reduced reproduction, and increased shedding of decimated individuals (bluer curves in Figure S6). However, counter-intuitively, those predictions can be reversed if compromised individuals have much lower reproduction rates (*f*_*C*_ *≪ f*_*S*_; red curves in Figure S6C) or much larger transmission rates (*β*_*C*_ ≫ *β*_*S*_) than susceptible individuals or decimated individuals have much higher shedding rates than infected individuals (*χ*_*D*_ ≫ *χ*_*I*_; red curves in Figure S6ADE); see Table S3 for a complete list of conditions. We note that we were unable to find numerical examples where total host density increased with lower compromised reproduction rates or higher decimated shedding rates (hence why all curves are increasing in Figure S6B and decreasing in Figure S6F, respectively).

The counter-intuitive responses in total host density allow for the possibility that total host density is higher with transgenerational virulence than not. However, we were unable to find any numerical examples where total host density was higher with transgenerational virulence. We predict that higher total density with transgenerational virulence will be exceeding rare, if not impossible, in nature because the above conditions for the counter-intuitive responses cause large decreases in total host density (i.e., low compromised reproduction rates, high compromised transmission rates, and high decimated shedding rates cause large decreases in total host density).

###### Infection Prevalence

We predict that in many systems, infection prevalence decreases with increased mortality and reduced reproduction of compromised and decimated individuals, decreased transmission rates of compromised individuals, and decreased shedding of decimated individuals (bluer curves in Figure S7). However, counter-intuitively, those predictions can be reversed if compromised individuals have much lower reproduction rates (*f*_*C*_ *≪ f*_*S*_; red curves in Figure S7C) or much lower transmission rates (*β*_*C*_ ≪ *β*_*S*_) than susceptible individuals or decimated individuals have much lower shedding rates than infected individuals (*χ*_*D*_ ≪ *χ*_*I*_; red curves in Figure S7ABE); see Table S3 for a complete list of conditions. We note that we were unable to find numerical examples where infection prevalence decreased with lower decimated reproduction rates or higher decimated shedding rates (hence why all curves are decreasing in Figure S7D and increasing in Figure S7F, respectively).

The counter-intuitive responses in infection prevalence allow for the possibility that infection prevalence is greater with transgenerational virulence than not. We were only able to find such examples when compromised individuals had much higher transmission rates than susceptible individuals (*β*_*C*_ *≫ β*_*S*_) or decimated individuals had much higher shedding rates than infected individuals (*χ*_*D*_ ≫ *χ*_*I*_). We predict that greater infection prevalence with transgenerational virulence will be exceeding rare, if not impossible, whenever those two conditions are not met (i.e, when compromised individuals have lower transmission rates than susceptible individuals and decimated individuals have lower shedding rates than infected individuals). The reason is that the conditions for the counter-intuitive responses in infection prevalence cause large decreases in infection prevalence (e.g., low compromised reproduction rates, low compromised transmission rates, and low decimated shedding rates cause large decreases in infection prevalence).

Total infected density: In many systems, total infected density changes in the same direction as infection prevalence; see previous paragraph and Table S3 for specific conditions. Figure S8 shows examples where the responses in infected density match intuition (bluer curves) and do not (red curve). The responses in total infected density can be the opposite of the responses in infection prevalence when the effects of transgenerational virulence cause large changes in total host density. This is more likely to occur when (i) infected individuals have much lower reproduction rates and much higher mortality rates than susceptible individuals (*f*_*I*_ *≪ f*_*S*_, *m*_*I*_ *≫ m*_*S*_), (ii) infected individuals have high mortality rates and susceptible individuals have high transmission rates (*m*_*I*_ and *β*_*I*_ large), or (iii) infected individuals have sufficiently large shedding rates (*χ*_*I*_ large; right side of S8F). We note that we were unable to find numerical examples where total infected density decreased with higher compromised reproduction rates (hence why all curves are increasing in Figure S8D). If this holds generally, then increased reproduction rates of compromised individuals would cause infection prevalence and total infected density to responses in opposite directions whenever infection prevalence decreases with increased reproduction rates of compromised individuals.

###### Direct transmission models

All of the above predictions directly translate to predictions for direct transmission pathogens with density-dependent direct transmission, frequencydependent direct transmission, or transmission that is intermediate; see Section Section S2.3.2 for mathematical details. The only differences are that (i) conditions about large and small shedding rates (*χ*_*j*_ for *j ∈* {*I, D*}) should be interpreted in terms of large and small direct transmission parameters (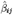for *i ∈* {*S, C*} and *j ∈* {*I, D*}) and (ii) conditions about large and small infection parameters (*β*_*i*_ for *i*∈ {*S, C*}) in the environmental transmission model should be interpreted in terms of large and small direct transmission parameters (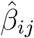for *i ∈* {*S, C*} and *j ∈* {*I, D*}) in the direct transmission models.

**Table S3:** Predicted sensitivities for parameters of compromised and decimated individuals.

| Sensitivity | Expected sign | Conditions for opposite sign |
| --- | --- | --- |
| <u>Total density</u> |  |  |
| $\frac{\partial N^*}{\partial m_C}$ | – | $f_C \ll f_S, \chi_D \gg \chi_I, \beta_C \gg \beta_S$ |
| $\frac{\partial N^*}{\partial r_C}$ | + | $\chi_D \gg \chi_I, \beta_C \gg \beta_S$ |
| $\frac{\partial N^*}{\partial \beta_C}$ | – | $f_C \ll f_S$ |
| $\frac{\partial N^*}{\partial m_D}$ | + | $f_C \ll f_S, m_D \gg m_I$ |
| $\frac{\partial N^*}{\partial r_D}$ | + | $f_C \ll f_S, \chi_D \gg \chi_I, \beta_C \gg \beta_S$ |
| $\frac{\partial N^*}{\partial \chi_D}$ | – | $f_C \ll f_S$ |
| <u>Infection Prevalence</u> |  |  |
| $\frac{\partial \Theta^*}{\partial m_C}$ | – | $f_C \ll f_S, \chi_D \ll \chi_I, \beta_C \ll \beta_S$ |
| $\frac{\partial \Theta^*}{\partial r_C}$ | + | $\chi_D \ll \chi_I, \beta_C \ll \beta_S$ |
| $\frac{\partial \Theta^*}{\partial \beta_C}$ | + | $f_C \ll f_S, \chi_D \ll \chi_I$ |
| $\frac{\partial \Theta^*}{\partial m_D}$ | – | $f_C \ll f_S, \chi_D \ll \chi_I$ |
| $\frac{\partial \Theta^*}{\partial r_D}$ | + | $f_C \ll f_S, \chi_D \ll \chi_I, \beta_C \ll \beta_S$ |
| $\frac{\partial \Theta^*}{\partial \chi_D}$ | + | $f_C \ll f_S$ |
| <u>Infected Density<sup>†</sup></u> |  |  |
| $\frac{\partial(I^* + D^*)}{\partial m_C}$ | – | $f_C \ll f_S, \chi_D \ll \chi_I, \beta_C \ll \beta_S$, conditions (i)-(iii) |
| $\frac{\partial(I^* + D^*)}{\partial r_C}$ | + | $\chi_D \ll \chi_I, \beta_C \ll \beta_S$, conditions (ii)-(iii) |
| $\frac{\partial(I^* + D^*)}{\partial \beta_C}$ | + | $f_C \ll f_S, \chi_D \ll \chi_I$, conditions (i)-(iii) |
| $\frac{\partial(I^* + D^*)}{\partial m_D}$ | – | $f_C \ll f_S, \chi_D \ll \chi_I$, conditions (i)-(iii) |
| $\frac{\partial(I^* + D^*)}{\partial r_D}$ | + | $f_C \ll f_S, m_D \gg m_I$, conditions (i)-(iii) |
| $\frac{\partial(I^* + D^*)}{\partial \chi_D}$ | + | $f_C \ll f_S$, conditions (i)-(ii) |

<sup>†</sup>The sensitivities for infection prevalence and infected density can differ when (i) infected individuals have much smaller reproduction rates and much larger mortality rates than susceptible individuals ( $f_I \ll f_S$ and $m_I \gg m_S$ ), (ii) susceptible individuals have large transmission rates ( $\beta_S$ large) and compromised individuals have large mortality rates ( $m_C$ large), or (iii) compromised individuals sufficiently large shedding rates ( $\chi_C$ large).

**Figure S6:**
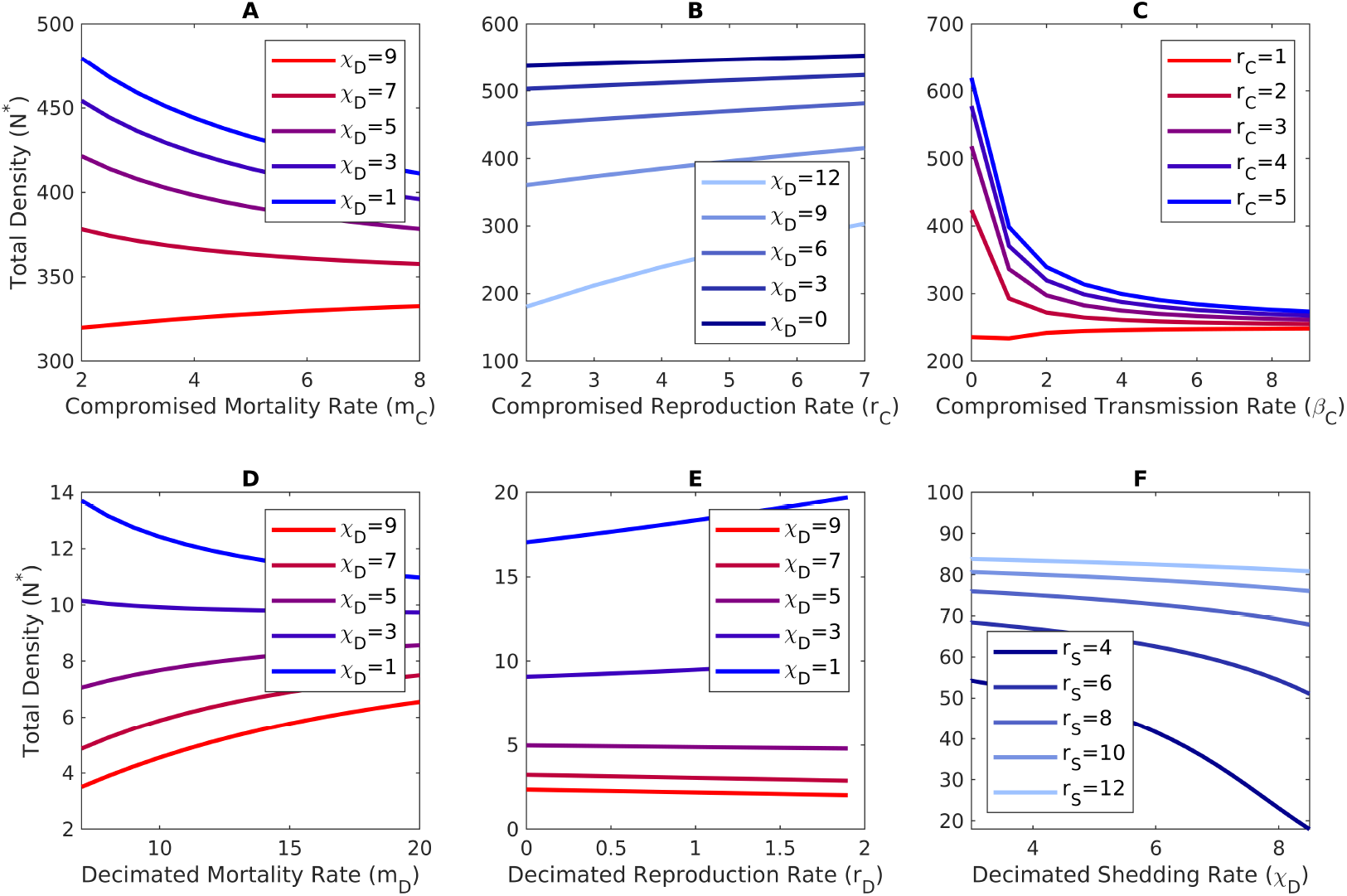
Responses of total host density to the effects of transgenerational virulence. In each panel, bluer curves illustrate the responses expected in most systems and redder curves illustrate responses of the opposite sign. (A) Total density decreases with increased mortality of compromised individuals (bluer curves), unless decimated individuals have sufficiently high shedding rates (redder curves). (B) Total density increases with increased reproduction of compromised individuals (all curves). In principle the opposite response can occur (see Table S3 for conditions), but we did not find any numerical examples. (C) Total density decreases with increase transmission rate of compromised individuals (bluer curves), unless susceptible individuals have sufficiently higher reproduction rates than compromised individuals. (D) Total density decreases with increased mortality of decimated individuals (bluer curves), unless decimated individuals have sufficiently high shedding rates (redder curves). (E) Total density increases with increased reproduction of decimated individuals (bluer curves), unless decimated individuals have sufficiently high shedding rates (redder curves). (F) Total density decreases with increased shedding rates of decimated individuals (all curves). In principle the opposite response can occur (see Table S3 for conditions), but we did not find any numerical examples. The parameter values are (A) *r*_*S*_ = 6, *r*_*C*_ = 6, *r*_*I*_ = 3, *r*_*D*_ = 3, *K* = 1000, *m*_*S*_ = 2, *m*_*I*_ = 10, *m*_*D*_ = 10, *β*_*S*_ = 2, *β*_*C*_ = 2, *χ*_*I*_ = 6, *u* = 1, *δ* = 2; (B) *r*_*S*_ = 7, *r*_*I*_ = 4, *r*_*D*_ = 4, *K* = 1000, *m*_*S*_ = 2, *m*_*C*_ = 3, *m*_*I*_ = 10, *m*_*D*_ = 10, *β*_*S*_ = 2, *β*_*C*_ = 2, *χ*_*I*_ = 6, *u* = 1, *δ* = 2; and (C) *r*_*S*_ = 6, *r*_*I*_ = 3, *r*_*D*_ = 3, *K* = 1000, *m*_*S*_ = 3, *m*_*C*_ = 1, *m*_*I*_ = 8, *m*_*D*_ = 8, *β*_*S*_ = 3, *χ*_*I*_ = 10, *χ*_*D*_ = 10, *u* = 1, *δ* = 1; (D) *r*_*S*_ = 4, *r*_*C*_ = 4, *r*_*I*_ = 2, *r*_*D*_ = 2, *K* = 100, *m*_*S*_ = 1, *m*_*C*_ = 4, *m*_*I*_ = 2, *β*_*S*_ = 4, *β*_*C*_ = 4, *χ*_*I*_ = 3, *u* = 1.5, *δ* = 1.5; (E) *r*_*S*_ = 4, *r*_*C*_ = 4, *r*_*I*_ = 2, *K* = 100, *m*_*S*_ = 1, *m*_*C*_ = 1, *m*_*I*_ = 2, *m*_*D*_ = 3, *β*_*S*_ = 4, *β*_*C*_ = 2, *χ*_*I*_ = 10, *u* = 2, *δ* = 4; and (F) *r*_*C*_ = 2, *r*_*I*_ = 2, *r*_*D*_ = 2, *K* = 100, *m*_*S*_ = 1, *m*_*C*_ = 2, *m*_*I*_ = 3, *m*_*D*_ = 3, *β*_*S*_ = 4, *β*_*C*_ = 4, *χ*_*I*_ = 3, *u* = 3, *δ* = 3.

**Figure S7:**
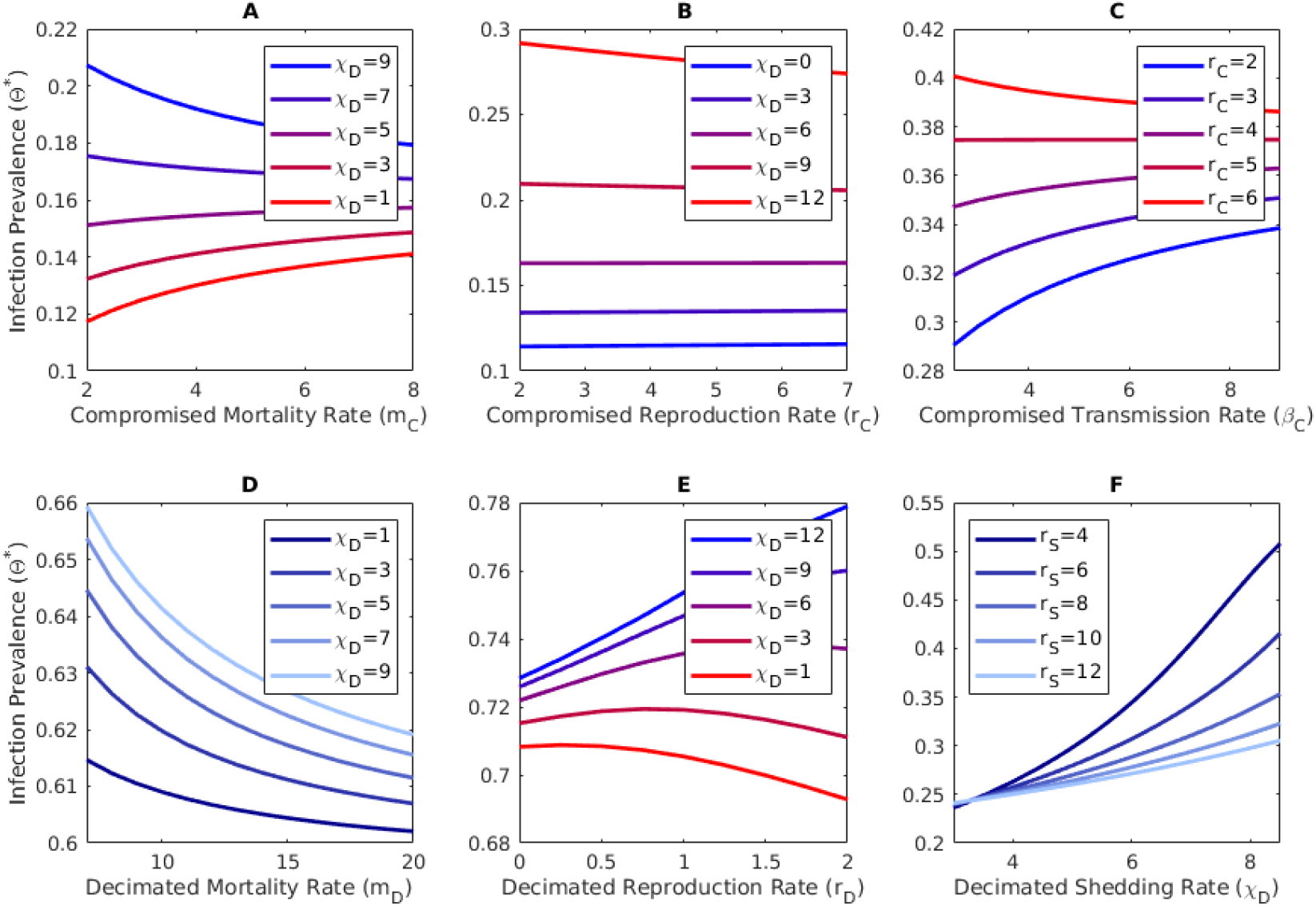
Responses of infection prevalence to the effects of transgenerational virulence. In each panel, bluer curves illustrate the responses expected in most systems and redder curves illustrate responses of the opposite sign. (A) Infection prevalence decreases with increased mortality of compromised individuals (bluer curves), unless decimated individuals have sufficiently high shedding rates (redder curves). (B) Infection prevalence increases with increased reproduction of compromised individuals (bluer curves), unless decimated individuals have sufficiently high shedding rates (redder curves). (C) Infection prevalence increases with increase transmission rate of compromised individuals (bluer curves), unless compromised individuals have sufficiently high reproduction rates. (D) Infection prevalence decreases with increased mortality of decimated individuals (all curves). In principle the opposite response can occur (see Table S3 for conditions), but we did not find any numerical examples. (E) Infection prevalence increases with increased reproduction of decimated individuals (bluer curves), unless decimated individuals have sufficiently high shedding rates (redder curves). (F) Infection prevalence increases with increased shedding rates of decimated individuals (all curves). In principle the opposite response can occur (see Table S3 for conditions), but we did not find any numerical examples. Parameter values are (A) *r*_*S*_ = 6, *r*_*C*_ = 6, *r*_*I*_ = 3, *r*_*D*_ = 3, *K* = 1000, *m*_*S*_ = 2, *m*_*I*_ = 10, *m*_*D*_ = 10, *β*_*S*_ = 2, *β*_*C*_ = 2, *χ*_*I*_ = 6, *u* = 1, *δ* = 2; (B) *r*_*S*_ = 7, *r*_*I*_ = 4, *r*_*D*_ = 4, *K* = 1000, *m*_*S*_ = 2, *m*_*C*_ = 3, *m*_*I*_ = 10, *m*_*D*_ = 10, *β*_*S*_ = 2, *β*_*C*_ = 2, *χ*_*I*_ = 6, *u* = 1, *δ* = 2; (C) *r*_*S*_ = 6, *r*_*I*_ = 4, *r*_*D*_ = 4, *K* = 1000, *m*_*S*_ = 1.5, *m*_*C*_ = 1, *m*_*I*_ = 3, *m*_*D*_ = 3, *β*_*S*_ = 6, *χ*_*I*_ = 10, *χ*_*D*_ = 10, *u* = 1, *δ* = 1; (D) *r*_*S*_ = 4, *r*_*C*_ = 4, *r*_*I*_ = 2, *r*_*D*_ = 2, *K* = 100, *m*_*S*_ = 1, *m*_*C*_ = 4, *m*_*I*_ = 2, *β*_*S*_ = 4, *β*_*C*_ = 4, *χ*_*I*_ = 3, *u* = 1.5, *δ* = 1.5; (E) *r*_*S*_ = 4, *r*_*C*_ = 4, *r*_*I*_ = 2, *K* = 100, *m*_*S*_ = 1, *m*_*C*_ = 4, *m*_*I*_ = 4, *m*_*D*_ = 4, *β*_*S*_ = 4, *β*_*C*_ = 6, *χ*_*I*_ = 3, *u* = 1.5, *δ* = 2; and (F) *r*_*C*_ = 0.5, *r*_*I*_ = 0.5, *r*_*D*_ = 0.5, *K* = 100, *m*_*S*_ = 1, *m*_*C*_ = 3, *m*_*I*_ = 3, *m*_*D*_ = 4, *β*_*S*_ = 3, *β*_*C*_ = 6, *χ*_*I*_ = 3, *u* = 1, *δ* = 511.

**Figure S8:**
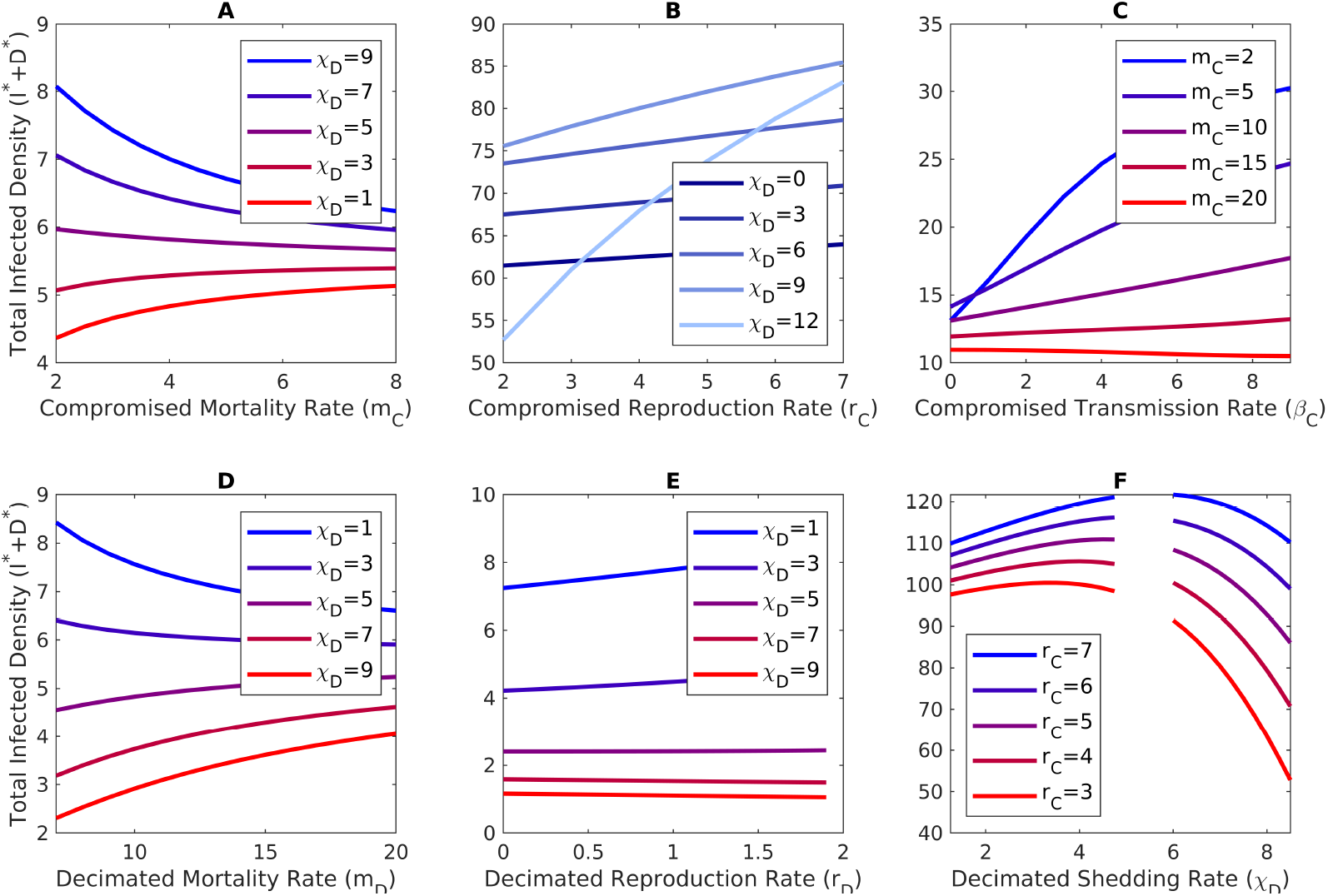
Responses of total infected density to the effects of transgenerational virulence. In each panel, bluer curves illustrate the responses expected in most systems and redder curves illustrate responses of the opposite sign. (A) Total infected density decreases with increased mortality of compromised individuals (bluer curves), unless decimated individuals have sufficiently high shedding rates (redder curves). (B) Total infected density increases with increased reproduction of compromised individuals (all curves). In principle the opposite response can occur (see Table S3 for conditions), but we did not find any numerical examples. (C) Total infected density increases with increase transmission rate of compromised individuals (bluer curves), unless compromised individuals have sufficiently high mortality rates. Total infected density (D) decreases with increased mortality of decimated individuals (bluer curves) and (E) increases with increased reproduction of decimated individuals (bluer curves), unless (D,E) decimated individuals have sufficiently high shedding rates (redder curves). (F) Total infected density increases with increased shedding rates of decimated individuals (blue curves on left side), unless compromised individuals have sufficiently low reproduction rates (redder curves on left side) or infected individuals have sufficiently high shedding rates (all curves right side); the gap is due to the equilibria being unstable. Parameter values are (A) *r*_*S*_ = 6, *r*_*C*_ = 6, *r*_*I*_ = 6, *r*_*D*_ = 6, *K* = 100, *m*_*S*_ = 2, *m*_*I*_ = 10, *m*_*D*_ = 10, *β*_*S*_ = 2, *β*_*C*_ = 2, *χ*_*I*_ = 6, *u* = 1, *δ* = 2; (B) *r*_*S*_ = 7, *r*_*I*_ = 4, *r*_*D*_ = 4, *K* = 1000, *m*_*S*_ = 2, *m*_*C*_ = 3, *m*_*I*_ = 10, *m*_*D*_ = 10, *β*_*S*_ = 2, *β*_*C*_ = 2, *χ*_*I*_ = 6, *u* = 1, *δ* = 2; (C) *r*_*S*_ = *r*_*C*_ = *r*_*I*_ = *r*_*D*_ = 6, *K* = 100, *m*_*S*_ = 2, *m*_*I*_ = 4, *m*_*D*_ = 4, *β*_*S*_ = 4, *χ*_*I*_ = 3, *χ*_*D*_ = 3, *u* = 1.5, *δ* = 1.5; (D) *r*_*S*_ = 4, *r*_*C*_ = 4, *r*_*I*_ = 2, *r*_*D*_ = 2, *K* = 100, *m*_*S*_ = 1, *m*_*C*_ = 4, *m*_*I*_ = 2, *β*_*S*_ = 4, *β*_*C*_ = 4, *χ*_*I*_ = 3, *u* = 1.5, *δ* = 1.5; (E) *r*_*S*_ = 4, *r*_*C*_ = 4, *r*_*I*_ = 2, *K* = 100, *m*_*S*_ = 1, *m*_*C*_ = 4, *m*_*I*_ = 4, *m*_*D*_ = 4, *β*_*S*_ = 4, *β*_*C*_ = 6, *χ*_*I*_ = 3, *u* = 1.5, *δ* = 2; and (F) *r*_*S*_ = 7, *r*_*I*_ = 7, *r*_*D*_ = 2, *K* = 1000, *m*_*S*_ = 2, *m*_*C*_ = 3, *m*_*I*_ = 8, *m*_*D*_ = 8, *β*_*S*_ = 2, *β*_*C*_ = 2, *χ*_*I*_ = 6, *u* = 1, *δ* = 2.

### Section S3 Application of theory to *D. dentifera*-microsporidia system

Here, we use the theory in Sections Section S2.3-Section S2.4 to predict how transgenerational virulence affects the densities of *D. dentifera* and levels of infection by the microsporidian parasite. The first section explains how we roughly estimated the model parameter values. The second section presents the predictions.

#### Section S3.1 Estimates for model parameters

We estimated parameter values for the model using values from our prior studies on *D. dentifera* (Searle et al., 2016) and empirical data collected for this study. Note the following. First, because we did not measure many of the parameter values directly and we did not perform mesocosm experiments, our goal was to get rough estimates for each parameter value. Importantly, while the estimates for the parameter values may be inaccurate, our predictions remain qualitatively unchanged (see next section for details). Second, because our prior work (Searle et al., 2016) focused on a different pathogen, we only used that study to estimate parameter values associated with susceptible individuals (e.g., uptake rates). Third, our parameterized model assumes Lotka-Volterra competition. Because we did not measure filtering rates for susceptible, infected, and compromised individuals, we assume all individuals have equal competitive ability. Thus, the per capita reproduction rate for class *i* is *f*_*i*_ = *r*_*i*_(1 − *N/K*) where *K* is the carrying capacity. Fourth, we do not have data to estimate the carrying capacity (*K*) or the spore degradation rate (*δ*). To account for this, we picked reasonable values and then used simulations to explore how our predictions depend on the values of *K* and *δ*.

##### Maximum reproduction rates (*r*_*S*_, *r*_*I*_, *r*_*C*_, *r*_*D*_)

We estimated *r*_*S*_ =0.3/day based on the rounded value of 0.291/day taken from Searle et al. (2016). The reproduction rate for infected individuals (*r*_*I*_) was set to the same value because measurements of clutches sizes showed that infection did not affect clutch size or the rate at which clutches were produced. The reproduction rates for compromised and decimated individuals were set to *r*_*C*_ = *r*_*D*_ = 0.3 ⋗ 0.4 = 0.12 because measurements of clutch sizes showed that transgenerational virulence reduced reproduction rates by approximately 60%.

Carrying Capacity (*K*): We used a value of *K* = 100 indiv./L, which is the rounded value of 97.5 indiv./L taken from Searle et al. (2016).

##### Non-disease mortality rate (*m*_*S*_)

We estimated *m*_*S*_ = 1*/*30/day. Our empirical observations suggested that susceptible individuals lived 30-60 days at the temperatures used in the experiments. We used the lower value as a conservative estimate.

##### Compromised mortality rate (*m*_*C*_)

We estimated *m*_*C*_ = 5.4*m*_*S*_, i.e., the mortality rate for compromised individuals is 5.66 times larger than the mortality rate for susceptible individuals. Our justification for this estimate is the following. The survival analysis for the standard (‘S’) clone showed that after 20 days survival probability is approximately 80% for susceptible individuals and about 30% for compromised individuals. We modeled mortality as *dS/dt* = −*m*_*S*_*S* and *dC/dt* = *m*_*C*_*C* where *S* is susceptible density and *C* is compromised density. Solving the differential equations for *m*_*S*_ and *m*_*C*_ yields the ratio *m*_*C*_*/m*_*S*_ = *ln*(*C*_*t*_*/C*_0_)*/ln*(*S*_*t*_*/S*_0_) where *S*_0_ and *C*_0_ are initial densities and *S*_*t*_ and *C*_*t*_ are the densities at the end of the 20-day experiment. Substituting *S*_*t*_*/S*_0_ = 0.9 and *C*_*t*_*/C*_0_ = 0.55 yields *m*_*C*_*/m* = ln(0.3)*/* ln(0.8) = 5.4.

##### Infected and decimated mortality rates (*m*_*I*_, *m*_*D*_)

We estimated *m*_*I*_ = *m*_*S*_ = 1*/*30/day and *m*_*D*_ = *m*_*C*_ = 5.4*m*_*S*_ because our empirical observations showed that infection did not significantly affect host lifespan.

Uptake rate (*u*): We estimated *u* = 0.0348 L/day/indv. based on the filtering rate in Searle et al. (2016).

##### Shedding rates (*χ*_*I*_, *χ*_*D*_)

We estimated *χ*_*I*_ = 666 spores/day/indiv. Our observations suggested that infected individuals have around 40 clusters at a time where each cluster has around 50 spores. We estimated that each cluster released spores every 3 days. This means the daily shedding rate is *χ*_*I*_ = 40⋗ 50*/*3 = 666 spores/day. We estimated *χ*_*D*_ = *χ*_*I*_ because our observation suggest that spore production was not affected by material exposure.

Per spore probability of infection (*p*): We estimated *p* = 0.000201. Our justification is the following. Individuals in 10mL (=0.01L) of water were exposed to one ground infected host for 48 hours (=2 days). One ground host yields approximately 666 spores (see shedding rate). The decrease in spore density during exposure was modeled as *dP/dt* = −*uSP* where *P* is spore density, *S* = 1 is the number of susceptible individuals in each beaker, and *u* is the host uptake rate. The initial spore density is *P* (0) = 666*/*0.01 where 0.01L is the volume of the beaker. Solving the ODE and subtracting from the initial spore densities yields that the number of spores encountered and consumed by the individual host is *E* = 666(1 −exp(− 2*u*))*/*0.01 = 4478 spores/L. In the experiments, 90% of individuals became infected, which yields a spore probability of infection of 0.9*/E* = 0.000201.

##### Transmission rates (*β*_*S*_, *β*_*C*_)

Transmission only occurs when individuals ingest spores. Thus, the transmission rate is the product of the uptake rate and the per spore probability of infection, *β*_*S*_ = *pu*. The transmission rates for susceptible and compromised individuals are assumed to be equal (*β*_*C*_ = *β*_*S*_) because the exposure experiments showed that their transmission rates did not differ in the exposure experiments.

##### Spore degradation rate (*δ*)

We used a value of 10/day.

#### Section S3.2 Predicted effects of transgenerational virulence on host densities and infection levels

The sensitivity formulas predict lower total density, lower infection prevalence, and lower infected density at equilibrium because compromised individuals have higher mortality rates (*m*_*C*_ > *m*_*S*_), lower reproduction rates (*r*_*C*_ < *r*_*S*_ implies *f*_*C*_ < *f*_*S*_), and equal transmission rates (*β*_*C*_ = *β*_*S*_) when compared to susceptible individuals and decimated individuals have higher mortality rates (*m*_*D*_ > *m*_*I*_), lower reproduction rates (*r*_*D*_ < *r*_*I*_ implies *f*_*D*_ < *f*_*I*_), and equal shedding rates (*χ*_*D*_ = *χ*_*I*_) when compared to infected individuals. These predictions match the effects of increased mortality and decreased reproduction predicted for most systems; see the summary in Section Section S2.4.4. Note that we do not need to worry about special cases where the predictions are reversed (see Section Section S2.4.4, because in our system (a) compromised individuals do not have much lower production rates than susceptible individuals (i.e., *r*_*C*_ is smaller than *r*_*S*_ for our systems, but not much smaller), (b) decimated and infected individuals have equal shedding rates (*χ*_*D*_ = *χ*_*I*_), and (c) compromised and infected individuals have equal shedding rates (*β*_*C*_ = *β*_*S*_).

Simulations of the parameterized model are shown in Figure 1. The solid curves show the parameterized model dynamics when compromised and decimated individuals have greater mortality rates than susceptible and infected individuals, respectively (*m*_*C*_ > *m*_*S*_ and *m*_*D*_ > *m*_*I*_; solid curves). In this simulation, there is transgenerational virulence because parental infection status affects host mortality rates. The dot-dashed curves show the parameterized model dynamics when compromised and decimated individuals have mortality rates equal to susceptible and infected individuals, respectively (*m*_*C*_ = *m*_*S*_ and *m*_*D*_ = *m*_*I*_; dashed curves). In this simulation, there is no transgenerational virulence because parental infection status does not affect host mortality rates. Note that to facilitate the comparison of the two cases, we keep track of compromised and decimated individuals (i.e., susceptible and infected individuals whose parent was infected) in the simulation without transgenerational virulence even though parental infection status has no effect on host mortality rates.

For the estimated parameters, the reductions in total host density, infection prevalence, and total infected density are 30.3%, 21.7%, and 45.5%, respectively; see the middle column in Table S4. Because we did not have data to estimate the carrying capacity (*K*) and the spore degradation rate (*δ*), we used numerical simulations to explore how variation in those parameters affected the magnitudes of the effects. The percent reductions in total host density, infection prevalence, and infected density are shown in Tables S4 and S5, where the reductions for the default parameters values are given in the middle column. The parameter values are asymmetrically distributed around the default values because sufficiently low carrying capacities (*K*) or sufficiently high degradation rates (*δ*) cause the basic reproduction number (ℛ _0_) to be less than 1.

There is a monotonic relationship between the host carrying capacity (*K*) and the effects of transgenerational virulence on the endemic equilibrium values (Table S4). Specifically, larger values of *K* yield the larger percent decreases in total host density, smaller percent decreases in infection prevalence, and larger percent decreases in infected density. There is also a monotonic relationship between the spore degradation rate (*δ*) and the effects of transgenerational virulence on the endemic equilibrium values (Table S5). In particular, larger values of *δ* yield the smaller percent decreases in total host density, larger percent decreases in infection prevalence, and smaller percent decreases in infected density. We note that for all values of *K* and *δ* we considered, the reductions in all variables were substantial, ranging between between 8% and 74%. In addition, for all values of *K* and *δ* we considered, transgenerational virulence reduced total host density, infection prevalence, and total infected density at all points in time (e.g., solid curves below dashed curves in Figures S9CD).

**Table S4:** Variation in the host carrying capacity (*K*) alters the magnitudes of the effects of TGV on endemic equilibrium values. All parameters except *K* are set to their default values.

| $K$ | 25 | 50 | 100 | 1000 | 5000 |
| --- | --- | --- | --- | --- | --- |
| Total density reduction | 12.1% | 21% | 30.3% | 57.9% | 68.7% |
| Prevalence reduction | 24.2% | 23.9% | 21.7% | 12.1% | 8.9% |
| Infected density reduction | 33.3% | 39.9% | 45.4% | 64% | 71.5% |

**Table S5:** Variation in the spore degradation rate (*δ*) alters the magnitudes of the effects of TGV on endemic equilibrium values. All parameters except *δ* are set to their default values.

| $\delta$ | 0.1 | 1 | 10 | 20 | 50 |
| --- | --- | --- | --- | --- | --- |
| Total density reduction | 71.1% | 57.9% | 30.3% | 21% | 9.3% |
| Prevalence reduction | 8.3% | 12.1% | 21.7% | 23.9% | 23.9% |
| Infected density reduction | 73.5% | 63% | 45.4% | 40% | 31% |

In total, our simulations predict that transgenerational virulence could substantially decrease total density, infection prevalence, and infected density of *D. dentifera*.

**Figure S9:**
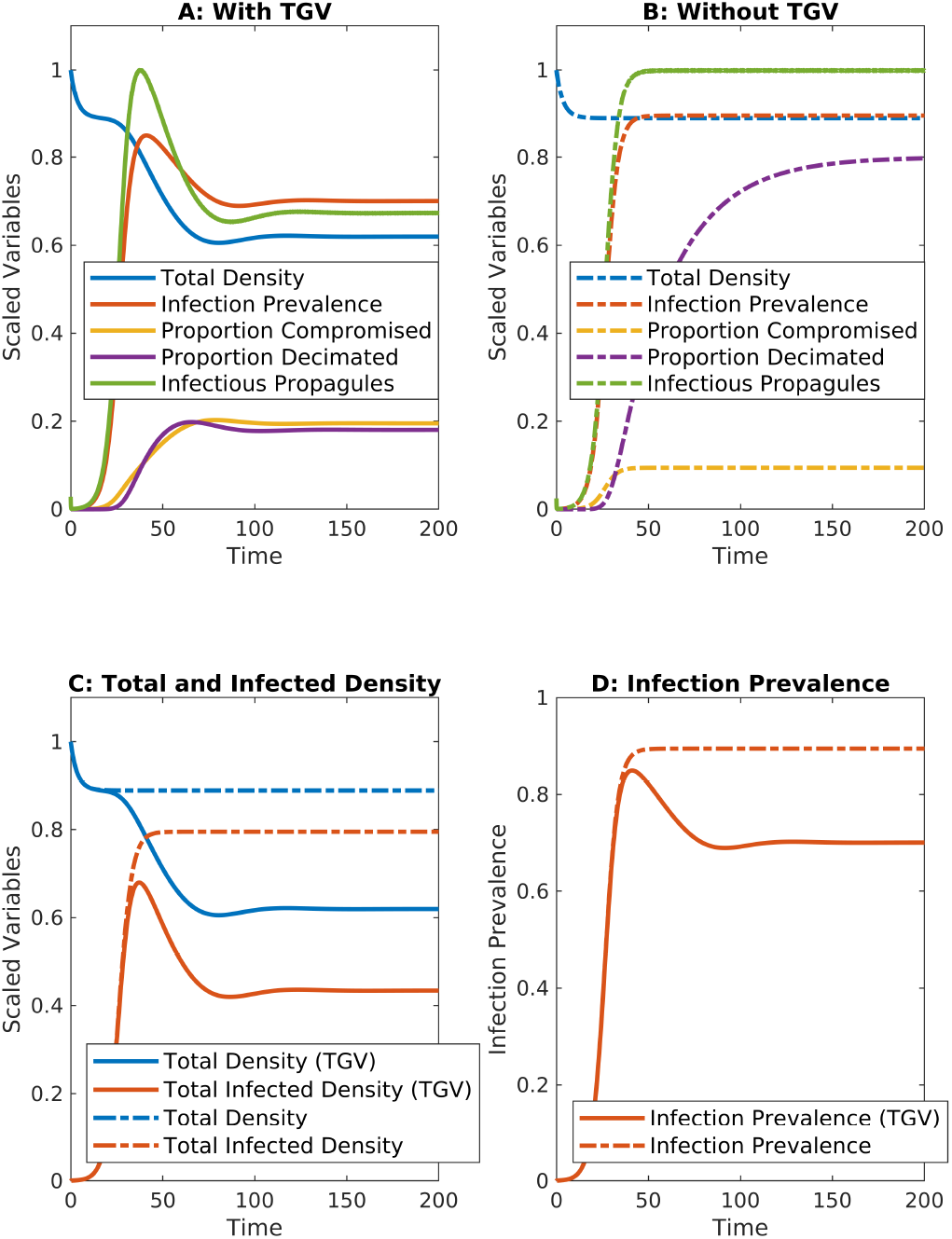
Predicted effects of transgenerational virulence (TGV) on the dynamics of the *D. dentifera*-microsporidian system. The “with transgenerational virulence” (with TGV) simulation is from the parameterized SCIDP model (S86) where compromised and decimated individuals have greater mortality rates and lower reproduction rates than susceptible and infected individuals, respectively (*m*_*C*_ > *m*_*S*_, *r*_*C*_ < *r*_*S*_, *m*_*D*_ > *m*_*I*_, *r*_*D*_ < *r*_*I*_; solid curves). The “without transgenerational virulence” (without TGV) simulation is from the parameterized SCIDP model (S86) where compromised and decimated individuals have mortality rates equal to susceptible and infected individuals, respectively (*m*_*C*_ = *m*_*S*_, *r*_*C*_ = *r*_*S*_, *m*_*D*_ = *m*_*I*_, and *r*_*D*_ = *r*_*I*_; dashed curves). To facilitate the comparison of the two cases, we keep track of compromised and decimated individuals in the simulation without transgenerational virulence even though parental infection status has no effect on host mortality rates. In all panels, total host density and infectious propagule density have been scaled by their maximum value. (A,B) Predicted dynamics for all host classes and infectious propagule density. The time series for total host density and infection prevalence are identical to those in Figure 2 of the main text. (C,D) Direct comparisons of time series showing that at all points in time transgenerational virulence reduces (C) total and infected host density and (D) infection prevalence.

